# A patterned human heart tube organoid model generated by pluripotent stem cell self-assembly

**DOI:** 10.1101/2022.12.16.519611

**Authors:** Brett Volmert, Ashlin Riggs, Fei Wang, Aniwat Juhong, Artem Kiselev, Aleksandra Kostina, Colin O’Hern, Priyadharshni Muniyandi, Aaron Wasserman, Amanda Huang, Yonatan Lewis-Israeli, Sangbum Park, Zhen Qiu, Chao Zhou, Aitor Aguirre

**Affiliations:** Institute for Quantitative Health Science and Engineering, Division of Developmental and Stem Cell Biology, Michigan State University, MI, USA; Department of Biomedical Engineering, College of Engineering, Michigan State University, MI, USA; Department of Biomedical Engineering, Washington University in Saint Louis, MO, USA; Institute for Quantitative Health Science and Engineering, Division of Biomedical Devices, Michigan State University, MI, USA; Department of Electrical and Computer Engineering, College of Engineering, Michigan State University, MI, USA; Department of Pharmacology and Toxicology, College of Human Medicine, Michigan State University, MI, USA; Division of Dermatology, Department of Medicine, College of Human Medicine, Michigan State University, East Lansing, MI 48824, USA

## Abstract

Human pluripotent stem cells can recapitulate significant features of mammalian organ development *in vitro*, including key aspects of heart development. We hypothesized that the organoids thus created can be made substantially more relevant by mimicking aspects of *in utero* gestation, leading to higher physiological and anatomical resemblance to their *in vivo* counterparts. Here, we report steps towards generating developmentally inspired maturation methodologies to differentiate early human heart organoids into patterned heart-tube-like structures in a reproducible and high-throughput fashion by complete self-organization. The maturation strategy consists of the controlled and stepwise exposure to metabolic (glucose, fatty acids) and hormonal signals (T3, IGF-1) as present during early heart development. These conditions elicit important transcriptomic, cellular, morphological, metabolomic, and functional changes over a 10-day period consistent with continuously increasing heart complexity, maturation, and patterning. Our data reveals the emergence of atrial and ventricular cardiomyocyte populations, valvular cells, epicardial cells, proepicardial-derived cells, endothelial cells, stromal cells, conductance cells, and cardiac progenitors, all of them cell types present in the primitive heart tube. Anatomically, the organoids elongate and develop well-differentiated atrial and ventricular chambers with compacted myocardial muscle walls and a proepicardial organ. For the first time in a completely self-organizing heart organoid, we show anterior-posterior patterning due to an endogenous retinoic acid gradient originating at the atrial pole, where proepicardial and atrial populations reside, mimicking the developmental process present within the primitive heart tube. Collectively, these findings highlight the ability of self-organization and developmental maturation strategies to recapitulate human heart development. Our patterned human heart tube model constitutes a powerful *in vitro* tool for dissecting the role of different cell types and genes in human heart development, as well as disease modeling congenital heart defects, and represents a step forward in creating fully synthetic human hearts.

## Introduction

Cardiovascular diseases (CVDs), including disorders of the heart and blood vessels, are the leading causes of death globally, contributing to an estimated 17.9 million deaths annually^1^. Laboratory models of the heart are used to better understand the etiology and mechanisms of CVDs in high detail. Several model systems to research CVDs are used, ranging from primary and induced pluripotent stem cell (iPSC)-derived cardiomyocyte cultures to animal models and 3D-culture systems such as spheroids and engineered heart tissues^2–7^. Nevertheless, many of these systems fail to fully recapitulate aspects of the complex nature of the human heart due to a variety of reasons, including the absence of endogenous extracellular matrix (ECM) and non-cardiomyocyte cardiac cell types, as well as the lack of physiological morphology and cellular organization^8, 9^. In addition, animal models possess distinct, non-human physiology, metabolism, electrophysiology and pharmacokinetic profiles which often do not predict human-relevant responses^8, 9^ accurately.

The introduction of human-relevant models is paramount to the discovery of effective, clinically translatable solutions to CVDs. Over the last ten years, innovations in human induced pluripotent stem cell (hiPSC)^10–12^ and organoid^13, 14^ technologies have advanced techniques to better model and study human systems with increasing precision. Recently, we and others have described methodologies to create human heart organoids from pluripotent stem cells. These methods enable the study of human heart development and disease^15–19^ in a dish to a degree unseen before due to their cellular complexity and physiological relevance. Yet, these systems still fall short of recapitulating important aspects of human heart development and the late embryonic human heart, such as anterior-posterior patterning, coronary vascularization and lack important cell populations contributing to heart structure (e.g. neural crest). There is a pressing need to develop more sophisticated *in vitro* heart organoid model systems for better understanding human heart development and disease pathology.

Here, we report an advanced set of developmentally inspired conditions to induce further developmentally relevant cellular, biochemical, and structural changes in human heart organoids in a high throughput setting by complete self-assembly, bringing heart organoids one step closer to 4–6-week-old gestational hearts. The methods introduced here take inspiration from developmental and maturation timeline paradigms which elicit distinct intra-organoid cellular compositions, transcriptomes, functionalities, and metabolic profiles generated in part through a retinoic acid morphogen gradient present only in our most advanced developmental maturation strategy. Our method is a simple, streamlined, low-cost, and high-fidelity approach compared to similar *in vitro* heart tissue culture techniques. The platform is also highly automatable, scalable, adaptable, and overall amenable to high throughput screening approaches for the investigation of human heart development, cardiac diseases, toxicity testing and pharmacological discovery.

## Results

### Extended heart development modeling through improved developmental induction strategies

We recently detailed a protocol for the generation of self-organizing early embryonic human heart organoids which constitutes the starting step for the methodology described below^15^. Heart organoids were differentiated from hiPSC embryoid bodies to the cardiac lineage between days 0 and 7 through a timewise 3-step Wnt pathway modulation strategy, and then cultured until day 20 in RPMI^15^. To examine the effect of more advanced organoid culture strategies mimicking *in utero* conditions on heart organoid development, we took day 20 early embryonic-like heart organoids and employed four different developmental induction strategies from day 20 to day 30 **(Fig. 1a)**. These strategies are partly based on previous human and animal developmental studies^20–24^ and represent gradual increasing steps in complexity relevant to *in utero* conditions (in order of less complex to more complex: control, maturation medium, enhanced maturation medium 1, enhanced maturation medium 2/1). Our Control strategy represents a continuation of organoid culture in the base medium used for organoid formation, RPMI/B27. The maturation medium (MM) strategy uses RPMI/B27 with added fatty acids (an embryonic relevant concentration of oleic acid, linoleic acid, and palmitic acid)^23, 24^ and L-carnitine^25^ to facilitate a developmentally relevant transition from glucose utilization to fatty acid metabolism characteristic of the fetal human heart^26–30^. The MM strategy also uses T3 hormone, a potent activator of organ growth during embryonic development and metabolic maturation and has been shown to stimulate cardiovascular growth^31, 32^. The enhanced maturation medium 1 (EMM1) strategy uses the same basal composition as MM but decreases the concentration of glucose to cardiac physiological levels^33–35^ (from 11.1 mM to 4 mM to further encourage the transition to fatty acid oxidation) and adds ascorbic acid as a reactive oxygen species scavenger to counteract the increased oxidative stress^36, 37^. Enhanced maturation medium strategy 2/1 (EMM2/1) utilizes a combination of two different media formulations. During days 20-26, EMM2 media is utilized and is the same basal composition as EMM1 and contains added IGF-1. IGF-1 plays important roles during embryonic and fetal development in tissue growth and maturation, especially in the heart, as proven in murine and human studies^38–40^. From day 26 onwards, EMM1 media is utilized in the EMM2/1 strategy. EMM2/1 constituted our most advanced condition and mimicked *in utero* heart development to the greatest extent. More detailed descriptions of all developmental induction strategies along with concentrations of respective media formulations can be found in the Materials and Methods section.

**Fig. 1.**
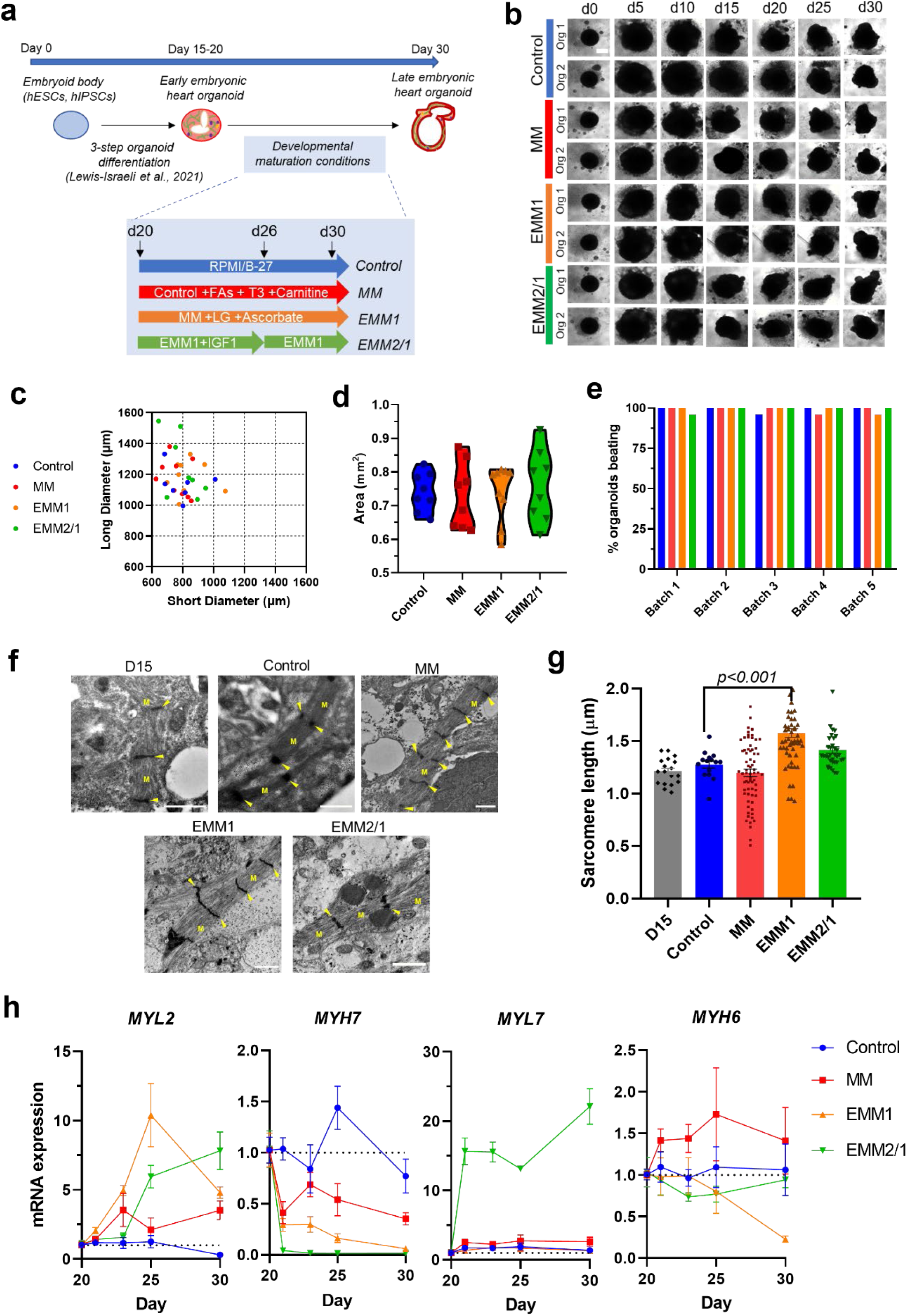
Developmental induction methods for improving human heart organoid developmental modeling. **a,** A schematic diagram depicting the differentiation protocol for creating human heart organoids and the media conditions for the four maturation strategies (Control, MM, EMM1, and EMM2/1). **b,** Brightfield images of organoids throughout the 30-day culture period. Two representative organoids are shown for each condition (data representative of 23-24 organoids per condition). Scale bar = 400 μm. **c,** Quantification of organoid long diameter and short diameter at day 30 of culture for each maturation strategy (n=7-8 organoids per condition). **d,** Quantification of organoid area at day 30 of organoid culture for each maturation strategy (n=7-8 organoids per condition). Data presented as a violin plot with all points. **e,** Quantification of percentage of organoids visibly beating under brightfield microscopy in each condition from 5 different organoid batches (n=22-24 organoids in each condition for each batch). **f,** TEM images displaying sarcomeres, myofibrils (M) and I-bands (arrows) in day 15 organoids and in organoids from each maturation condition at day 30 (n=4 organoids per condition). Scale bars = 1 μm. **g,** Quantification of sarcomere length within TEM images. Data presented as mean ± s.e.m (n=4 organoids per condition). One-way ANOVA with Brown-Forsythe and Welch multiple comparisons tests. **h,** mRNA expression of key sarcomeric genes involved in cardiomyocyte maturation between days 20 and 30 of culture for each condition (n=4 organoids). Values = mean ± s.e.m.

Heart organoids treated with the different developmental induction strategies continued to grow and develop, with drastic changes in morphology depending on condition **(Fig. 1b-d, Extended Data Fig. 1 a, b)**. Organoids experienced a period of rapid growth from day 0 to day 10, increasing in diameter while retaining their spherical structure **(Fig. 1b)** and continuing to grow until day 30. Organoids developed distinct elliptical morphologies after day 20, elongating and contorting as observed by brightfield microscopy and growing to possess long diameters between 1000 and 1600 μm, while short diameters ranged from 600 to ∼1000 μm on day 30 **(Fig. 1b, c)**. Organoid area measured by brightfield microscopy revealed similar trends for each condition, between 0.6 mm^2^ to ∼0.9 mm^2^ **(Fig. 1d)**. Measured over time, organoid area rose sharply over the initial 10 days of culture to a peak of 1.38 mm^2^ and declined steadily until day 30 where organoids possessed areas of 0.74 ± 0.0072 mm^2^, 0.73 ± 0.012 mm^2^, 0.74 ± 0.0093 mm^2^ and 0.76 ± 0.013 mm^2^ for Control, MM, EMM1, and EMM2/1 organoids, respectively **(Extended Data Fig. 1a)**. Moreover, organoid circularity rapidly declined from day 0 to day 10, then remained steady until day 25 where it experienced a slight decline until day 30 **(Extended Data Fig. 1b)**. By day 30 of culture, nearly 100% of organoids in every condition were observed to be beating over five different batches of organoids (n=22-24 organoids per condition per batch) **(Fig. 1e, Supplementary Videos 1-4)**. Transmission electron microscopy (TEM) images indicated the presence of well-developed myofibrils and the formation of sarcomeres within the organoids **(Fig. 1f)** in all conditions, with sarcomeres in the EMM1 condition displaying a significantly increased sarcomere length of 1.58 ± 0.323 μm relative to control **(Fig. 1g)**. qRT-PCR revealed the differential expression of hallmark sarcomeric cardiomyocyte genes from day 20 to day 30 as expected, indicating cardiomyocyte function remained fundamentally unaffected throughout different developmental induction conditions at a general level **(Fig. 1h)**.

### scRNA-seq of human heart organoids under developmental induction reveals differences in cellular composition

To characterize the cellular and transcriptomic composition of heart organoids in each of our developmental induction conditions, we performed scRNA-seq at day 34 of organoid culture. UMAP projections display unsupervised K-means clustering analyses **(Fig. 2a)**. Ventricular and atrial cardiomyocytes (VCMs and ACMs, respectively), valve cells (VCs), proepicardial derived cells (PEDCs), stromal cells (SCs), cardiac progenitor cells (CPCs), conductance cells (CCs), and endothelial cells (ECs) were revealed in the control condition. MM, EMM1, and EMM2/1 exhibited generally the same cardiac cell lineage populations, however the abundance of several significant groups varied according to the developmental medium conditions. Additionally, these strategies elicited the emergence of a distinct epicardial cell (EPC) cluster. Control organoids were composed of 17% VCMs, 16% ACMs, 9% VCs, 17% PEDCs, 26% SCs, 9% CPCs, 5% CCs, and 1% ECs **(Fig. 2b)**. Relative to control, MM organoids displayed an increased percentage of both VCMs and ACMs (25% and 34%, respectively), decreased VCs (4%), decreased PEDCs (12%), 1% EPCs, decreased SCs (15%), 9% CPCs, and decreased CCs (1%). Relative to control, EMM1 organoids contained increased VCMs (38%), decreased ACMs (9%), decreased VCs (4%), decreased PEDCs (16%), 4% EPCs, decreased SCs (18%), increased CPCs (11%), and decreased CCs (1%). Relative to control, EMM2/1 organoids exhibited increased VCMs (28%), decreased ACMs (5%), 9% VCs, decreased PEDCs (14%), 3% EPCs, 26% SCs, increased CPCs (13%), and decreased CCs (1%). Cluster identification was performed through the spatial identification of key genes **(Fig. 2c, Supplementary Figs 1-8).** VCMs possessed high expression of *MYH7* and *MYL3*^41, 42^, while ACMs expressed *MYH6* and *MYL7*^41, 42^. VCs were identified through the expression of *SOX9*, *NPR3*, *CDH11*, *DCN*, and *COL1A1*^42–45^. CCs displayed high expression of *NEUROD1*, *VSNL1*, *TAGLN3*, *SOX1*, *ISL1*, *STMN2*, and *ALDH1A1*^41, 46–50^. PEDCs showed high expression of *WT1*, *TBX18*, *TCF21*, and *ALDH1A2*.^51–53^ EPCs displayed similar markers as PEDCs mentioned previously, yet also expressed *ITLN1*, *MSLN*, *ALOX15*, and *SLPI*^41, 54^. Cardiac fibroblasts were also identified within the EPDC cluster, showing expression of *DCN*, *LUM*, *OGN*, and *POSTN*^41, 55^. SCs were identified by expression of *SOX18*, *TUBB3*, *SOX2*, and *FOXA2*^56–61^. ECs possessed high expression of *PECAM1*, *CDH5*, *ENG*, and *ETS1*^41, 62^. These results suggested, in agreement with previous studies on cardiac development^41, 42^, that by day 20 the main cardiac cell lineages are already determined, but that developmental induction conditions can exert dramatic effects on the expansion and maturation of these cell types to better reflect *in vivo* heart development. Organoids in all conditions also recapitulated key genes involved in left-right asymmetry such as *PITX2*, *PRRX2*, *LEFTY1 and PRRX1*^63–66^ **(Supplementary Fig. 9).** PITX2 was upregulated in the VCM, ACM, PEDC, EPC, and EC clusters. PRRX2 was upregulated in the SC, PEDC, and EC clusters and *LEFTY1* was highly upregulated in the SC cluster. *PLCXD3* was upregulated in VCMs in all conditions, whereas *PRRX1* was upregulated in ACMs in control and MM organoids but upregulated in VCMs for EMM1 and EMM2/1 organoids. Additionally, organoids displayed high upregulation of proliferation markers such as *MKI67*, *PCNA*, *AURKB*, and *CDK1* in the VC clusters in all conditions, indicating that important growth and remodeling are still undergoing at day 34 of differentiation^67–69^ **(Supplementary Fig. 10)**. Cells of the first heart field (FHF) and the anterior and posterior second heart fields (aSHFs and pSHFs, respectively) contribute to linear heart tube expansion and subsequent chamber formation and are important for proper cardiac morphogenesis^70^. Various FHF and SHF heart field markers were observed in organoids in all conditions^71, 72^ **(Supplementary Fig. 11-14)**. *HAND1, HAND2*, *TBX5*, and *HCN4* were all upregulated in the VCM and ACM clusters for all conditions. *ISL1* was upregulated in the VCM and ACM clusters as well as the CC cluster for each condition. In addition, outflow tract markers such as *RSPO3*^73^ and *WNT5A*^74, 75^ were upregulated in the PEDC, ACM, VCM, and SC clusters for all conditions **(Supplementary Fig. 15)**.

**Fig. 2.**
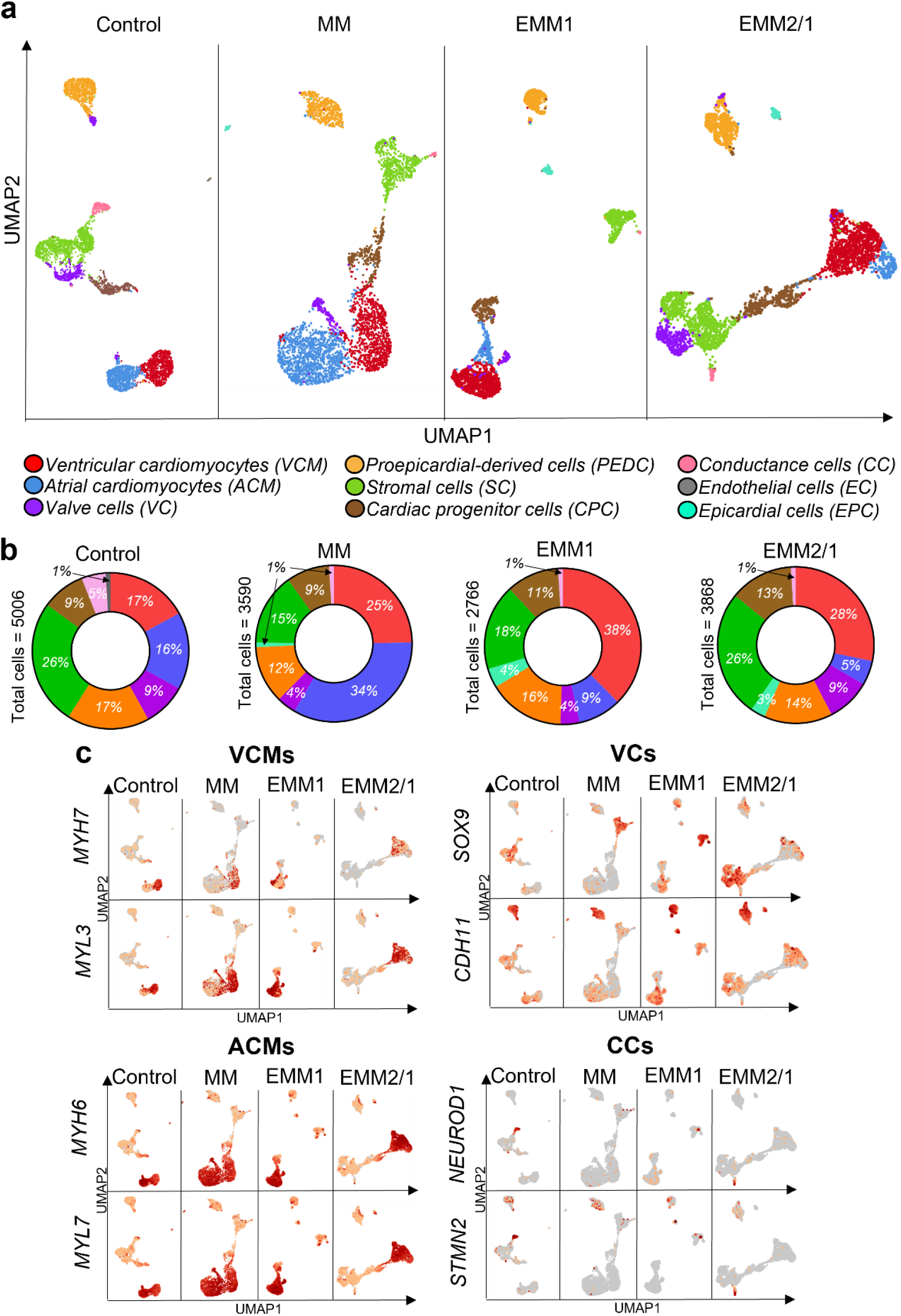
Single-cell RNA sequencing of human heart organoids reveals distinct cardiac cell populations. **a,** UMAP projections of k-means 8 clustering of single-cell RNA sequencing data for each condition in day 34 organoids. Cluster identities are located in the legend below. **b,** Quantification of total cell count percentages per cluster. Colors of regions correspond to those found in the legend in **(a)**. **c,** Differential expression heatmaps for key genes in the ACM, VCM, VC and CC clusters for each developmental maturation condition.

Extending these analyses, we utilized publicly available data from the Human Cell Atlas project^41^ from gestational day 45 (GD45) to compare our conditions to that of the developing human heart **(Extended Data Fig. 2)**. Based on their time in culture, human heart organoids should be closest to GD45 human fetal hearts **(Extended Data Fig. 2a)**. Differential expression of a wide array of genes comprising key cardiac populations and including ventricular and atrial cardiomyocytes, cardiac fibroblasts, epicardial cells, endothelial cells and conductance markers showed that our human heart organoids possessed similar gene expression profiles to GD45 when exposed to developmental induction conditions, especially in the case of EMM2/1, our closest condition mimicking heart development **(Extended Data Fig. 2b)**. In addition, control and Day 15 organoids exhibited the closest similarity to each other and the least similarity to GD45 human hearts, demonstrating that our organoids gradually matured transcriptionally as we employed more advanced maturation strategies.

To complement the above scRNA-seq analyses, dot plots describing the average and percent expression of key lineage-defining differentially expressed genes for individual clusters are depicted for each developmental induction condition illustrating the cellular complexity of our heart organoids at day 34 **(Fig. 3a)**. As we and others have shown before^13–15, 17, 76^, the high cellular complexity of the organoids drives self-organization and cell-cell communication. We performed computational analysis of cell-cell communication networks for key genes found in the organoids. We identified various complex receptor-ligand communication pathways within our human heart organoids in each condition **(Fig. 3b)**. Receptor-ligand networks include JAG1-NOTCH1, PDGFRs, IGF2-IGF2R, INSR, and VEGF, among others. We also performed Gene Ontology (GO) analysis for biological process terms corresponding to top differentially expressed genes contributing to the ontology for each cluster, as well as top shared genes between all four conditions per cluster (**Supplementary Figures 16, 17**). To further investigate cell-cell communication networks, we utilized scRNA-seq data to highlight key receptor-ligand pairs within the organoids from each maturation condition **(Supplementary Fig. 18-21)**. This data highlights the ability and sensitivity of our organoids to respond to various developmental maturation stimuli surrounding cell-cell communication paradigms.

**Fig. 3.**
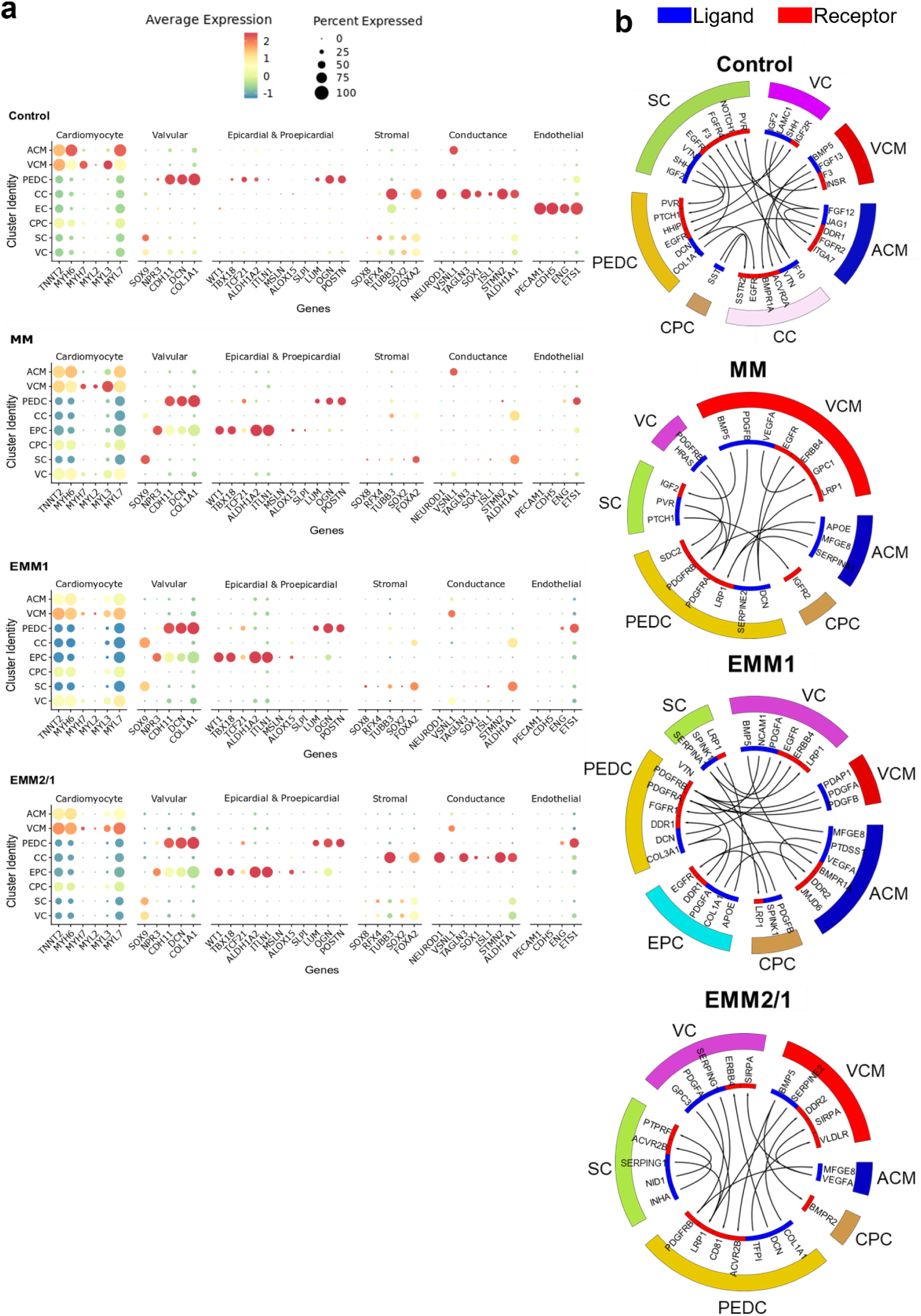
Cluster identity and cell-cell communication networks highlight the importance of self-organization in heart organoid development. **a,** Dot plot of differentially expressed genes in each cluster for each condition. Color is indicative of the average expression level across all cells, and the size of the circle is indicative of the percentage of cells within a particular cluster that express the respective gene. **b,** Visualization of cell-cell ligand-receptor communication networks for each condition. Colors of clusters (exterior) matches that of scRNAseq UMAP projections. Ligands are indicated as blue bands and receptors are indicated by red bands. Arrows within depict pairing from ligands to receptors.

### Mitochondrial maturation and oxidative metabolism in human heart organoids under developmental induction conditions

The early developing human heart relies heavily on glycolysis for energy expenditure. As it continues to grow, it decreases its reliance on glycolysis and switches to fatty acid oxidation for the bulk of energy consumption^28, 30, 77–79^. Therefore, we sought to determine the effect that our developmental induction conditions, and particularly EMM2/1, exerted on mitochondrial growth and metabolic transcriptional activity within heart organoids. Through the addition of MitoTracker, a mitochondrial-permeable fluorescent dye, we visualized live mitochondrial content within heart organoids at day 30 of culture **(Fig. 4a)**. Control organoids displayed few and diffuse mitochondria, while EMM2/1 organoids possessed the most developed mitochondrial content of all conditions (abundance, morphology) (**Fig. 4a, b**). An increasing trend of mitochondrial content in MM, EMM1, and EMM2/1 organoids (fold change of 1.73 ± 0.10, 2.60 ± 0.11, and 3.10 ± 0.18, respectively) was quantified relative to control, suggesting that developmentally maturated organoids had an increasingly higher capacity for aerobic respiration and responded positively to maturation stimuli **(Fig. 4b)**. TEM revealed high-magnification detail on mitochondrial presence within organoids at day 30 of culture **(Fig. 4c)**. Compared to day 15 mitochondrial size, Control organoid mitochondrial size was similar **(Fig. 4d)**. However, mitochondrial size within MM, EMM1 and EMM2/1 organoids dramatically increased relative to that of Control organoids. We employed qRT-PCR at different timepoints from day 20 to day 30 of organoid culture to explore the differential gene expression of two key OXPHOS genes in cardiac metabolic maturation: *CPT1B*, a critical rate-limiting fatty acid transporter element^80, 81^, and *PPARGC1A*, a master regulator of mitochondrial biogenesis^82^ **(Fig. 4e)**. *CPT1B* expression increased 1.5-fold at day 30 in the EMM2/1 condition relative to control, yet expression in MM and EMM1 organoids decreased ∼1-2-fold. *PPARGC1A* levels were up to 3-fold higher in all maturation conditions between day 20 and day 25 relative to control. At day 30, MM and EMM2/1 organoid expression remained elevated relative to control (1.5-fold) and expression had normalized in EMM1 organoids. We then used scRNA-seq gene expression data from the ACM and VCM clusters to look for a wider set of metabolic markers as the organoids developed in the different conditions. We found that organoids in the EMM2/1 condition expressed much higher levels of key metabolic genes, including those involved in mitochondrial biogenesis and function, oxidative phosphorylation, TCA cycle, and glycolysis **(Fig. 4f)**. Furthermore, we performed computational transcriptomic analysis and mapping to KEGG metabolic pathways using Pathview^83, 84^ **(Supplementary Fig. 22-29)**. In agreement with our other metabolic data, EMM2/1 organoids showed reduced activity of glycolytic complexes (**Supplementary Fig. 22-25**) and increased activity of mitochondrial respiratory complexes (**Supplementary Fig. 26-29**), indicative of progressive developmental maturation. Overall, these results suggested that EMM2/1 organoids recapitulate significant aspects of cardiac metabolism *in vitro* reminiscent of fetal cardiac development at similar stages.

**Fig. 4.**
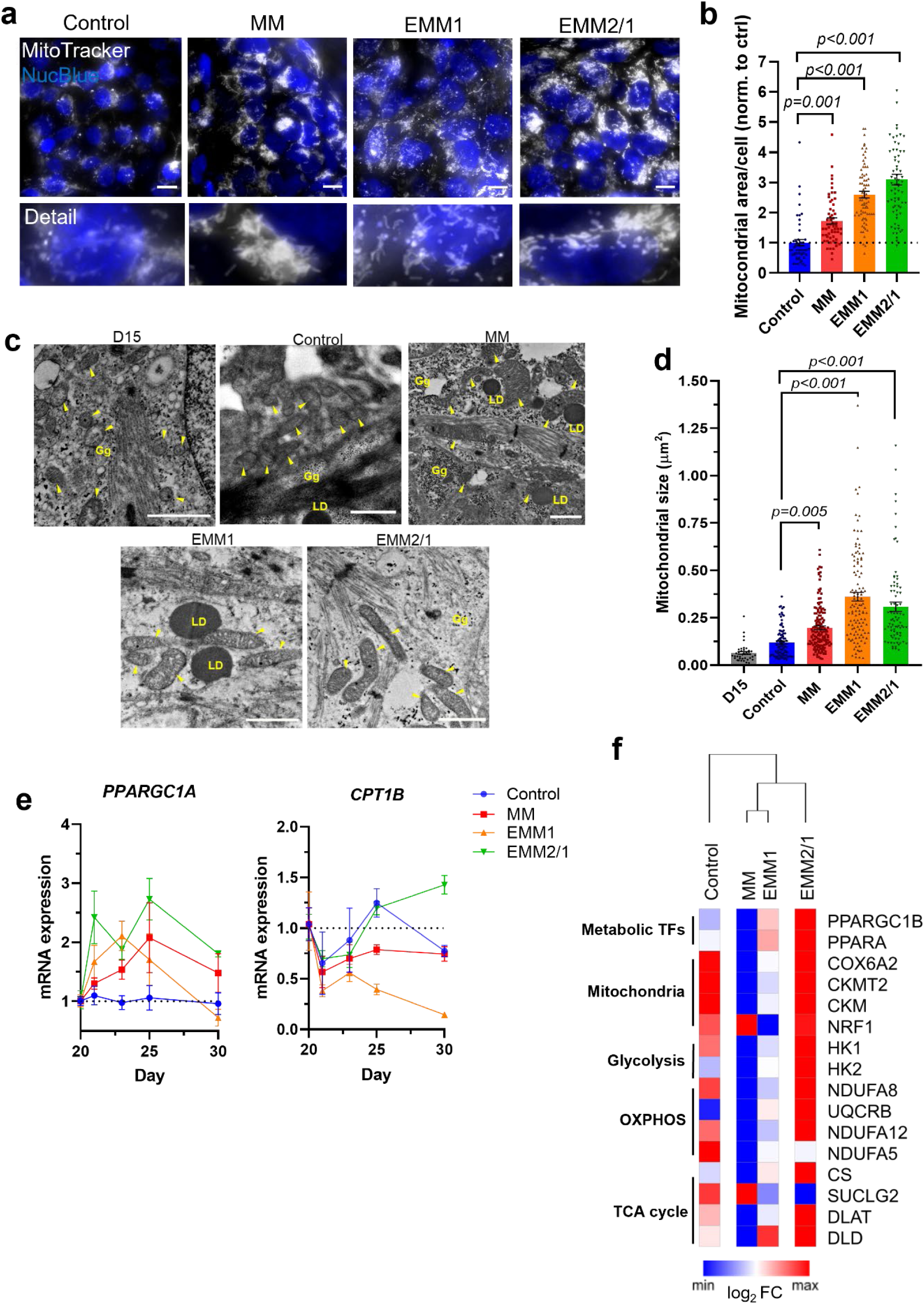
Human heart organoids develop increasingly mature metabolic profiles following developmental induction conditions. **a,** Mitochondrial labeling within day 30 human heart organoids in each condition (n=6 organoids per condition). Mitotracker = white, NucBlue = blue. Scale bars = 10 μm. Detailed images of mitochondria are shown below each main image. **b,** Quantification of mitochondrial area surrounding each individual nucleus. Values = mean ± s.e.m., one-way ANOVA with Brown-Forsythe and Welch multiple comparisons tests. **c,** TEM images displaying mitochondria in day 15 organoids and in organoids from each maturation condition at day 30 (n=4 organoids per condition). Blue arrows indicate mitochondria, LD = lipid droplets, Gg = glycogen granules. Scale bars = 1 μm. **d,** Quantification of mitochondrial area from TEM images. Values = mean ± s.e.m., one-way ANOVA with Brown-Forsythe and Welch multiple comparisons tests. **e,** mRNA expression of metabolic genes *PPARGC1A* and *CPT1B* between days 20 and 30 of culture for each condition (n=4 organoids per condition). Values = mean ± s.e.m. **f,** mRNA expression heatmap of metabolic genes from D34 organoids in each condition. Data shown as log_2_ fold change and normalized to each row.

### Developmental induction conditions promote progressive electrophysiological maturation in human heart organoids

The emergence and presence of the cardiac conduction system, including specific ion channels and membrane receptors, such as those surrounding calcium, potassium, and sodium currents, represent critical elements of the cardiomyocyte action potential^85–87^ and fetal heart development^88, 89^. We sought to characterize the functionality of heart organoids under developmental induction conditions through electrophysiology and immunofluorescence for key markers. We assessed calcium transient activity of individual cardiomyocytes within human heart organoids at day 30 using the membrane-permeable dye Fluo-4 **(Fig. 5a, Supplementary Videos 5-8)**. Organoids in all conditions exhibited distinct and regular calcium transient activities with varying peak amplitude and action potential frequencies **(Fig.5 b, c)**. Control and MM organoids presented smaller peak amplitudes when compared to EMM1 and EMM2/1 organoids, indicating less robust contractions **(Fig. 5b)**, and presented similar beat frequencies ∼1.5 Hz. EMM1 organoids displayed abnormally high beating rates (∼2.5 Hz) for the heart at this stage, while EMM2/1 organoids showed beat frequencies at ∼1-1.5 Hz **(Fig. 5c)**. In general, and except for EMM1 organoids, developmentally induced organoids presented beating rates compatible with what has been described for early human embryos at GD45^90, 91^ (60-80 beats per minute). In addition to calcium, the total electrophysiological response of the cardiomyocyte action potential incorporates additional ion currents such as potassium and sodium. We investigated the voltage activity within EMM2/1 heart organoids via the potentiometric dye di-8-ANEPPS and identified unique actional potentials within individual cardiomyocytes indicative of the presence of specialized ventricular-, atrial-, and nodal-like cells **(Fig. 5d)**. Furthermore, proper excitation-contraction coupling, and depolarization and repolarization of cardiomyocytes depends on specialized invaginations of the sarcolemma (t-tubules), which are indicative of cardiomyocyte maturation^92, 93^. We assessed t-tubule presence via fluorescently labeled wheat germ agglutinin (WGA) staining in human heart organoids at day 30 **(Fig. 5e)**. T-tubules were observed surrounding the nucleus and sarcomeres (TNNT2+) within organoids in each condition, with increasing t-tubule density in the EMM1 and EMM2/1 conditions **(Fig. 5f)**. We also assessed the presence of the potassium ion channel KCNJ2 via confocal microscopy **(Fig. 5g)**. KCNJ2+ puncta were observed in each condition, with a 2-fold increased presence in the EMM2/1 condition **(Fig. 5h)**. To investigate the temporal dynamics of key ion channels through the application of our developmental maturation strategies, we utilized qRT-PCR from day 20 to day 30 of organoid culture to assess levels of calcium (*ATP2A2*), potassium (*KCNJ2*), and sodium (*SCN5A*) transporters **(Extended Data Fig. 3a)**. *ATP2A2* expression increased in all conditions relative to control, with EMM2/1 exhibiting the most marked upregulation of 7-fold. *KCNJ2* expression steadily decreased in the EMM1 condition relative to control, with MM organoids exhibiting upregulation at day 30 and EMM2/1 displaying upregulation throughout the culture period. *SCN5A* expression showed upregulation for the MM condition at day 25 and 30, while EMM2/1 also displayed upregulation at day 30 relative to control. EMM1 expression remained consistent with control until day 25 and was downregulated at day 30.

**Fig. 5.**
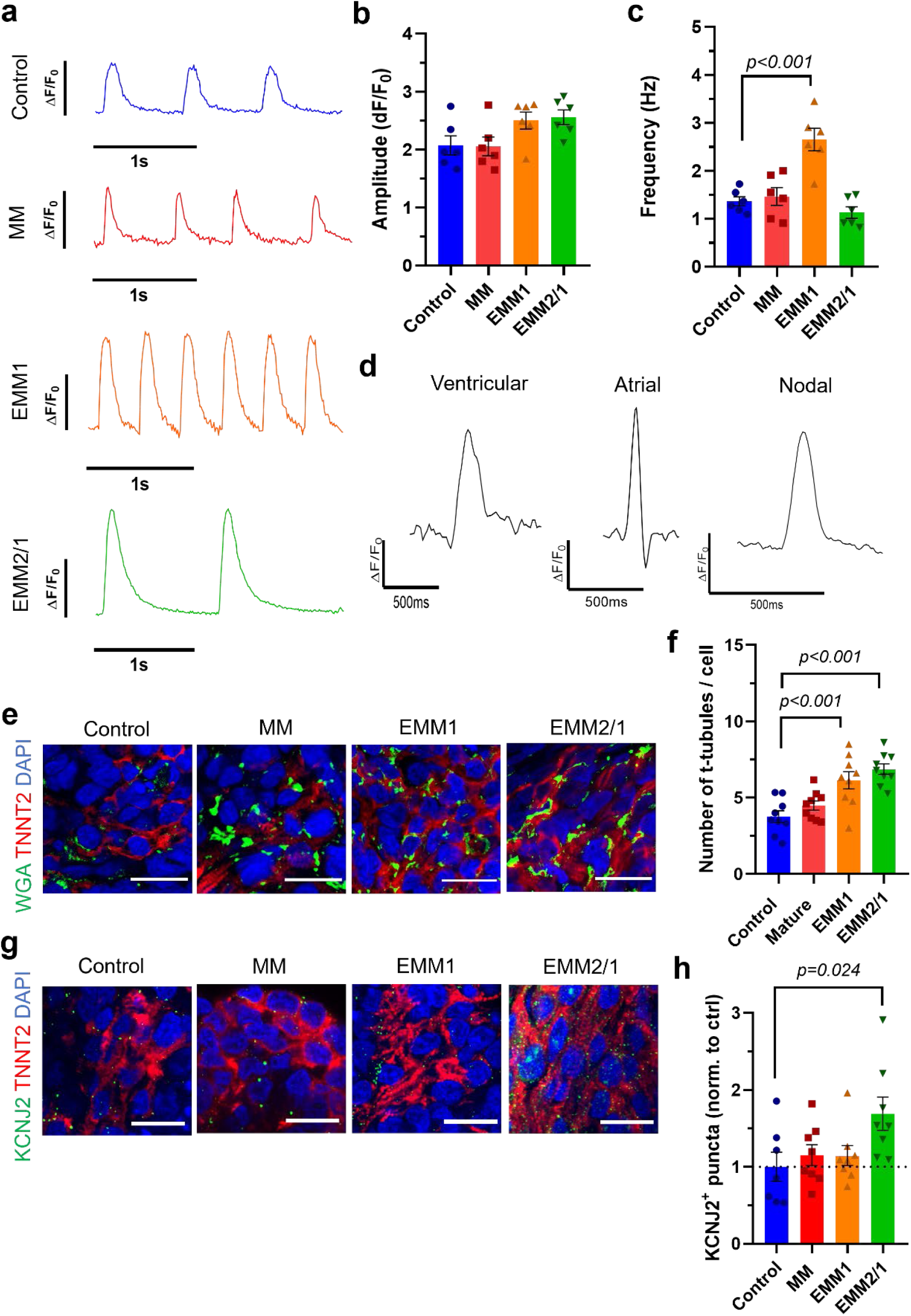
Developmental induction conditions promote progressive electrophysiological maturation in human heart organoids. **a,** Representative calcium transient traces within day 30 human heart organoids from each condition. **b,** Quantification of peak amplitude of calcium transient traces from each condition with and without the addition of isoproterenol. A minimum of 2 regions and 16 peaks were quantified and averaged for each organoid. Values = mean ± s.e.m. **c,** Quantification of calcium transient peak frequency for each condition with and without the addition of isoproterenol. A minimum of 2 regions and 16 peaks were quantified and averaged for each organoid. Values = mean ± s.e.m. **d,** Representative voltage tracings of organoids in the EMM2/1 condition depicting ventricular, atrial, and nodal action potentials. **e,** Immunofluorescence images of WGA staining (t-tubules) within TNNT2+ regions in day 30 organoids for each condition (n=8 organoids per condition). **f,** Quantification of t-tubules surrounding individual nuclei for each condition. Values = mean ± s.e.m., one-way ANOVA with Brown-Forsythe and Welch multiple comparisons tests. **g,** Immunofluorescence images of KCNJ2+ puncta within TNNT2+ regions in organoids for each condition (n=8 organoids per condition). **h,** Quantification of the number of KCNJ2+ puncta for each condition. Data presented as fold change relative to control organoid mean value. Values = mean ± s.e.m., one-way ANOVA with Brown-Forsythe and Welch multiple comparisons tests.

An additional ion channel of critical importance is the hERG channel^94–96^ encoded by the gene *KCNH2*. Mutations and perturbations in this channel can lead to shortening or prolongation of the QT interval^95, 97–99^, and drug interactions with this channel can lead to cardiac arrythmia which represents a critical bottleneck surrounding drug discovery and development^100,101^. We assessed the expression of *KCNH2* within our organoids and discovered high expression within the ACM and VCM clusters in all conditions **(Extended Data Fig. 3b)**. The expression of *KCNH2* was quantified and displayed a decreasing trend in the ACM cluster throughout the maturation conditions **(Extended Data Fig. 3c)** and maximum expression within the VCM cluster in EMM2/1 conditions. Further, autonomic control of the cardiac conduction system through adrenergic signaling plays a large role in physiological functionality^102–104^, and underlies a range of CVDs from heart failure and hypertension to arrythmia^105, 106^. We identified the presence of critical beta-adrenergic receptor genes, *ADRB1* and *ADRB2*, encoding beta-adrenergic receptors 1 and 2 within our organoids in each condition **(Supplementary Fig. 26)**. While *ADRB2* was expressed in both the ACM and VCM clusters in each condition*, ADRB1* showed expression within the ACM and VCM clusters in MM, EMM1 and EMM2/1 conditions, but were only expressed in the ACM cluster in the control condition. *ADRB3* was sparsely expressed relative to *ADRB1* and *ADRB2*, which stays true to cardiac physiology^41, 107–109^. Together, this data shows that our developmentally maturated organoid platform, specifically the EMM2/1 strategy, produces organoids that recapitulate significant electrophysiological aspects of cardiac development, physiology, and disease.

### Developmental induction promotes the emergence of a proepicardial organ and formation of atrial and ventricular chambers by self-organization

We have shown that developmentally induced heart organoids presented improved cellular, biochemical and functional properties when compared to their control counterparts and exhibited multiple features reminiscent of GD45 human fetal hearts. However, previous heart organoid attempts have been lacking in producing anatomically relevant cardiac structure and morphology to a great extent, including our previous work^15–17^. Given the significant changes observed through EMM2/1, we decided to characterize morphological changes that took place under this improved condition. Organoids were harvested on day 30 of culture and stained for WT1 (proepicardium and epicardial cells) and TNNT2 (cardiomyocytes) **(Fig. 6a)**. Organoids in each developmental induction condition displayed TNNT2^+^ and WT1^+^ cells, consistent with our previous observations^15^, indicating the presence of epicardial and cardiomyocyte populations widely distributed through the organoids. Furthermore, a budding upper region could be found in all conditions, however this region did only expand significantly in EMM2/1 conditions, followed by an enlargement of this region to give rise to a well-developed secondary chamber (**Fig. 6a**). In EMM2/1, WT1^+^ cells were found densely covering the outer surface of the budding region, while existing in scattered, distant populations on the surface of the lower region. TNNT2^+^ cells were densely packed forming a thick myocardial wall in the lower chamber, while also present in the upper region in a less dense arrangement directly underneath the WT1^+^ cells. These staining patterns were not observed in the control, MM or EMM1 culture conditions. We quantified the area of WT1^+^ and TNNT2^+^ chambers across all maturation conditions **(Fig. 6b, c).** We found no difference in WT1+ area in MM organoids relative to control, whereas EMM1 and EMM2/1 organoids displayed increased areas (fold change) of 2.34 ± 0.201 and 2.11 ± 0.335, respectively. Additionally, we found no difference in TNNT2^+^ chamber area in MM and EMM1 organoids relative to control but found EMM2/1 organoids to display an increased area (fold change) of 1.62 ± 0.099 relative to control. This data shows that organoids in all conditions undergo significant morphological changes that lead to highly specific and reproducible cellular organization, including the emergence of an organoid with advanced myocardial dual-chamber morphology as well as a proepicardial pole. Upon further examination, we could determine that ventricular (MYL2) and atrial (MYL7) myosins, indicative of ventricular- and atrial cardiomyocyte subpopulations respectively, were spatially restricted to a great extent, particularly in EMM2/1 organoids. **(Fig. 6d)**. All organoids expressed MYL7 throughout the bulk of the organoid, but expression was stronger in the upper chamber in EMM2/1, suggesting an atrial chamber. In the Control and MM conditions, organoids possessed MYL2 in a high variety of locations that were not restricted to a polar end of the organoid or to either chamber in particular. On the other hand, organoids in the EMM1 and EMM2/1 conditions displayed an increasing degree of organization, with 75% of organoids possessing MYL2 restricted to one polar end of the organoid in the EMM2/1 condition, suggesting the formation of a ventricular chamber **(Fig 6e)**. These findings were particularly interesting because in these EMM2/1 organoids the proepicardial region was directly located over the atrial chamber, with the ventricular chamber in the opposite side of the organoid. Overall, this organization is comparable to the anterior-posterior patterning axis present *in utero* in the forming heart tube (see **Fig. 7a** for a schematic).

**Fig. 6.**
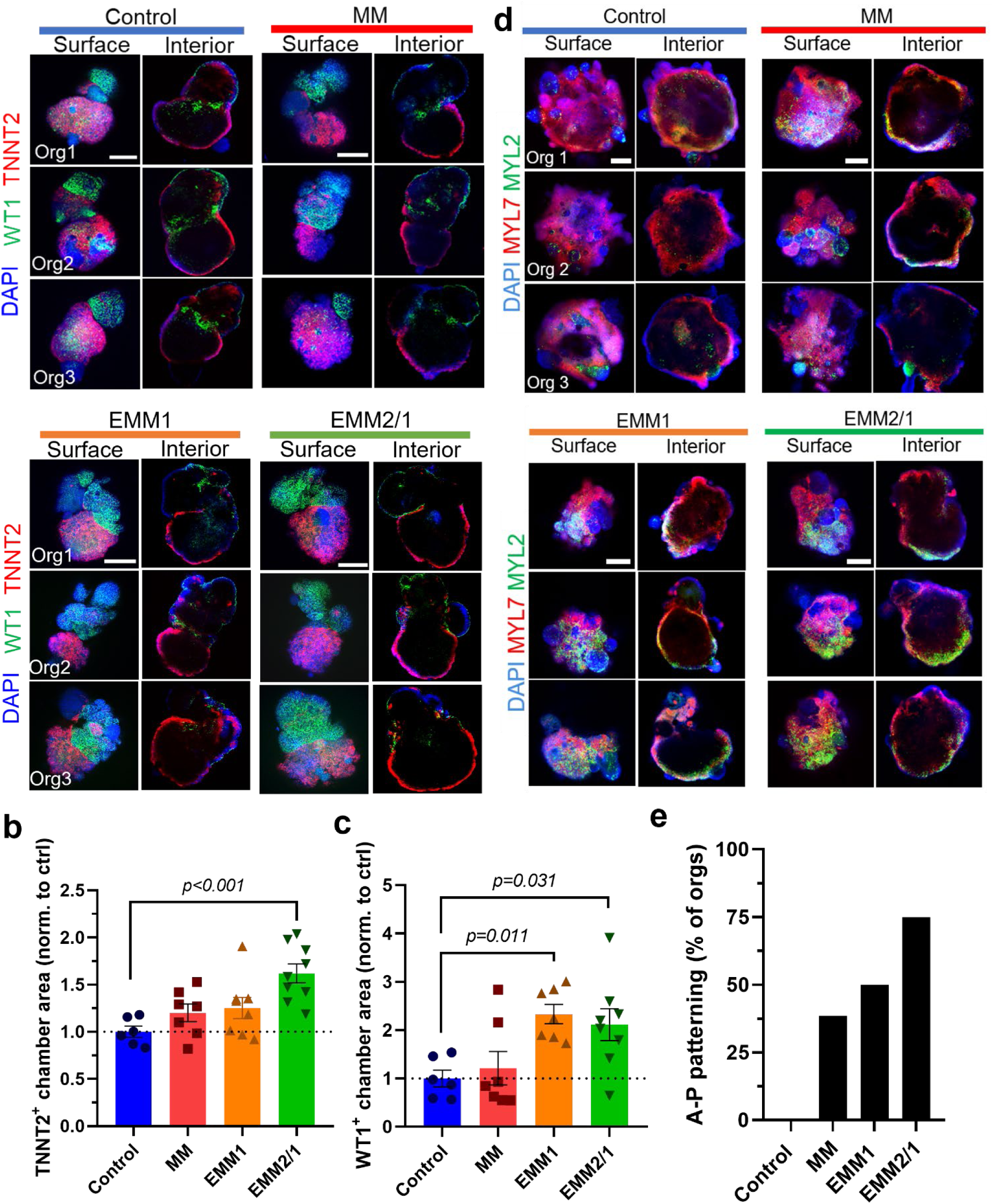
Developmental induction promotes the emergence of a proepicardial organ and formation of well-developed atrial and ventricular chambers by self-organization. **a,** Representative surface and interior immunofluorescence images of individual day 30 organoids in all conditions. Three organoids are displayed for each condition (n=7-8 organoids per condition). DAPI (blue), TNNT2 (red), WT1 (green). Scale bars = 400 μm. **(b,c,)** Quantification of TNNT2+ chamber area **(b)** and WT1+ chamber area **(c)** in each condition from images presented in **(a)** (n=7-8 organoids per condition). One-way ANOVA with Dunnett’s multiple comparisons test. **d,** Representative surface and interior immunofluorescence images of individual day 30 organoids in all conditions. Three organoids are displayed for each condition (n=9-13 organoids per condition). DAPI (blue), MYL7 (red), MYL2 (green). Scale bars = 400 μm. **e,** Quantification of percent of organoids with MYL2-MYL7 anterior-posterior patterning in each condition from images presented in **(d)** (n=9-13 organoids per condition).

**Fig. 7.**
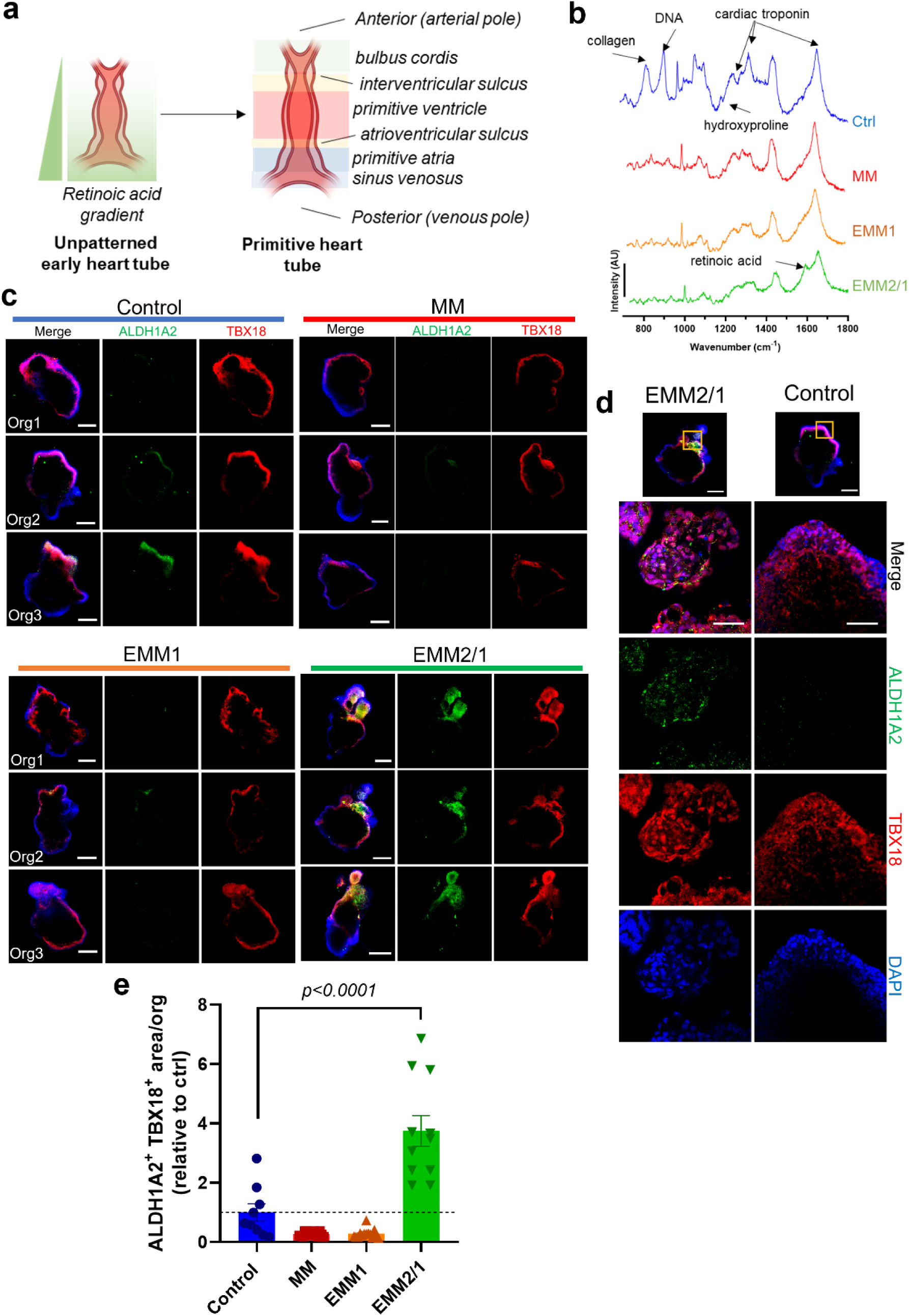
An endogenous retinoic acid gradient is responsible for spontaneous anterior-posterior heart tube patterning. **a,** Schematic portraying *in utero* cardiac heart tube formation, highlighting the localization and intensity of the retinoic acid gradient from the anterior (arterial pole) to the posterior (venous pole) of the primitive heart tube. **b,** Raman spectroscopy intensity plots for organoids from all four developmental maturation conditions at day 30 of culture. Peaks of interest are marked, such as DNA, cardiac troponin, and retinoic acid. Data presented is representative of n=3 organoids per condition. **c,** Representative immunofluorescence images of day 30 organoids in all conditions (n=9-14 organoids per condition). ALDH1A2 = green, TBX18 = red, DAPI = blue. Scale bar = 200 μm. **d,** High magnification images of EMM2/1 and Control organoids shown in **(c)**. Yellow square in image at top (Scale bar = 200 μm) represents area of high magnification. ALDH1A2 = green, TBX18 = red, DAPI = blue. Scale bar = 50 μm. **e,** Quantification of ALDH1A2^+^ TBX18^+^ area within organoids in each condition (n=9-14 organoids per condition). Data presented as fold change relative to control organoid mean value. Values = mean ± s.e.m., one-way ANOVA with Dunnett’s multiple comparisons test.

We also assessed vasculature formation in developmental induction conditions. Endothelial cell (PECAM1+) vasculature formation was examined at day 30 of culture via immunofluorescence and confocal microscopy **(Extended Data Fig. 4, Supplementary Videos 9-12)**. Assessment of organoids on surface and interior planes revealed the presence of endothelial cells amongst and lining the myocardial regions of all organoids **(Extended Data Fig. 4a).** Organoids in the EMM1 and EMM2/1 conditions presented less PECAM1+ cells than control and MM organoids. Control and MM organoids displayed robust, interconnected endothelial cell networks and throughout myocardial (TNNT2+) tissue. We quantified total PECAM1+ area, and MM organoids showed no significant difference compared to control organoids, while EMM1 and EMM2/1 organoids possessed only 52% and 61% of PECAM1+ area compared to controls, respectively **(Extended Data Fig. 4b)**. High magnification images of organoids further show the morphological transitory state of the endothelial cells within cardiomyocyte-rich regions **(Extended Data Fig. 5).** Overall, this data suggests that vascularization of the organoids might be partially eclipsed by factors in EMM1 and EMM2/1 conditions, possibly due to timing or concentration of growth factors, and will require further investigation to fine tune medium conditions.

### An endogenous retinoic acid gradient is responsible for the spontaneous anterior-posterior heart tube patterning in EMM2/1 organoids

The emergence of a spatially restricted retinoic acid gradient originating at the posterior pole of the heart tube (produced by the epicardium and primitive atrium) is a critical developmental step in heart development in mammals. This gradient establishes the anterior-posterior axis that provides cues for the formation of the ventricles and the inflow and outflow tract, while also contributing to the specification of cardiogenic progenitors, and possible other structures^110, 111^ **(Fig. 7a)**. To determine whether the heart tube-like structure observed in EMM2/1 conditions (**Fig. 6**) was indeed reminiscent of retinoic acid-mediated heart patterning, we performed Raman microscopy to detect its molecular signature using a microscope designed for this purpose (**Supplementary Fig. 31)**. We identified the presence of myosin, troponin T, tropomyosin, collagen I and other related molecular signatures in organoids in all conditions as expected, and the presence of retinoic acid specifically in EMM2/1 **(Fig. 7b)**. Retinoic acid synthesis is carried out largely by retinaldehyde dehydrogenase 2 (ALDH1A2) during embryogenesis^111–113^. To assess the localization of retinoic acid production in our organoids at day 30, we performed immunostaining with antibodies for ALDH1A2 and for TBX18 (an epicardial transcription factor, to label the proepicardial organ/atrial pole)^51–53^. We found that organoids in the EMM2/1 condition possess a localized, polarized expression of ALDH1A2 which colocalizes with TBX18^+^ cells, confirming that the retinoic acid gradient patterning the organoids was coming from the proepicardial/atrial pole (posterior pole of the heart tube *in utero*) **(Fig. 7a, c, d)**. Control, MM and EMM1 organoids did not display ALDH1A2 expression. We quantified the area of colocalization between ALDH1A2 and TBX18 and show that organoids in the EMM2/1 condition are significantly more responsive to the induction of retinoic acid synthesis **(Fig. 7e)**. Together, this data shows the ability of EMM2/1 organoids to endogenously synthesize retinoic acid in a spatially restricted manner colocalized with the epicardium (TBX18), a phenomenon that closely mimics the processes observed in *in utero* heart development and primitive heart tube patterning.

### Optical coherence tomography reveals dynamics of chamber formation in real time

We decided to employ optical coherence tomography (OCT) to live image organoids over time, and to measure the growth and monitor dynamics of chamber development under developmental induction conditions via a custom-made OCT microscopy system amenable to high-content screening^114, 115^ **(Fig. 8a)**. We found that chambers exhibited a highly dynamic behavior initially and coalesced into larger structures as time passed. EMM2/1 conditions led to the largest internal chambers within our human heart organoids between day 20 and day 30 of culture, with typically two large internal chambers as previously observed by confocal microscopy **(Fig. 8b, c, and Supplementary Videos 13-16)**. While MM organoids displayed a single internal chamber, organoids grown in the control, EMM1 and EMM2/1 conditions possessed multiple, smaller, interconnected chambers. Control and EMM2/1 organoids possessed chambers throughout the bulk of the organoids while EMM1 organoids showed chambers predominantly towards one side of the organoid. These data confirmed the formation of well-established cardiac chambers and further supported our observations on the effects of developmental induction conditions.

**Fig. 8.**
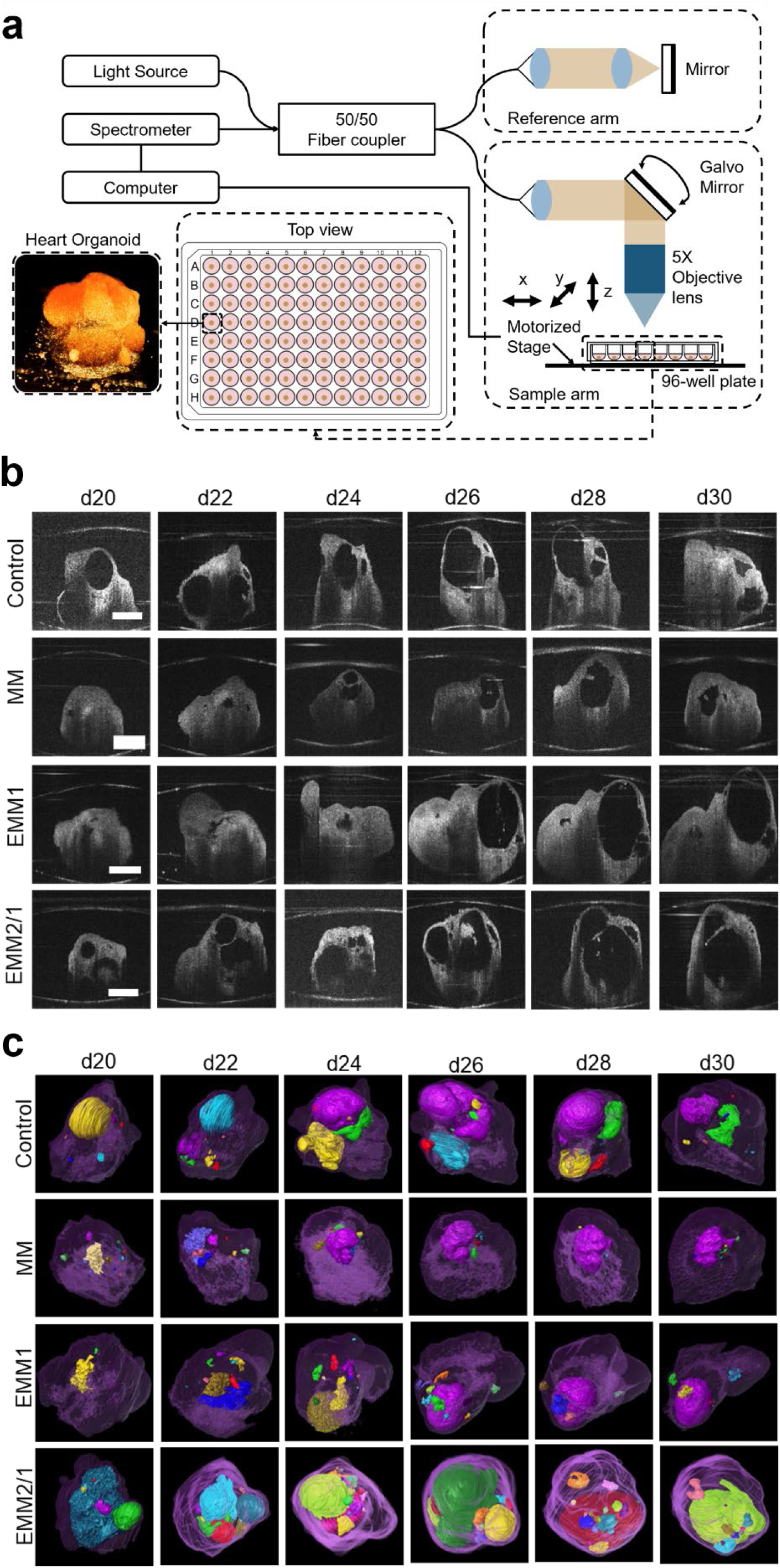
Live longitudinal imaging by optical coherence tomography reveals large, interconnected chambers within human heart organoids. **a,** Schematic of custom-built optical coherence tomography (OCT) system for human heart organoid imaging. **b,** Longitudinal OCT cross-sectional scans of human heart organoids from day 20 to day 30 in each condition. Scale bars = 500 μm. **c,** 3D segmentation of OCT scans reveal the temporally dynamic volumetric visualization of chamber identity in each condition.

## Discussion

Laboratory models of the human heart have made considerable progress over the last decades, beginning with animal models and primary cardiomyocyte culture and moving onwards to induced pluripotent stem cell-derived cardiac tissues (e.g. cardiomyocytes) and tissue engineering approaches (3D printing, biomaterials). The latest advances in human heart models are heart organoids generated from pluripotent stem cells^15–17, 116^. While these model systems have certainly yielded transformative research findings in the fields of cardiac disease, heart development, and cardiac toxicity testing^5, 15, 117–119^, these systems do not possess the true complexity of the *in utero* human heart, owing to a lack of maturity and faithfulness to human physiology, morphology, cellular organization, and functionality. These shortcomings severely limit the scope of relevance of traditional model systems. To circumvent these limitations, we designed and implemented relatively simple and highly reproducible methods for developmental induction strategies inspired by *in utero* biological steps, producing human heart organoids with higher anatomical complexity and physiological relevance along first trimester fetal development. Among these strategies, we found that EMM2/1 most closely recapitulated heart development *in vitro* (see ***Materials and Methods*** for composition details), and enabled organoids to acquire high levels of complexity and anatomical relevance by inducing progressive mitochondrial and metabolic maturation, electrophysiological maturation, increased morphological and cellular complexity, and most importantly, recapitulating anterior-posterior heart tube patterning by endogenous retinoic acid signaling and self-organization.

Single-cell gene expression for key genes relating to multiple cell clusters revealed that EMM2/1 organoids, compared to organoids from other maturation strategies, yielded the highest similarity to *in vivo* 6.5 post-conception week (GD45) developing human hearts^41^. Implementation of developmental induction strategies did not lead to the emergence of new cardiac lineages as the same cardiac cell types were observed in all conditions but did lead to expansion and reduction of certain populations, such as atrial and ventricular cardiomyocytes, and mesenchymal cell types (stromal cells) in what seems to be a process of fine tuning and remodeling. Interestingly, we could also observe the appearance of valvular and conductance cell types throughout all developmental induction conditions, a phenomenon not described before in heart organoids. The metabolic transition from glycolysis to fatty acid oxidation is a paramount step in the late stages of cardiac development, preparing the heart for increased energy expenditure as well as inducing transcriptional regulation and stimulating physiological maturation^28, 30, 77–79, 120^. Efforts have been pursued to simulate these phenomena in vitro with cardiomyocytes and engineered heart tissues and have found beneficial effects from modified glucose concentrations and the addition of fatty acids^7, 20, 21, 23, 121^. However, these systems are simplistic models and do not possess the high physiological complexity as observed in human heart organoids. We showed that human heart organoids respond dramatically to developmental maturation stimuli and metabolically maturate and possess increased mitochondrial growth, density, and gene expression, particularly through the EMM2/1 strategy. These dramatic responses compared to traditional methods may be the result of synergy between multiple cardiac cell subtypes, such as epicardial cells and cardiac fibroblasts, which have been shown to stimulate cardiomyocyte growth and function^122, 123^.

Proper and gradual electrophysiological maturation throughout the cardiac syncytium, including the complex interplay between various ion channels and their subtypes as well as depolarization through t-tubules, comprises critical aspects of cardiac development and functionality^85–88, 92, 93, 124^. Here, we show that organoids from the EMM2/1 strategy develop distinct calcium transients with increasing physiological mimicry due to their increased amplitude and decreased frequency. Additionally, organoids in the EMM2/1 strategy develop higher levels of t-tubules, inward-rectifying potassium ion channels, and hERG channels compared to other maturation strategies. In fact, many efforts towards in vitro cardiomyocyte maturation have struggled or failed to elicit the presence of t-tubules^125, 126^ and inward-rectifying potassium ion channels remain critical to establishing low resting membrane potential^127, 128^. Moreover, cardiac hERG channels represent a paramount channel of importance for pharmacological screening due to its high arrhythmogenic potential if interfered with^100, 101^. Nonetheless, the summation of multiple ion transients results in the cardiac action potential which is the ultimate driving force for human heart contraction and functionality. We show that cardiomyocytes within our EMM2/1 organoids possess ventricular-, atrial-, and nodal-like action potentials, opening the door for electrophysiological applications in drug screening.

The embryonic heart begins as an unpatterned heart tube and undergoes cellular and structural changes through morphogenetic signaling events to pattern along the anterior-posterior axis, loop, and eventually form the 4-chambered heart^129, 130^. We investigated the morphological landscape of our organoids following the application of our maturation strategies and found that EMM2/1 organoids formed a two-chambered structure with cardiomyocytes forming one chamber with atrial identity and another with ventricular fate. Dense epicardial layering at the atrial chamber identified the proepicardial organ as the posterior pole of the heart tube^131^ and revealed that EMM2/1 organoids were spontaneously patterning along the aforementioned anterior-posterior axis, a phenomenon that was exclusively observed in EMM2/1 organoids. Further investigations found that self-organization and patterning in EMM2/1 was driven by an endogenous retinoic acid signaling gradient, identified by using a combination of Raman and confocal microscopy. ALDH1A2, an enzyme required for retinoic acid synthesis, was observed to be spatially restricted to the posterior end of the organoid, and co-localized with TBX18, an epicardial transcription factor confirming that the proepicardial organ was functional. Taken together, these data support the hypothesis that our organoids recapitulate events that take place during *in utero* gestation where the proepicardial organ surrounds the posterior pole of the patterned heart tube where posterior atrial cardiomyocytes and proepicardial cells produce retinoic acid to form a signaling gradient that further instructs the remainder of the heart tube with patterning and specification information^41, 51, 53, 111, 131^

While our maturated human heart organoid technology opens exciting avenues for modeling the human heart *in vitro*, important limitations remain. First, further investigation is necessary to clarify the role of other important cellular populations, such as conductance cells and endothelial/coronary vasculature. EMM2/1 conditions had negative effects on organoid vascularization, a topic that deserves more attention as conditions can be further refined. Second, the lack of circulation constitutes a significant drawback that needs to be addressed, possibly through using microfluidic devices. Third, further developmental steps need to be introduced to continue increasing the physiological relevance of the organoids. This includes correcting the lack of embryonic tissue-resident macrophage populations or the contributions of the neural crest. Finally, more anatomical events need to be modeled, such as outflow tract and atrioventricular channel formation, heart looping and septation.

In summary, we describe here a developmental induction strategy inspired by hallmarks of *in utero* development for producing patterned human heart tube organoids highly recapitulating human first trimester heart development. Our EMM2/1 developmental induction strategy yields several unique and crucial characteristics representative of an early fetal human heart, including the presence of anterior-posterior patterning with an endogenous retinoic acid gradient originating at the posterior end, polar separation of atrioventricular chambers, and a posterior, proepicardial pole.. The EMM2/1 strategy also results in valvular cells, conductance cells, proepicardial cells and more, as well as large hollow chambers, functional electrophysiology, and increased mitochondrial density and metabolic transcriptional profiles similar to that of gestational human hearts. To the extent of our knowledge, this is the first time the human heart tube has been reconstructed to this level of detail *in vitro*. All these developmental features highlight the ability to model the developing human heart in vitro and prove the potential of our methodology for establishing heart models for the study of normal heart development, congenital heart defects, cardiac pharmacology, regeneration, and possibly other cardiovascular disorders in the future.

## Materials and Methods

### Stem cell culture

The following human induced pluripotent stem cell lines were used for this study: iPSC-L1, AICS-0061-036 (Coriell Institute for Medical Research, Alias: AICS). Pluripotency and genomic stability were tested for all hiPSC lines used. hiPSCs were cultured in Essential 8 Flex medium with 1% penicillin streptomycin (Gibco) in 6-well plates on growth factor reduced Matrigel (Corning) inside an incubator at 37 °C and 5% CO_2_. hiPSCs were passaged using ReLeSR passaging reagent (STEMCELL Technologies) upon reaching 60-80% confluency.

### Self-assembling human heart organoid differentiation

Step-by-step, detailed protocols which describe the generation and differentiation of the human heart organoids are provided^14^. In brief, hiPSCs were grown to 60% confluency on 6-well plates and dissociated using Accutase (Innovative Cell Technologies) to obtain a single-cell solution. hiPSCs were collected and centrifuged at 300 *g* for 5 minutes and resuspended in Essential 8 Flex medium containing 2μM ROCK inhibitor (Thiazovivin) (Millipore Sigma). hiPSCs were counted using a Moxi cell counter (Orflo Technologies) and were seeded at a concentration of 10,000 cells per well in round bottom 96 well ultra-low attachment plates (Costar) on day −2 in a volume of 100 μL. The plate was then centrifuged at 100 *g* for 3 minutes and subsequently placed inside a 37 °C and 5% CO_2_ incubator. After 24 hours (day −1), 50 μL was removed from each well and 200 μL of fresh Essential 8 Flex Medium was added to each well to obtain a final volume of 250 μL per well. The plate was then placed inside a 37 °C and 5% CO_2_ incubator. After 24 hours (day 0), 166 μL of medium was removed from each well. Then, 166 μL of RPMI with B27 supplement without insulin (Gibco) supplemented with 1% penicillin streptomycin (Gibco) (hereafter termed “RPMI/B27 minus insulin”) containing CHIR99021, BMP4, and Activin A was added to each well to obtain final concentrations of 4 μM CHIR99021, 36 pM (1.25 ng/mL) BMP4, and 8 pM (1.00 ng/mL) Activin A. The plate was subsequently placed inside a 37 °C and 5% CO_2_ incubator. After exactly 24 hours (day 1), 166 μL of medium was removed from each well and replaced with 166 μL of fresh RPMI/B27 minus insulin. On day 2, 166 μL of medium was removed from each well and 166 μL of RPMI/B27 minus insulin with Wnt-C59 (Selleck) was added to obtain a final concentration of 2 μM Wnt-C59 inside each well. The plate was then incubated for 48 hours. On day 4, 166 μL was removed and replaced with fresh RPMI/B27 minus insulin and incubated for 48 hours. On day 6, 166 μL was removed and replaced with 166 μL RPMI with B27 supplement (with insulin) and 1% penicillin streptomycin (hereafter termed RPMI/B27). The plate was incubated for 24 hours. On day 7, 166 μL of media was removed from each well and 166 μL of RPMI/B27 containing CHIR99021 was added to obtain a final concentration of 2 μM CHIR99021 per well. The plate was incubated for 1 hour. After 1 hour, 166 μL of medium was removed from each well and 166 μL of fresh RPMI/B27 was added to each well. The plate was incubated for 48 hours. From days 9 to 19, every 48 hours, media changes were performed by removing 166 μL of media from each well and adding 166 μL of fresh RPMI/B27.

### Developmental induction conditions

Organoids were generated and differentiated according to the protocol outlined previously. Beginning on day 20, organoids were subjected to various maturation medium conditions. The control strategy is a continuation of culture within RPMI/B27 from day 20 to day 30, performing standard media changes every 48 hours. The maturation medium (MM) strategy is employed from day 20 to day 30, performing media changes every 48 hours using MM media, consisting of stock RPMI/B27 with 52.5 μM palmitate-BSA, 40.5 μM oleate-BSA (Sigma), 22.5 μM lineolate-BSA (Sigma), 120 μM L-Carnitine (Sigma) and 30 nM T3 hormone (Sigma). The enhanced maturation medium 1 (EMM1) strategy is employed from day 20 to day 30, performing media changes every 48 hours using EMM1 media, consisting of stock RPMI 1640 Medium, no glucose (Gibco) supplemented with B27 (with insulin), 1% penicillin streptomycin (Gibco), 52.5 μM palmitate-BSA, 40.5 μM oleate-BSA (Sigma), 22.5 μM lineolate-BSA (Sigma), 120 μM L-Carnitine (Sigma), 30 nM T3 hormone (Sigma), 0.4 mM ascorbic acid (Thermo Fisher Scientific) and 4 mM Glucose (Gibco). The enhanced maturation medium 2/1 (EMM2/1) strategy is employed from day 20 to day 30, performing media changes every 48 hours, utilizing a combination of two medias. From day 20 to day 26, EMM2 media is utilized which consists of stock RPMI 1640 Medium, no glucose (Gibco) supplemented with B27 (with insulin), 1% penicillin streptomycin (Gibco), 52.5 μM palmitate-BSA, 40.5 μM oleate-BSA (Sigma), 22.5 μM lineolate-BSA (Sigma), 120 μM L-Carnitine (Sigma), 30 nM T3 hormone (Sigma), 0.4mM ascorbic acid (Thermo Fisher Scientific), 4 mM Glucose (Gibco) and 50 ng/mL IGF-1 (Sigma). Continuing the EMM2/1 strategy, from day 26 to day 30, EMM1 media is utilized. Organoids are collected on day 30 for analysis.

### Immunofluorescence

Human heart organoids were transferred from the round bottom ultra-low attachment 96 well plate to 1.5 mL microcentrifuge tubes (Eppendorf) using a cut 200 μL pipette tip (to increase tip bore diameter as to not disturb the organoid). Organoids were fixed in 4% paraformaldehyde (VWR) in PBS for 30 minutes. Following, organoids were washed using PBS-Glycine (1.5 g/L) three times for 5 minutes each. Organoids were then blocked and permeabilized using a solution containing 10% Donkey Normal Serum (Sigma), 0.5% Triton X-100 (Sigma), and 0.5% BSA (Thermo Fisher Scientific) in PBS on a thermal mixer at 300rpm at 4 °C overnight. Organoids were then washed 3 times using PBS and incubated with primary antibodies (Supplementary Table 1) within a solution containing 1% Donkey Normal Serum, 0.5% Triton X-100, and 0.5% BSA in PBS (hereafter termed “Antibody Solution”) on a thermal mixer at 300rpm at 4 °C for 24 hours. Following, organoids were washed 3 times for 5 minutes each using PBS. Organoids were then incubated with secondary antibodies (Supplementary Table 1) in Antibody Solution on a thermal mixer at 300rpm at 4 °C for 24 hours in the dark. Subsequently, organoids were washed 3 times for 5 minutes each using PBS and mounted on glass microscope slides (Fisher Scientific). 90 μm Polybead Microspheres (Polyscience, Inc.) were placed between the slide and a No. 1.5 coverslip (VWR) to provide support pillars such that the organoids could retain three dimensionality. Organoids were transferred to the glass microscope slides using a cut 200uL pipette tip and mounted using a clearing solution described previously^132^. T-tubule staining was performed using FITC-conjugated Wheat Germ Agglutinin (WGA) lectins (Sigma).

### Confocal Microscopy and Image Analysis

Immunofluorescence images were acquired using a confocal laser scanning microscope (Nikon Instruments A1 Confocal Laser Microscope). Images were analyzed using Fiji.

### Heart organoid dissociation

Organoids were collected on day 30 from each maturation strategy (Control, MM, EMM1, EMM2/1). Organoids were individually placed into separate 1.5mL microcentrifuge tubes (Eppendorf), dissociated and pooled. Organoids were dissociated into a single-celled suspension using a modified protocol of the STEMdiff Cardiomyocyte Dissociation Kit (STEMCELL Technologies). Upon being transferred to a microcentrifuge tube, organoids were washed with PBS, submerged in 200 μL of warm dissociation media (37 °C), and placed on a thermal mixer at 37 °C and 300rpm for 5 minutes. Then, the supernatant was collected and transferred to a 15 mL falcon tube (Corning) containing 5mL of respective media (Control, MM, EMM1, etc.) containing 2% BSA (Thermo Fisher Scientific). An additional 200 μL of warm dissociation media (37 °C) was then added back to the organoid on a thermal mixer (37 °C). The organoid dissociation media solution was then pipetted up and down gently 3-5 times. The organoid was allowed to sit on the thermal mixer for an additional 5 minutes. If the organoid remained visible, the process was repeated. Once the organoid was no longer visible, the microcentrifuge tube solution was pipetted up and down gently 3-5 times and its entire contents were transferred to the 15 mL falcon tube containing the respective media + 2% BSA and cells. These tubes were then centrifuged at 300 *g* for 5 minutes. The supernatant was aspirated, and the cell pellets were resuspended in respective media + 2% BSA. Using a hemocytometer, viability, cell counts, and aggregate percentage were acquired.

### Single cell RNA sequencing

Libraries were prepared using the 10x Chromium Next GEM Single Cell 3’ Kit, v3.1 and associated components. Completed libraries were QC’d and quantified using a combination of Qubit dsDNA HS, Agilent 4200 TapeStation HS DNA1000 and Invitrogen Collibri Library Quantification qPCR assays. The libraries were pooled in equimolar proportions and the pool quantified again using the Invitrogen Collibri qPCR assay. The pool was loaded onto two lanes of an Illumina NovaSeq 6000 SP flow cell (v1.5) and sequencing was performed in a custom paired end format, 28 cycles for read 1, 2 10 cycle index reads and 90 cycles for read 2. A v1.5, 100 cycle NovaSeq reagent cartridge was used for sequencing. The 28bp read 1 includes the 10x cell barcodes and UMIs, read 2 is the cDNA read. Output of Real Time Analysis (RTA) was demultiplexed and converted to FastQ format with Illumina Bcl2fastq v2.20.0. After demultiplexing, reads from each of the sample libraries were further processed using 10x Genomics cellranger count (v6.1.2). Analysis of files was performed using 10X Genomics Loupe Browser v6.3.0 using k-means clustering of 8 clusters and UMAP visualization. Enrichr^133–135^ was used to assess gene ontologies. Pathview Web was used to generate biological pathway graphs^83, 84^. Cell-cell communication analysis was performed using Liana^136^ and Celltalker^137^ (https://github.com/arc85/celltalker). To accomplish this task, the counts and clusters data from Loupe Browser were imported into Seurat^138^. Both programs perform differential expression between clusters using standard Seurat findmarkers differential expression function and then rank pairs of ligands and receptors by significant p-value and log2fold change. To identify pairs of ligands and receptors pairs, Liana uses OmniPath database^139^, and Celltalker uses Ramilowski-pairs database for ligands-receptors interactions^140^.

### Transmission Electron Microscopy (TEM)

Human heart organoids were fixed on Day 15 and Day 30 in 2.5% glutaraldehyde (Electron Microscopy Solutions) in PBS for 45 minutes, washed three times in PBS for 5 minutes each, then stored at 4 °C. Samples were then washed with 100mM phosphate buffer and postfixed with 1% osmium tetroxide in 100mM phosphate buffer, dehydrated in a gradient series of acetone and infiltrated and embedded in Spurr (Electron Microscopy Sciences). 70 nm thin sections were obtained with a Power Tome Ultramicrotome (RMC, Boeckeler Instruments. Tucson, AZ) and post stained with uranyl acetate and lead citrate. A JEOL 1400Flash Transmission Electron Microscope (Japan Electron Optics Laboratory, Japan) was used to acquire images at an accelerating voltage of 100k.

### Calcium imaging

Calcium transient activity within the human heart organoids was assessed using Fluo-4 AM (Thermo Fisher Scientific). Fluo-4 AM was solubilized in DMSO per the manufacturer’s instructions. A 1.5 μM solution of Fluo-4 was prepared in respective medium (Control, MM, EMM1, etc.). Organoids were washed twice using RPMI 1640 basal medium, then Fluo-4 AM was added at a final concentration of 1 μM and incubated for 30 minutes at 37 °C and 5% CO_2_. Organoids were then washed twice using their respective medium (Control, MM, EMM1, etc.) and transferred to a chambered coverglass slide (Cellvis) using a cut 200 μL pipette tip. Videos were acquired using a Cellvivo microscope (Olympus) at 100 frames per second over 10 total seconds as an image stack. Samples were excited at 494 nm excitation and 506 nm emission was collected. Data was processed using Fiji. Baseline F_0_ of fluorescence intensity F was calculated using the lowest 50 intensity values in the acquired dataset. Fluorescence change ΔF/F_0_ was calculated using the equation:

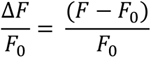

### Voltage imaging

Voltage activity within human heart organoids was assessed using di-8-ANEPPS (Thermo Fisher Scientific). Di-8-ANEPPS was solubilized in DMSO per the manufacturer’s instructions. A 15 μM solution of di-8-ANEPPS was prepared in respective medium (Control, MM, EMM1, etc.). Organoids were washed twice using RPMI 1640 basal medium, then di-8-ANEPPS was added at a final concentration of 10 μM and incubated for 30 minutes at 37 °C and 5% CO_2_. Organoids were then washed twice using their respective medium (Control, MM, EMM1, etc.) and transferred to a chambered coverglass slide (Cellvis) using a cut 200 μL pipette tip. Videos were acquired using a Cellvivo microscope (Olympus) at 100 frames per second over 10 total seconds as an image stack. Samples were excited at 465 nm excitation and 630 nm emission was collected. Data was processed using Fiji.

### Real time RT-PCR

Organoids were collected at day 20, 21, 23, 25, and 30 and stored in RNAprotect (Qiagen) at −20 °C. RNA was extracted using the Qiagen RNEasy Mini Kit according largely to the manufacturer’s instructions. Organoids were lysed using the Bead Mill 4 Homogenizer (Fisher Scientific) at speed 2 for 30 seconds. RNA concentration was measured using a NanoDrop One (Thermo Fisher Scientific). A minimum threshold of 10 ng/μL was required to proceed with reverse transcription. cDNA was generated using the Quantitect Reverse Transcription Kit (Qiagen) and stored at −20 °C. Primers for real time qPCR were designed using the Primer Quest tool (Integrated DNA Technologies). SYBR Green (Thermo Fisher Scientific) was used as the DNA intercalating dye for the reaction vessel. Real time qPCR was performed using the QuantStudio 5 Real-Time PCR system (Applied Biosystems) using a total reaction volume of 20 μL. Gene expression levels were normalized to *HPRT1* expression in each independent sample. Fold change values were obtained using the double delta CT method. At least 4 independent samples were run for each gene expression assay at each timepoint per condition. mRNA expression figures are displayed as fold change relative to Control.

### Mitochondrial staining

Intracellular mitochondrial presence within human heart organoids was visualized using Mitotracker Deep Red FM (Thermo Fisher Scientific). Mitotracker was prepared according to the manufacturer’s instructions. A 150 nM solution of Mitotracker was prepared in respective medium (Control, MM, EMM1, etc.). Organoids were washed twice using RPMI 1640 basal medium, then Mitotracker was added at a final concentration of 100 nM and incubated for 30 minutes at 37 °C and 5% CO_2_. Organoids were then washed twice using their respective medium (Control, MM, EMM1, etc.) and transferred to a chambered coverglass slide (Cellvis) using a cut 200 μL pipette tip. Images were acquired using a Cellvivo microscope (Olympus). Data was processed using Fiji.

### Raman microscopy

The Raman spectra of the organoids were acquired by using a Renishaw inVia Confocal Raman spectrometer connected to a Leica microscope (Leica DMLM, Leica Microsystems, Buffalo Grove, IL, USA). A 785 nm near-IR laser, Nikon Flour 60x NA= 1.00 water immersion objective lens, and 1,000 milli-second exposure time with the average number of 100 accumulations were used for the data acquisition of each scanning position of the organoids. To circumvent a strong background signal, a quartz slide (Chemglass Life Sciences, NJ, USA) was used as a substrate for the Raman spectra acquisition. Organoids were collected on Day 30 for analysis.

### Optical coherence tomography

A Spectral-Domain Optical Coherence Tomography (SD-OCT) system similar to our previous work^114, 115^ was used for label-free longitudinal imaging of the heart organoids. A superluminescent diode (EXALOS, EXC250023-00) was used as the light source with a center wavelength of ∼1300 nm and a 3 dB spectrum range of ∼180 nm. A spectrometer (Wasatch Photonics, Cobra 1300) based on a 2048-pixel InGaAs line-scan camera (Sensors Unlimited, GL2048) was used to provide a maximum A-scan rate of 147 kHz. A 5X objective lens was used and the transverse and axial resolutions were measured to be ∼2.83 μm and ∼3.04 μm in tissue, respectively. Longitudinal 3D OCT imaging was performed every other day from Day 20 to Day 30. Each 3D OCT scan comprised 600 A-scans per B scan and 600 B-scans. Each organoid requires ∼22 seconds for image acquisition using an exposure time of ∼40 μs for each A-scan. Eight organoids from each group were imaged and used for analysis. The media level in each well was adjusted during imaging to reduce image artifacts and minimize light absorption. Re-scaling of acquired OCT images was performed using ImageJ to obtain isotropic pixel size in x-y-z dimensions (Schneider et al., 2012). Registration of the same organoid on different days, cavity segmentation, and 3D rendering were performed using Amira software (Thermo Fisher Scientific). The total volume and cavities inside the organoids were quantified from the segmentation data.

### Statistics and reproducibility

Microsoft Excel was used to collect raw data. GraphPad Prism 9 software was used for all analyses. Data presented as normal distribution. Statistical significance was evaluated using one-way ANOVA with Dunnett or Brown-Forsyth and Welch post-test corrections when appropriate (p < 0.05). All data presented as mean ± s.e.m.

## Data availability

scRNA-Sequencing data sets have been deposited in the National Center for Biotechnology Information Gene Expression Omnibus repository under accession code GSE218582. All data generated and/or analyzed in this study are provided in the published article and its supplementary information files or can be obtained from the corresponding author upon request.

## Acknowledgements

We wish to thank the IQ and MSU Advanced Microscopy Cores for access to confocal microscopes, and the MSU Genomics Core for sequencing services. We also wish to thank all members of the Aguirre Lab for their valuable comments and advice. Work in Dr. Aguirre’s laboratory was supported by startup funds from MSU, the NIH under award numbers K01HL135464, R01HL151505, by the American Heart Association under award number 19IPLOI34660342 and by the Spectrum-MSU Foundation. Work in Dr. Zhou’s laboratory was supported by a startup funds from Washington University in St. Louis and NIH grants R01EB025209, R01HL156265 and R21EB03268401A1. Work in Dr. Qiu’s laboratory was supported by startup funds from MSU and the U.S. National Science Foundation (1808436, 1918074). Work in Dr. Park’s lab was supported by startup funds from MSU and by MSU’s Discretionary Funding Initiative.

## Author contributions

B.V. and A.A. designed all experiments and conceptualized the work. B.V. performed all experiments and data analysis. A.R. performed data analysis, scRNAseq data analysis, and cell and organoid culture. Al.K., P.M, C.O., A.H., Y.L.I and A.H.W performed cell and organoid culture. Ar.K. and S.P. performed scRNAseq data analysis. F.W. and C.Z. performed optical coherence tomography experiments and data analysis. A.J. and Z.Q. performed Raman spectroscopy experiments and data analysis. B.V. and A.A. wrote the manuscript.

---

**Extended Data Fig. 1.**
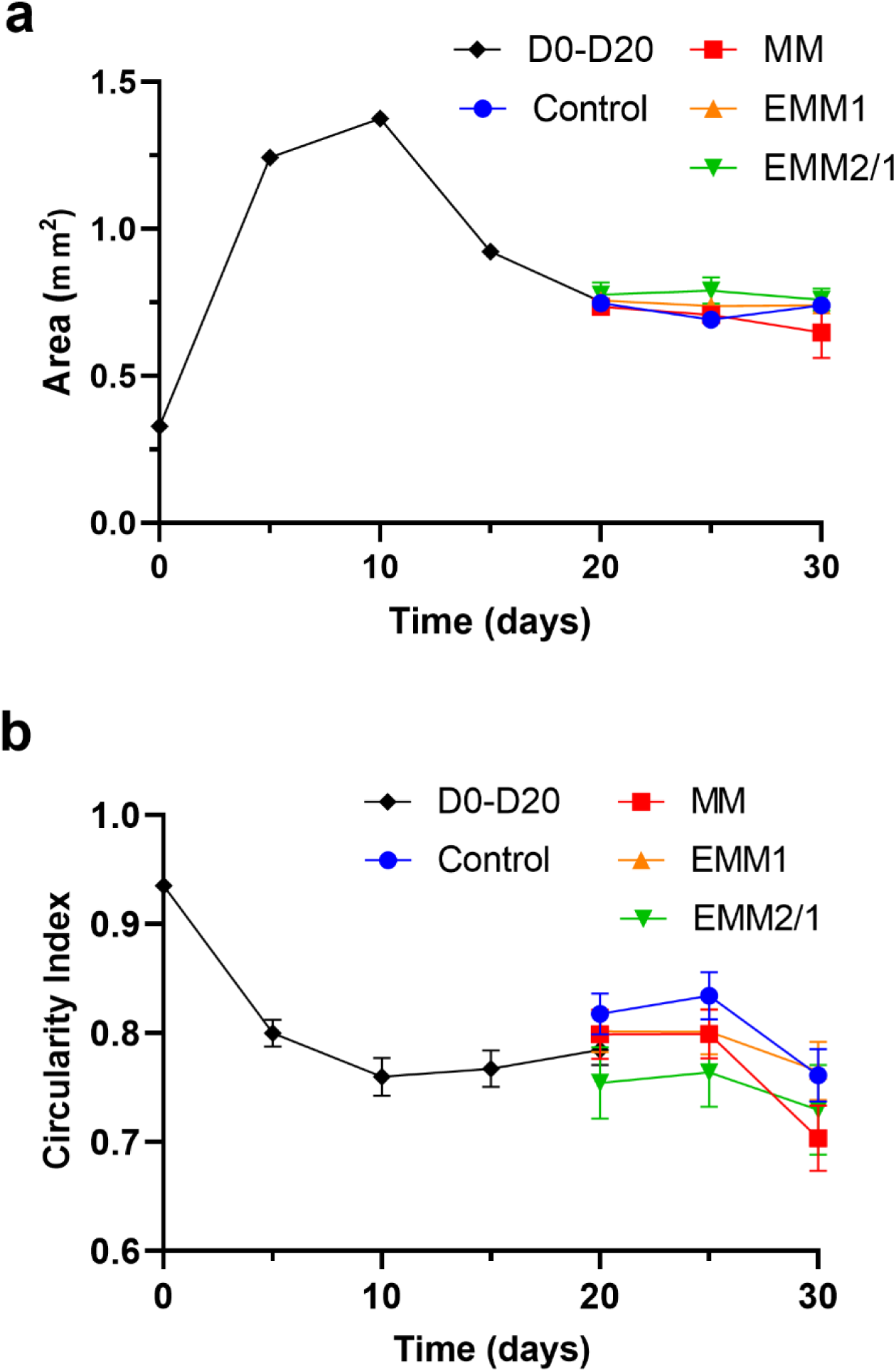
Longitudinal characterization of organoid morphology. **a,** Quantification of organoid area at 5-day intervals throughout organoid culture and maturation strategy (n=8 organoids per condition). Data presented as mean ± s.e.m. **b,** Quantification of organoid circularity at 5-day intervals throughout organoid culture and maturation strategy (n=8 organoids per condition). Data presented as mean ± s.e.m.

**Extended Data Fig. 2.**
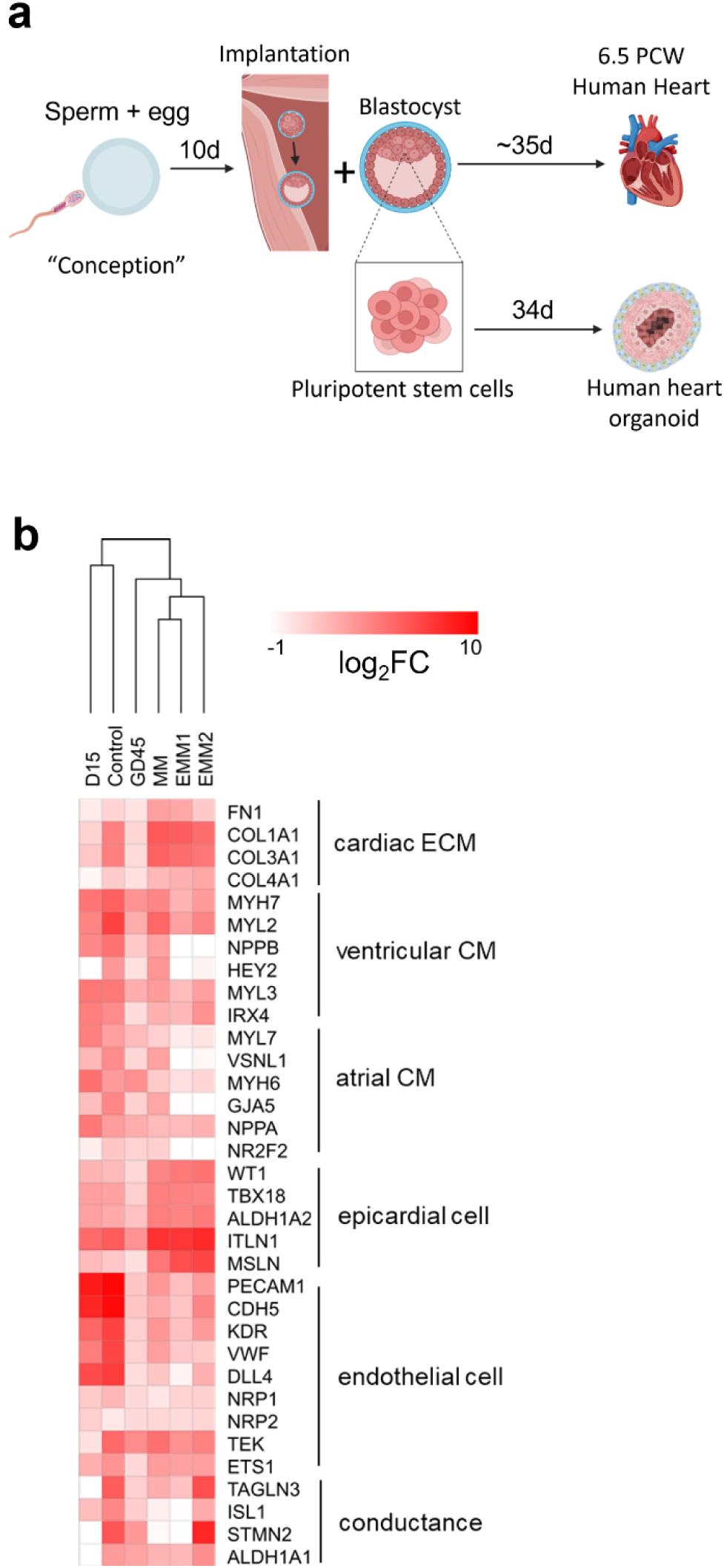
Transcriptomic organoid landscape reveals similarities to *in vivo* developing human hearts. **a,** Schematic of key human conception processes and events depicting the temporal similarities and differences between developing human hearts at a period 6.5 weeks from conception (Gestational Day 45 (GD45)) and the growth of our human heart organoids. **b,** Differential expression heatmap of log_2_ fold change values within scRNAseq datasets for day 15 organoids, day 34 organoids in the Control, MM, EMM1, and EMM2/1 conditions, and 6.5 PCW developing human hearts (GD45). Genes related to atrial and ventricular cardiomyocytes, epithelial cells, endocardial cells, epicardial cells, and cardiac ECM are displayed. Hierarchical clustering between conditions is also displayed above the heatmap. Schematic in **(a)** was created using Biorender.com.

**Extended Data Fig. 3.**
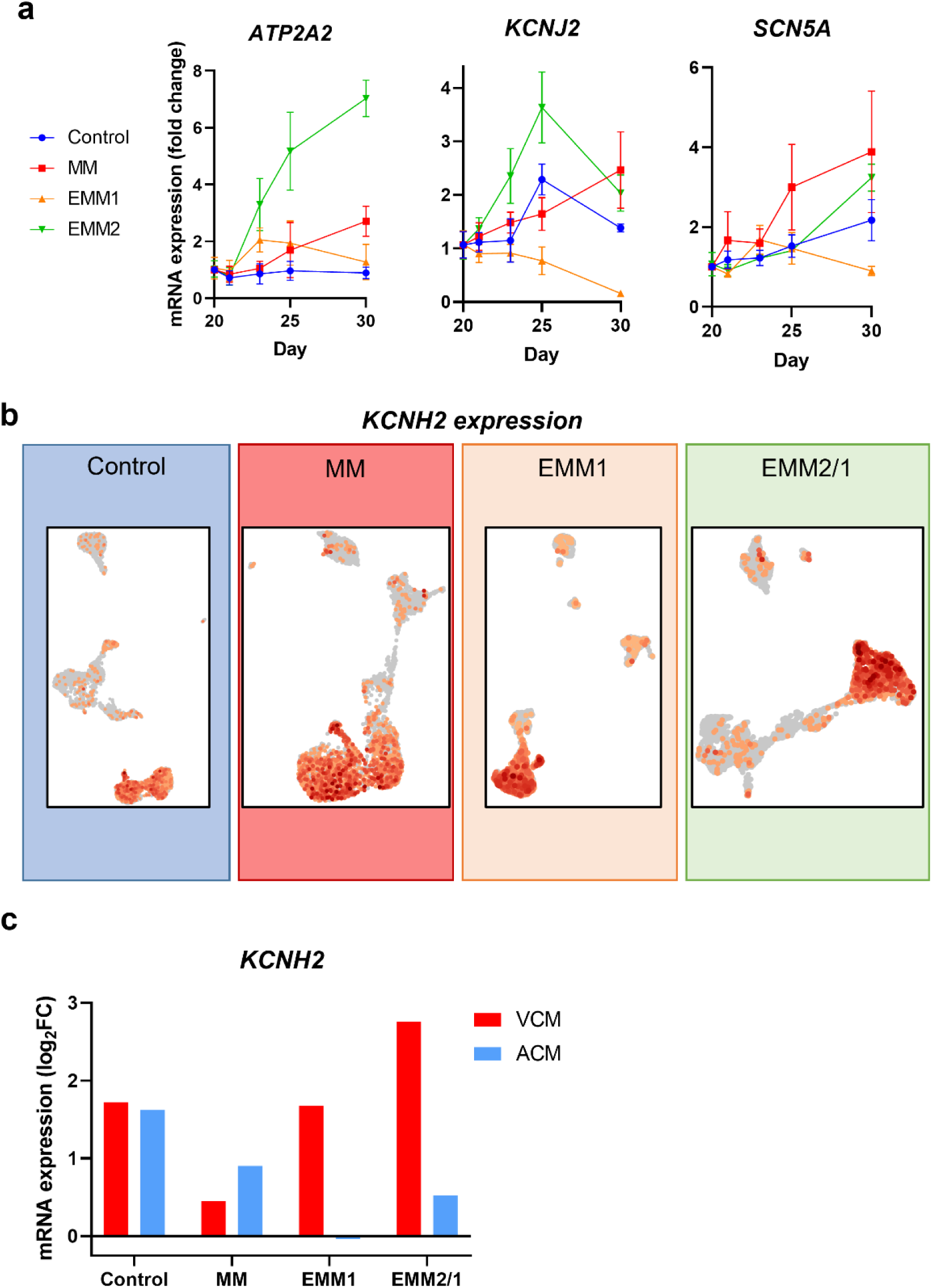
EMM2/1 organoids possess enhanced electrophysiological transcriptomes and hERG channel expression. **a,** mRNA expression of key electrophysiology-related genes between days 20 and 30 of culture for each condition (n=4 organoids). Values = mean ± s.e.m. **b,** *KCNH2* spatial expression within organoids from each condition. Data obtained from scRNAseq datasets. **c,** Quantification of *KCNH2* mRNA expression (log_2_ Fold Change) in VCM and ACM clusters in each condition.

**Extended Data Fig. 4.**
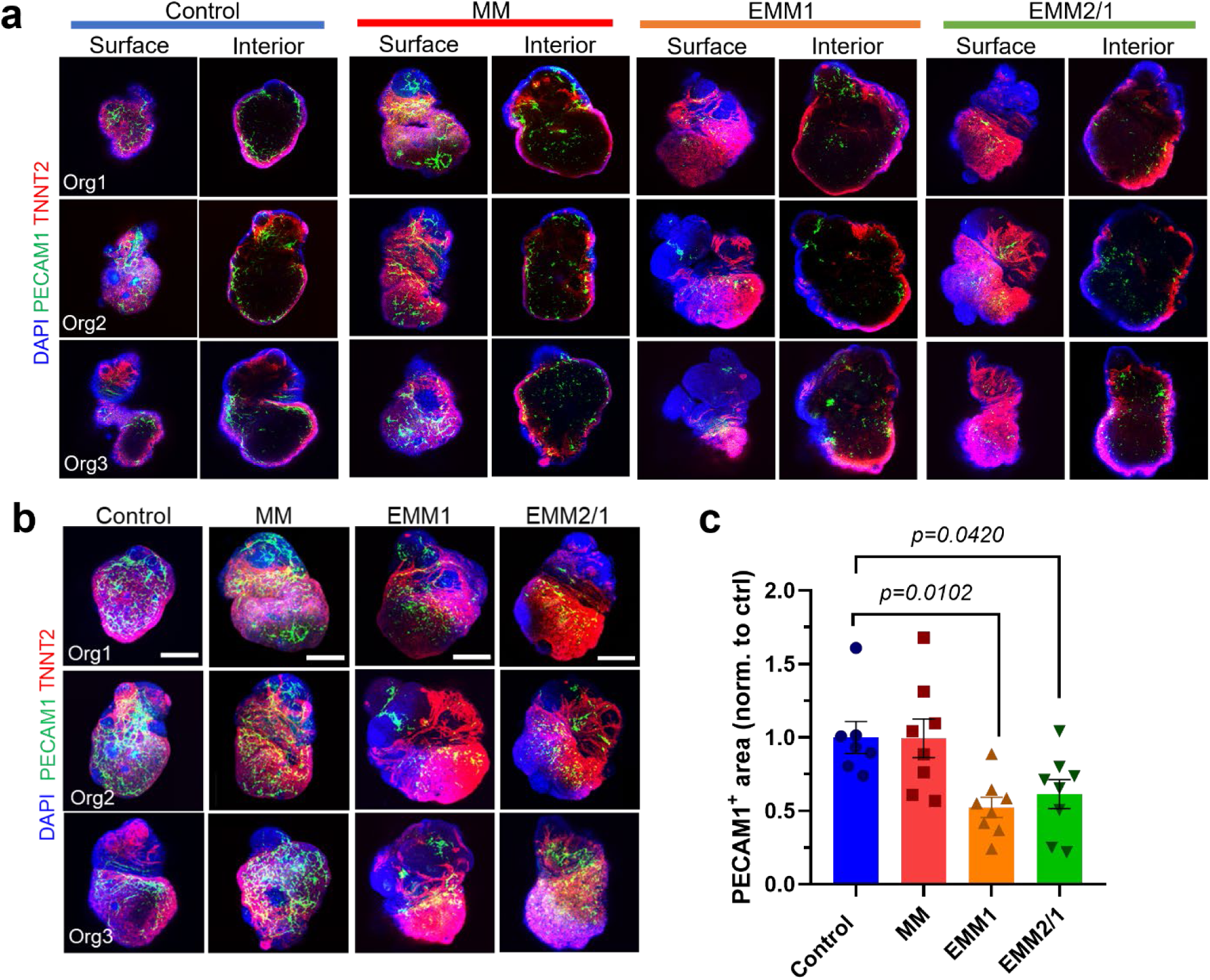
Endothelial cell localization and morphology is perturbed through enhanced developmental maturation strategies. **a,** Representative immunofluorescence images from the surface and interior of day 30 organoids with DAPI (blue), TNNT2 (red), and PECAM1 (green) in each condition (n=7-8 organoids per condition). Scale bars = 400 μm. **b,** Representative day 30 organoid immunofluorescence images with DAPI (blue), TNNT2 (red), and PECAM1 (green) (n=7-8 organoids per condition). Images presented as maximum intensity projections. Scale bars = 400 μm. **c,** Quantification of PECAM1+ area presented in **(b)** (n=7-8 organoids per condition). Values = mean ± s.e.m. One-way ANOVA with Dunnett’s multiple comparisons test.

**Extended Data Fig. 5.**
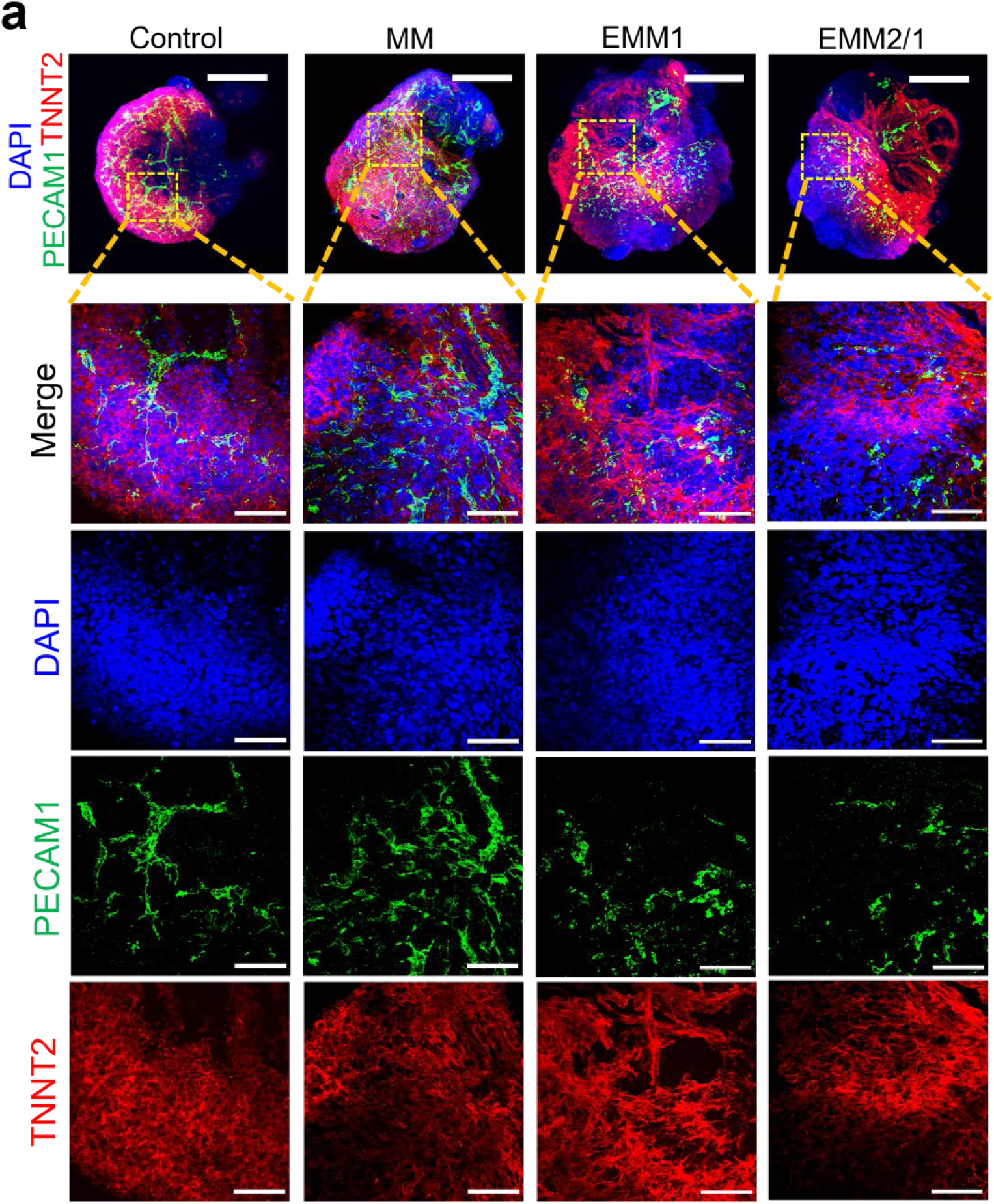
High magnification confocal microscopy reveals endothelial cell interconnectivity. **a,** Representative high magnification maximum intensity projection immunofluorescence images of organoids in each condition with DAPI (blue), TNNT2 (red), and PECAM1 (green) (n=7-8 organoids per condition). Scale bars = 50 μm. The images on top are representative low magnification organoids (Scale bar = 400 μm) for each condition with the yellow square representing area of high magnification.

