## Supplementary Video Legends for "A patterned human heart tube organoid model generated by pluripotent stem cell self-assembly"

**Title: Supplementary Video 1**

Description: Live imaging of representative Day 30 Control organoid at 10X magnification under brightfield microscopy.

**Title: Supplementary Video 2**

Description: Live imaging of representative Day 30 MM organoid at 10X magnification under brightfield microscopy.

**Title: Supplementary Video 3**

Description: Live imaging of representative Day 30 EMM1 organoid at 10X magnification under brightfield microscopy.

**Title: Supplementary Video 4**

Description: Live imaging of representative Day 30 EMM2/1 organoid at 10X magnification under brightfield microscopy.

**Title: Supplementary Video 5**

Description: Live imaging of calcium transients at 100X magnification within a representative Day 30 Control organoid displays robust beating and calcium activity.

**Title: Supplementary Video 6**

Description: Live imaging of calcium transients at 100X magnification within a representative Day 30 MM organoid displays robust beating and calcium activity.

**Title: Supplementary Video 7**

Description: Live imaging of calcium transients at 100X magnification within a representative Day 30 EMM1 organoid displays robust beating and calcium activity.

**Title: Supplementary Video 8**

Description: Live imaging of calcium transients at 100X magnification within a representative Day 30 EMM2/1 organoid displays robust beating and calcium activity.

**Title: Supplementary Video 9**

Description: 3D reconstruction of confocal immunofluorescence Z-series images of a Day 30 Control organoid stained for DAPI (blue), PECAM1 (green) and TNNT2 (red) showing an internal vascular network. Scale bar = 200 µm.

**Title: Supplementary Video 10**

Description: 3D reconstruction of confocal immunofluorescence Z-series images of a Day 30 MM organoid stained for DAPI (blue), PECAM1 (green) and TNNT2 (red) showing an internal vascular network. Scale bar = 200 µm.

**Title: Supplementary Video 11**

Description: 3D reconstruction of confocal immunofluorescence Z-series images of a Day 30 EMM1 organoid stained for DAPI (blue), PECAM1 (green) and TNNT2 (red) showing an internal vascular network. Scale bar = 200 µm.

**Title: Supplementary Video 12**

Description: 3D reconstruction of confocal immunofluorescence Z-series images of a Day 30 EMM2/1 organoid stained for DAPI (blue), PECAM1 (green) and TNNT2 (red) showing an internal vascular network. Scale bar = 200 µm.

**Title: Supplementary Video 13**

Description: 3D OCT cross-sectional scan of Day 30 Control organoid showing complex, internal chamber morphology and interconnectivity. Scale bar = 400 μm.

**Title: Supplementary Video 14**

Description: 3D OCT cross-sectional scan of Day 30 MM organoid showing complex, internal chamber morphology and interconnectivity. Scale bar = 400 μm.

**Title: Supplementary Video 15**

Description: 3D OCT cross-sectional scan of Day 30 EMM1 organoid showing complex, internal chamber morphology and interconnectivity. Scale bar = 400 μm.

**Title: Supplementary Video 16**

Description: 3D OCT cross-sectional scan of Day 30 EMM2/1 organoid showing complex, internal chamber morphology and interconnectivity. Scale bar = 400 μm.
