## Supplementary Information for "A patterned human heart tube organoid model generated by pluripotent stem cell self-assembly"

**Contents**

**Supplementary Table 1. |** Antibodies used for immunofluorescence.

**Supplementary Fig. 1.** Spatial expression of cardiomyocyte-related genes within single cell RNA sequencing datasets for day 34 organoids in each condition. Data presented as UMAP projections of k-means 8 clustering.

**Supplementary Fig. 2.** Spatial expression of valve-related genes within single cell RNA sequencing datasets for day 34 organoids in each condition. Data presented as UMAP projections of k-means 8 clustering.

**Supplementary Fig. 3.** Spatial expression of epicardial-related genes within single cell RNA sequencing datasets for day 34 organoids in each condition. Data presented as UMAP projections of k-means 8 clustering.

**Supplementary Fig. 4.** Spatial expression of epicardial-related genes within single cell RNA sequencing datasets for day 34 organoids in each condition. Data presented as UMAP projections of k-means 8 clustering.

**Supplementary Fig. 5.** Spatial expression of stromal-related genes within single cell RNA sequencing datasets for day 34 organoids in each condition. Data presented as UMAP projections of k-means 8 clustering.

**Supplementary Fig. 6.** Spatial expression of conductance-related genes within single cell RNA sequencing datasets for day 34 organoids in each condition. Data presented as UMAP projections of k-means 8 clustering.

**Supplementary Fig. 7.** Spatial expression of endothelial-related genes within single cell RNA sequencing datasets for day 34 organoids in each condition. Data presented as UMAP projections of k-means 8 clustering.

**Supplementary Fig. 8.** Spatial expression of cardiac fibroblast-related genes within single cell RNA sequencing datasets for day 34 organoids in each condition. Data presented as UMAP projections of k-means 8 clustering.

**Supplementary Fig. 9.** Spatial expression of left-right asymmetry-related genes within single cell RNA sequencing datasets for day 34 organoids in each condition. Data presented as UMAP projections of k-means 8 clustering.

**Supplementary Fig. 10.** Spatial expression of proliferation-related genes within single cell RNA sequencing datasets for day 34 organoids in each condition. Data presented as UMAP projections of k-means 8 clustering.

**Supplementary Fig. 11.** Spatial expression of heart field-related genes within single cell RNA sequencing datasets for day 34 organoids in each condition. Data presented as UMAP projections of k-means 8 clustering.

**Supplementary Fig. 12.** Spatial expression of anterior second heart field (aSHF) genes within single cell RNA sequencing datasets for day 34 organoids in each condition. Data presented as UMAP projections of k-means 8 clustering.

**Supplementary Fig. 13.** Spatial expression of posterior second heart field (pSHF) genes within single cell RNA sequencing datasets for day 34 organoids in each condition. Data presented as UMAP projections of k-means 8 clustering.

**Supplementary Fig. 14.** Spatial expression of first heart field (FHF) genes within single cell RNA sequencing datasets for day 34 organoids in each condition. Data presented as UMAP projections of k-means 8 clustering.

**Supplementary Fig. 15.** Spatial expression of outflow tract (OFT) genes within single cell RNA sequencing datasets for day 34 organoids in each condition. Data presented as UMAP projections of k-means 8 clustering.

**Supplementary Fig. 16.** Gene ontologies (GO) for differentially expressed genes within the VCM, ACM, VC, and CC clusters for each maturation condition. Common differentially expressed genes shared amongst all four maturation conditions are displayed in the center of each cluster, and the top differentially expressed genes which contribute most to the GO for each condition are displayed in each corner, respectively.

**Supplementary Fig. 17.** Gene ontologies (GO) for differentially expressed genes within the PEDC, SC, and EPC clusters for each maturation condition. Common differentially expressed genes shared amongst all four maturation conditions are displayed in the center of each cluster, and the top differentially expressed genes which contribute most to the GO for each condition are displayed in each corner, respectively.

**Supplementary Fig. 18.** Cell-cell communication via ligand-receptor pairing for each cluster from scRNAseq data from the control condition.

**Supplementary Fig. 19.** Cell-cell communication via ligand-receptor pairing for each cluster from scRNAseq data from the MM condition.

**Supplementary Fig. 20.** Cell-cell communication via ligand-receptor pairing for each cluster from scRNAseq data from the EMM1 condition.

**Supplementary Fig. 21.** Cell-cell communication via ligand-receptor pairing for each cluster from scRNAseq data from the EMM2/1 condition.

**Supplementary Fig. 22.** KEGG map of glycolysis and gluconeogenesis enzyme pathway rendered by Pathview. Log2-ratio calculated using scRNAseq data from the Control condition.

**Supplementary Fig. 23.** KEGG map of glycolysis and gluconeogenesis enzyme pathway rendered by Pathview. Log2-ratio calculated using scRNAseq data from the MM condition.

**Supplementary Fig. 24.** KEGG map of glycolysis and gluconeogenesis enzyme pathway rendered by Pathview. Log2-ratio calculated using scRNAseq data from the EMM1 condition.

**Supplementary Fig. 25.** KEGG map of glycolysis and gluconeogenesis enzyme pathway rendered by Pathview. Log2-ratio calculated using scRNAseq data from the EMM2/1 condition.

**Supplementary Fig. 26.** KEGG map of oxidative phosphorylation enzyme pathway rendered by Pathview. Log2-ratio calculated using scRNAseq data from the Control condition.

**Supplementary Fig. 27.** KEGG map of oxidative phosphorylation enzyme pathway rendered by Pathview. Log2-ratio calculated using scRNAseq data from the MM condition.

**Supplementary Fig. 28.** KEGG map of oxidative phosphorylation enzyme pathway rendered by Pathview. Log2-ratio calculated using scRNAseq data from the EMM1 condition.

**Supplementary Fig. 29.** KEGG map of oxidative phosphorylation enzyme pathway rendered by Pathview. Log2-ratio calculated using scRNAseq data from the EMM2/1 condition.

**Supplementary Fig. 30.** Spatial expression of β-adrenoreceptor genes within single cell RNA sequencing datasets for day 34 organoids in each condition. Data presented as UMAP projections of k-means 8 clustering.

**Supplementary Fig. 31.** Schematic diagram for the Renishaw confocal Raman spectrometer for use with human heart organoids.

|  | **Antibody Name** | **Host Species** | **Dilution** | **Catalogue Number** | **Vendor** |
| --- | --- | --- | --- | --- | --- |
| **Primary** | TNNT2 | Mouse | 1:200 | ab8295 | Abcam |
|  | TNNT2 | Rabbit | 1:200 | ab45932 | Abcam |
|  | WT1 | Rabbit | 1:200 | ab89901 | Abcam |
|  | PECAM1 | Mouse | 1:50 | P2B1 | DSHB |
|  | MYL2 | Rabbit | 1:200 | ab79935 | Abcam |
|  | MYL7 | Mouse | 1:200 | 311-011 | Synaptic Systems |
|  | TBX18 | Mouse | 1:200 | ab201587 | Abcam |
|  | ALDH1A2 | Rabbit | 1:200 | ABN420 | Sigma |
|  | KCNJ2 | Rabbit | 1:500 | HPA029109 | Sigma |
| **Secondary** | Alexa Fluor 488 | Donkey anti-mouse | 1:200 | A-21202 | Thermo Fisher Scientific |
|  | Alexa Fluor 488 | Donkey anti-rabbit | 1:200 | A-21206 | Thermo Fisher Scientific |
|  | Alexa Fluor 594 | Donkey anti-mouse | 1:200 | A-21203 | Thermo Fisher Scientific |
|  | Alexa Fluor 594 | Donkey anti-rabbit | 1:200 | A-21207 | Thermo Fisher Scientific |

**Supplementary Table 1.** Antibodies used for immunofluorescence.

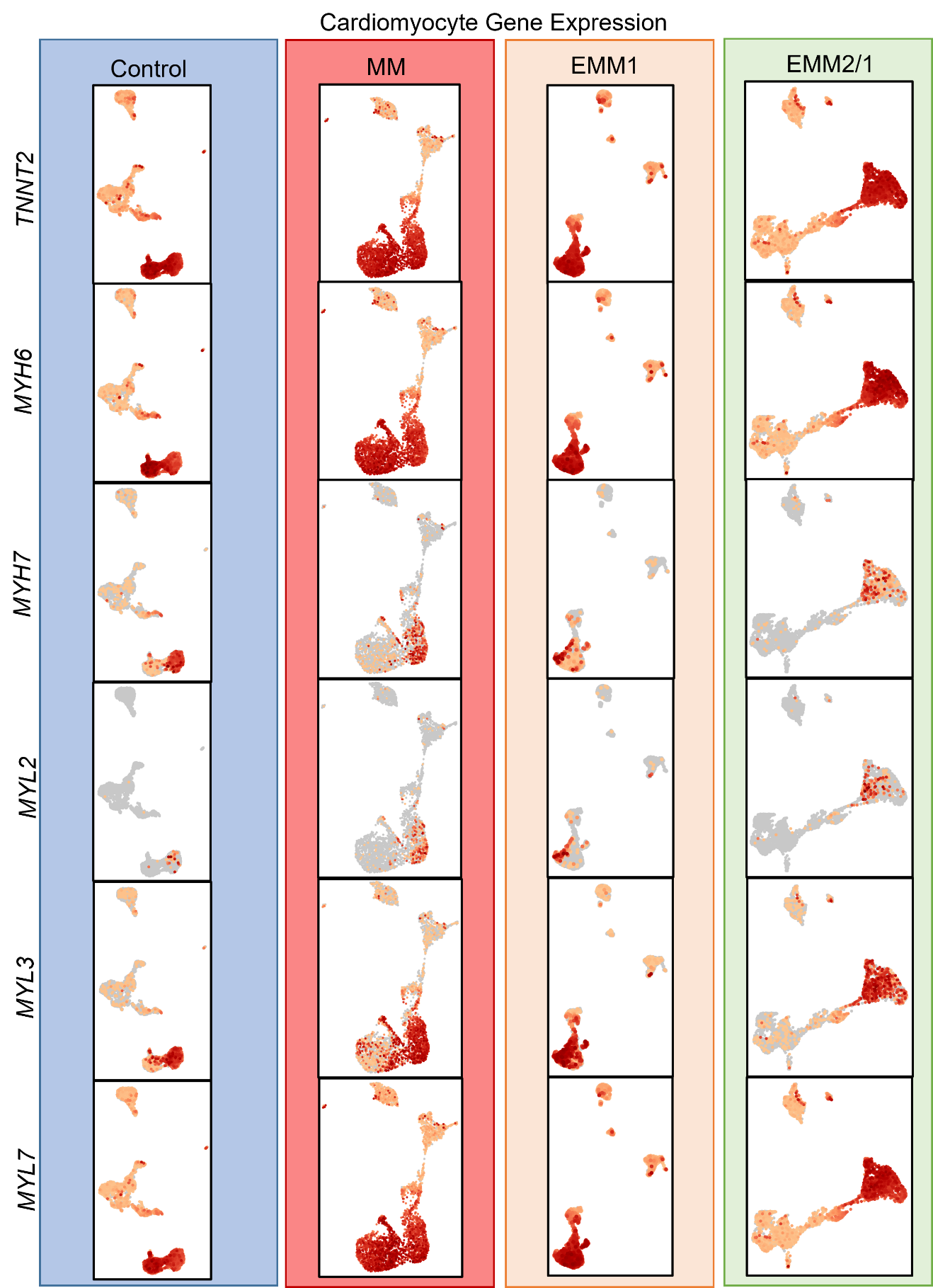

**Supplementary Fig. 1.** Spatial expression of cardiomyocyte-related genes within single cell RNA sequencing datasets for day 34 organoids in each condition. Data presented as UMAP projections of k-means 8 clustering.

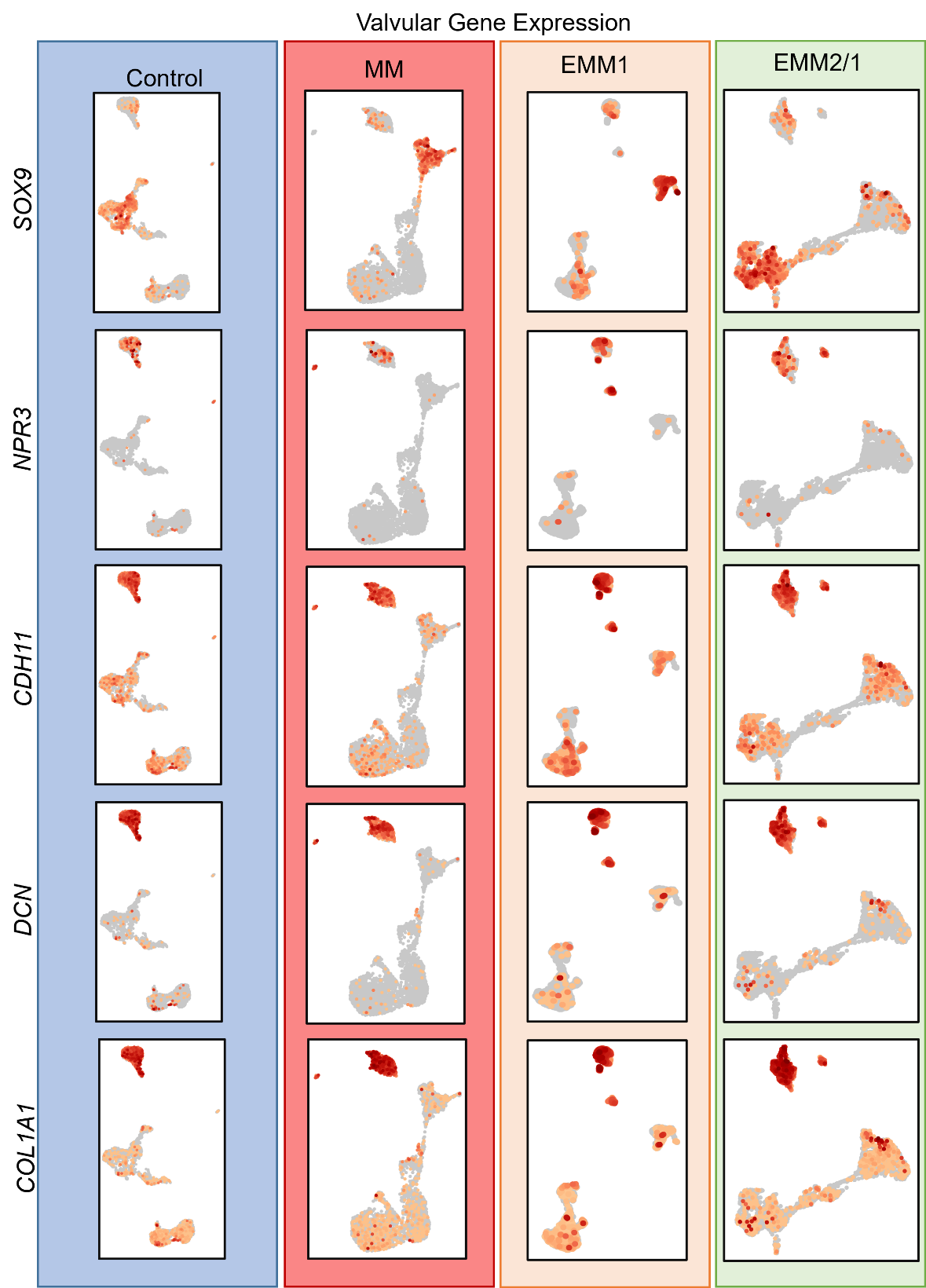

**Supplementary Fig. 2.** Spatial expression of valve-related genes within single cell RNA sequencing datasets for day 34 organoids in each condition. Data presented as UMAP projections of k-means 8 clustering.

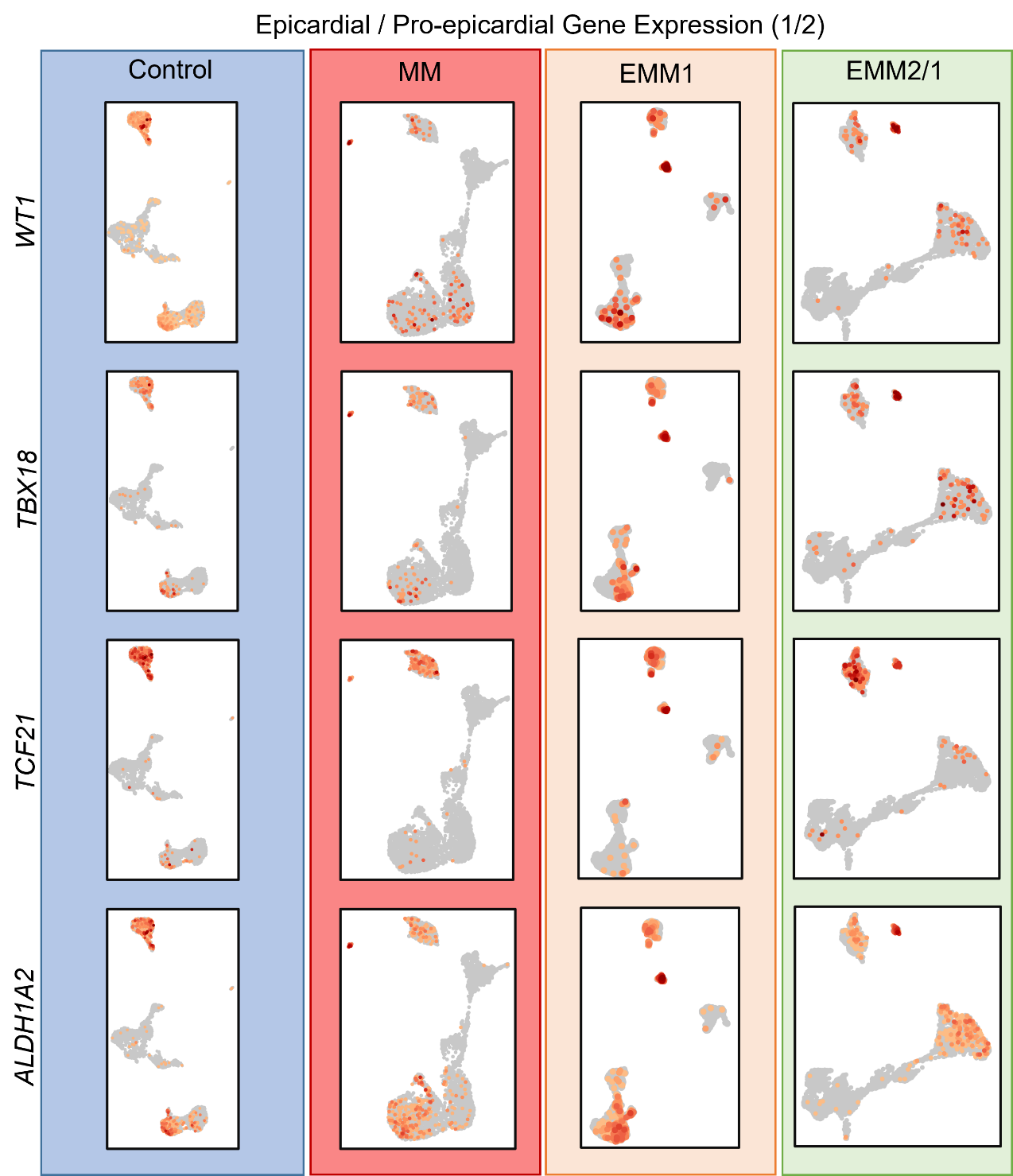

**Supplementary Fig. 3.** Spatial expression of epicardial-related genes within single cell RNA sequencing datasets for day 34 organoids in each condition. Data presented as UMAP projections of k-means 8 clustering.

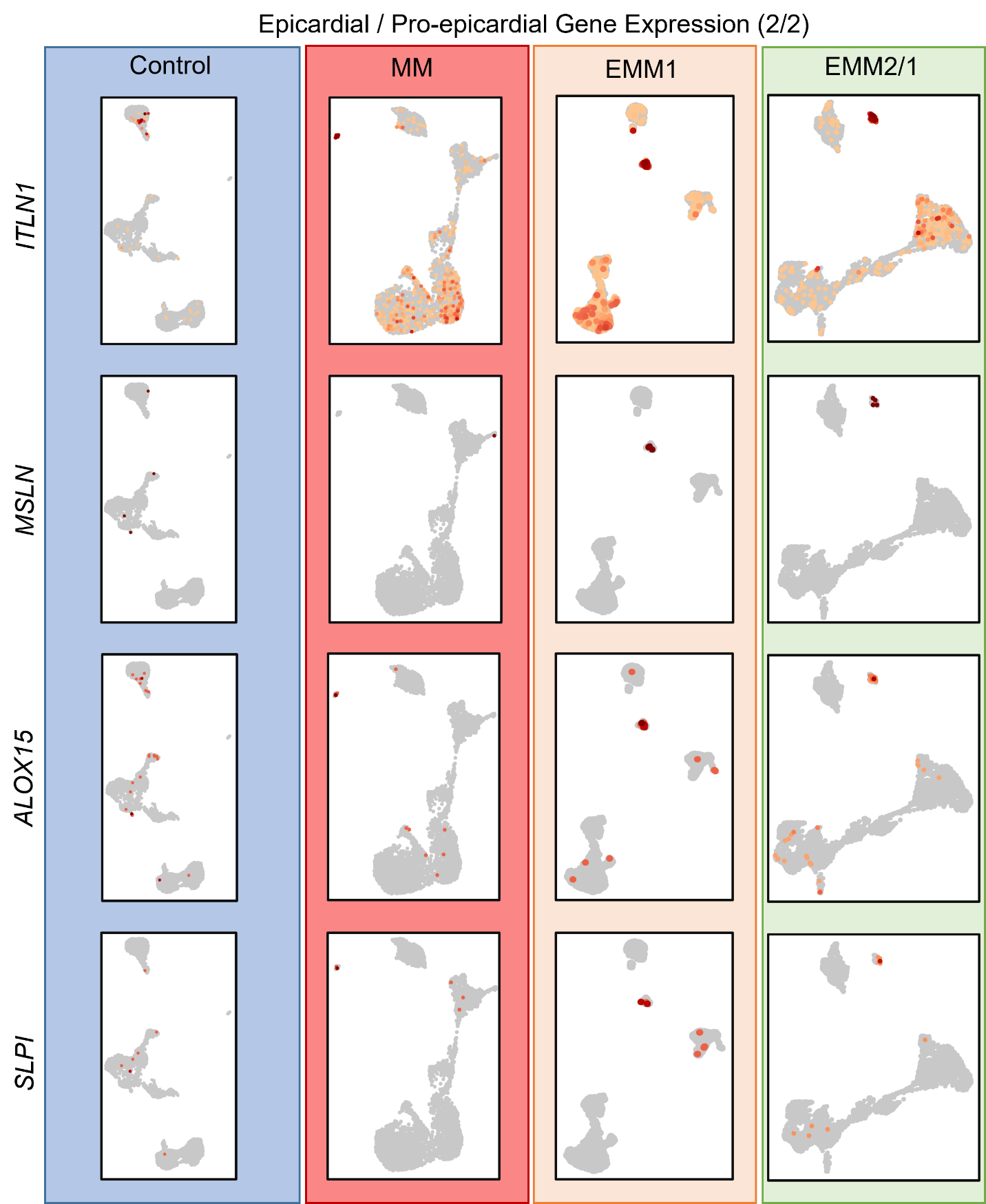

**Supplementary Fig. 4.** Spatial expression of epicardial-related genes within single cell RNA sequencing datasets for day 34 organoids in each condition. Data presented as UMAP projections of k-means 8 clustering.

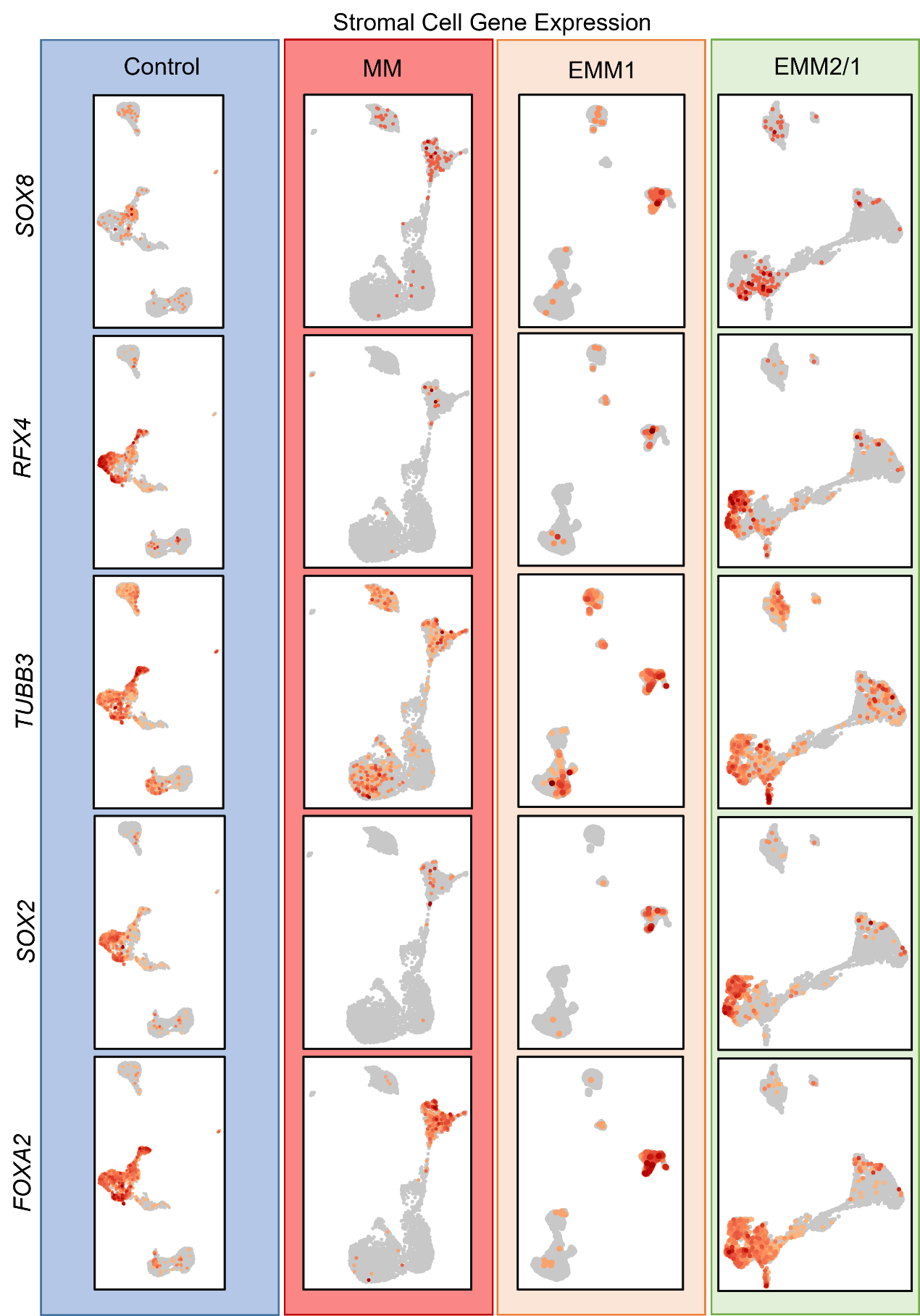

**Supplementary Fig. 5.** Spatial expression of stromal-related genes within single cell RNA sequencing datasets for day 34 organoids in each condition. Data presented as UMAP projections of k-means 8 clustering.

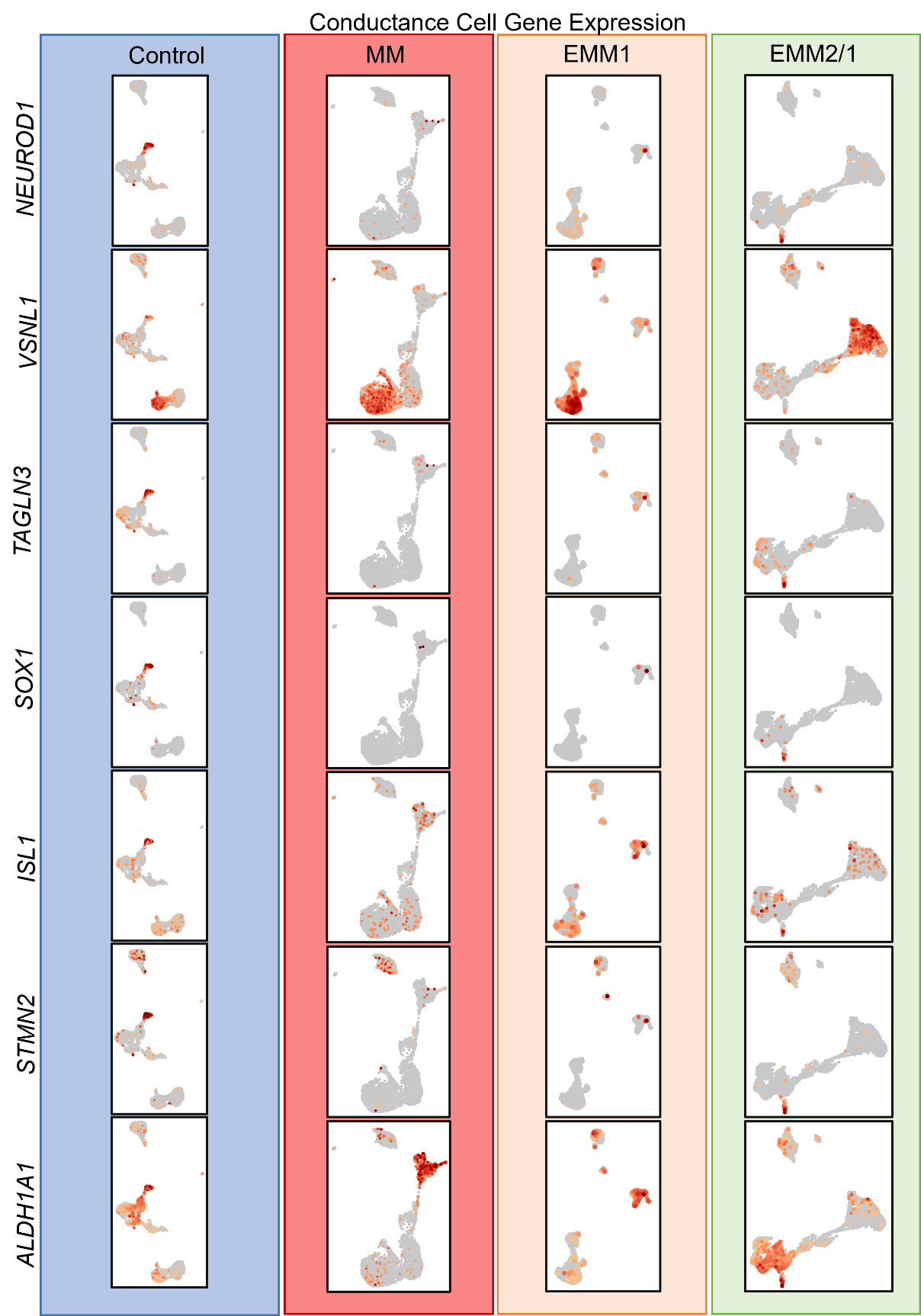

**Supplementary Fig. 6.** Spatial expression of conductance-related genes within single cell RNA sequencing datasets for day 34 organoids in each condition. Data presented as UMAP projections of k-means 8 clustering.

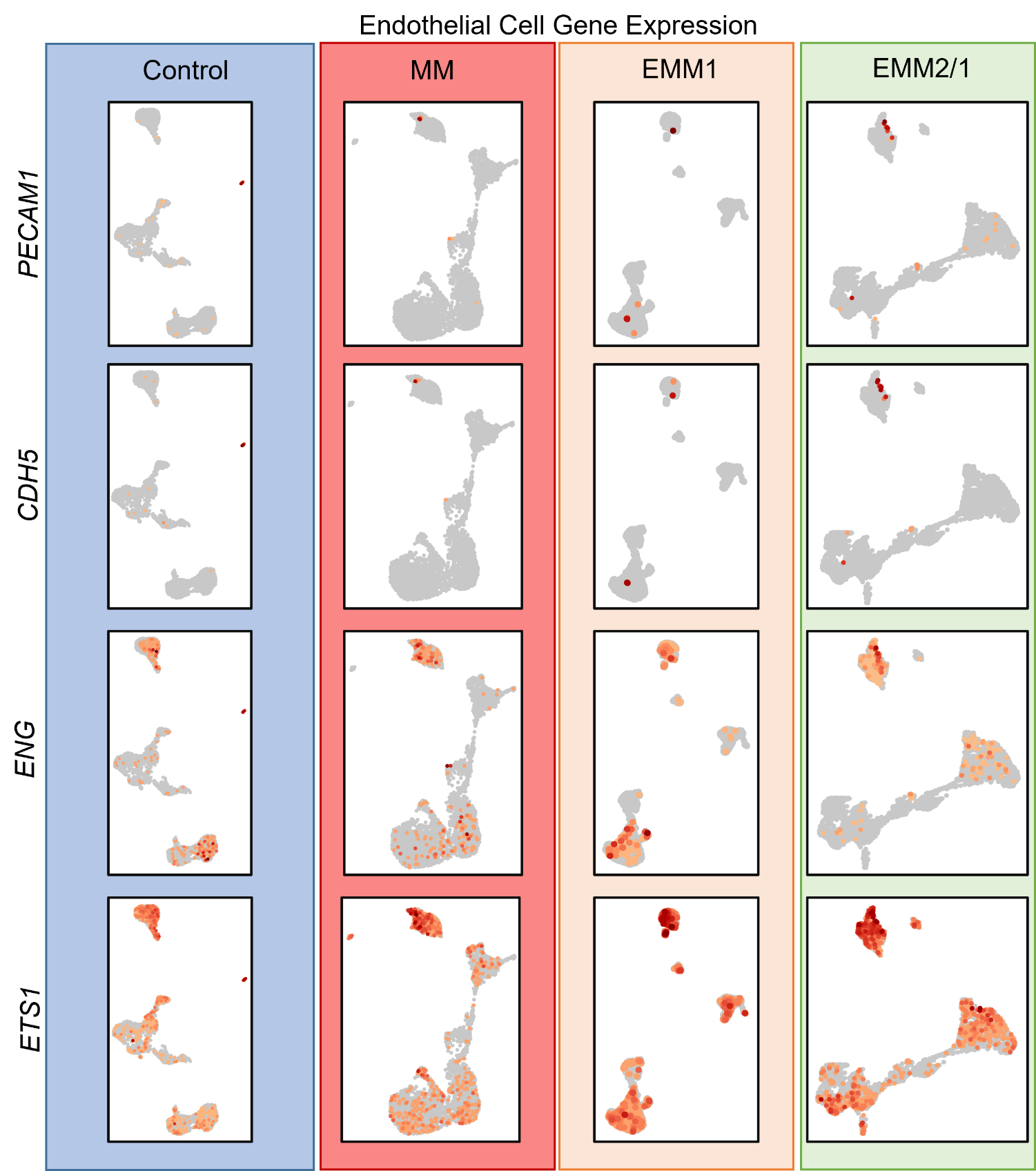

**Supplementary Fig. 7.** Spatial expression of endothelial-related genes within single cell RNA sequencing datasets for day 34 organoids in each condition. Data presented as UMAP projections of k-means 8 clustering.

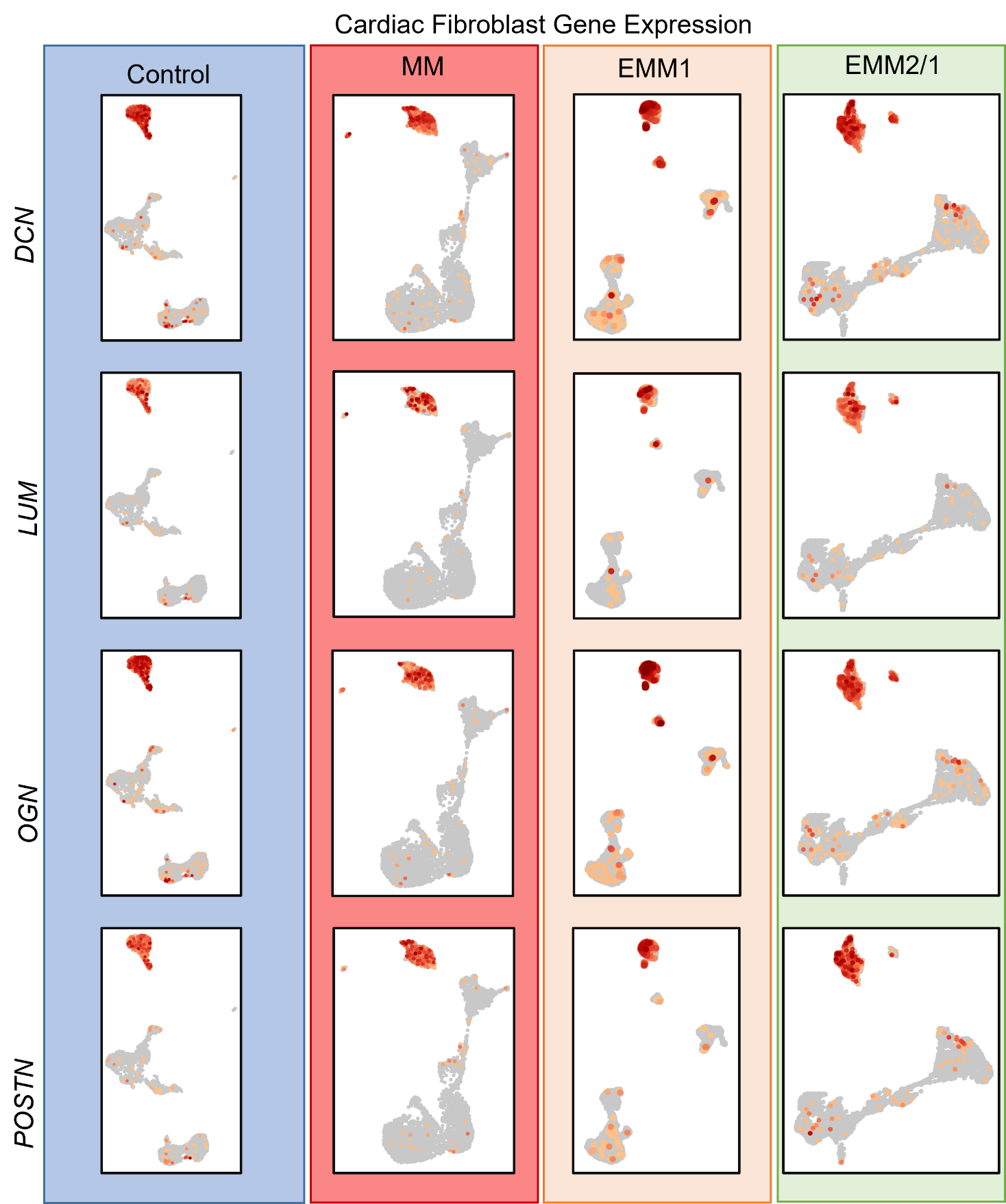

**Supplementary Fig. 8.** Spatial expression of cardiac fibroblast-related genes within single cell RNA sequencing datasets for day 34 organoids in each condition. Data presented as UMAP projections of k-means 8 clustering.

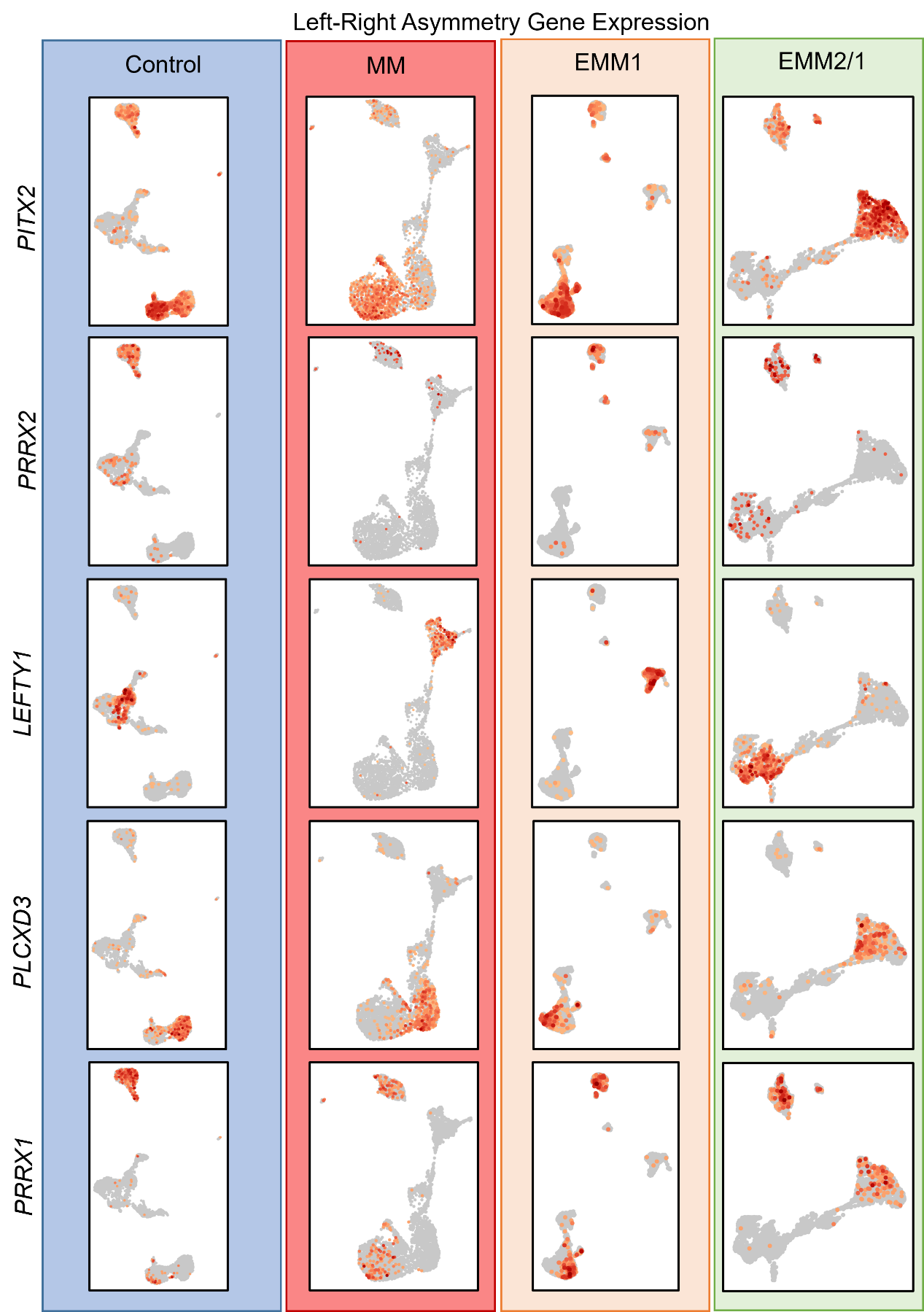

**Supplementary Fig. 9.** Spatial expression of left-right asymmetry-related genes within single cell RNA sequencing datasets for day 34 organoids in each condition. Data presented as UMAP projections of k-means 8 clustering.

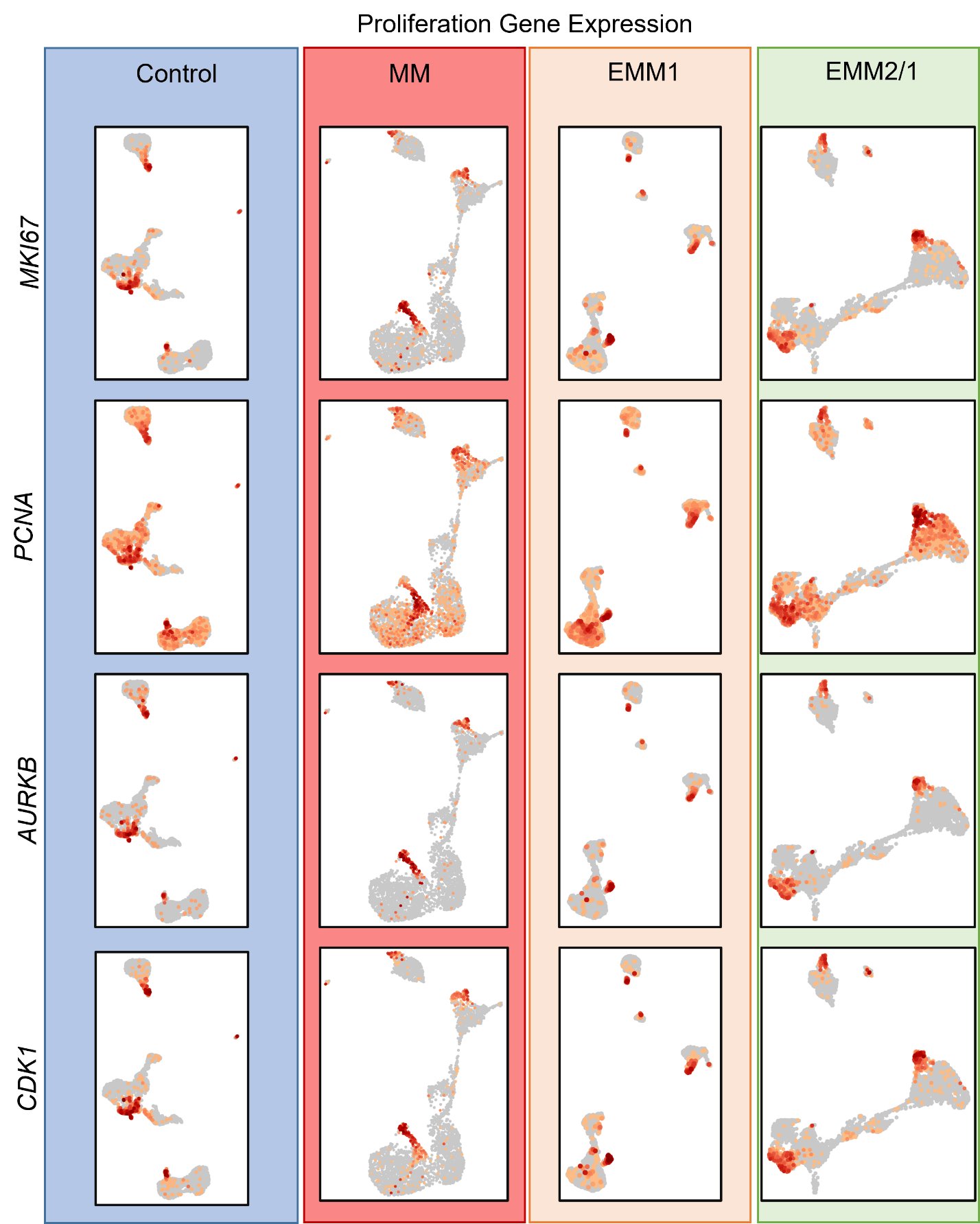

**Supplementary Fig. 10.** Spatial expression of proliferation-related genes within single cell RNA sequencing datasets for day 34 organoids in each condition. Data presented as UMAP projections of k-means 8 clustering.

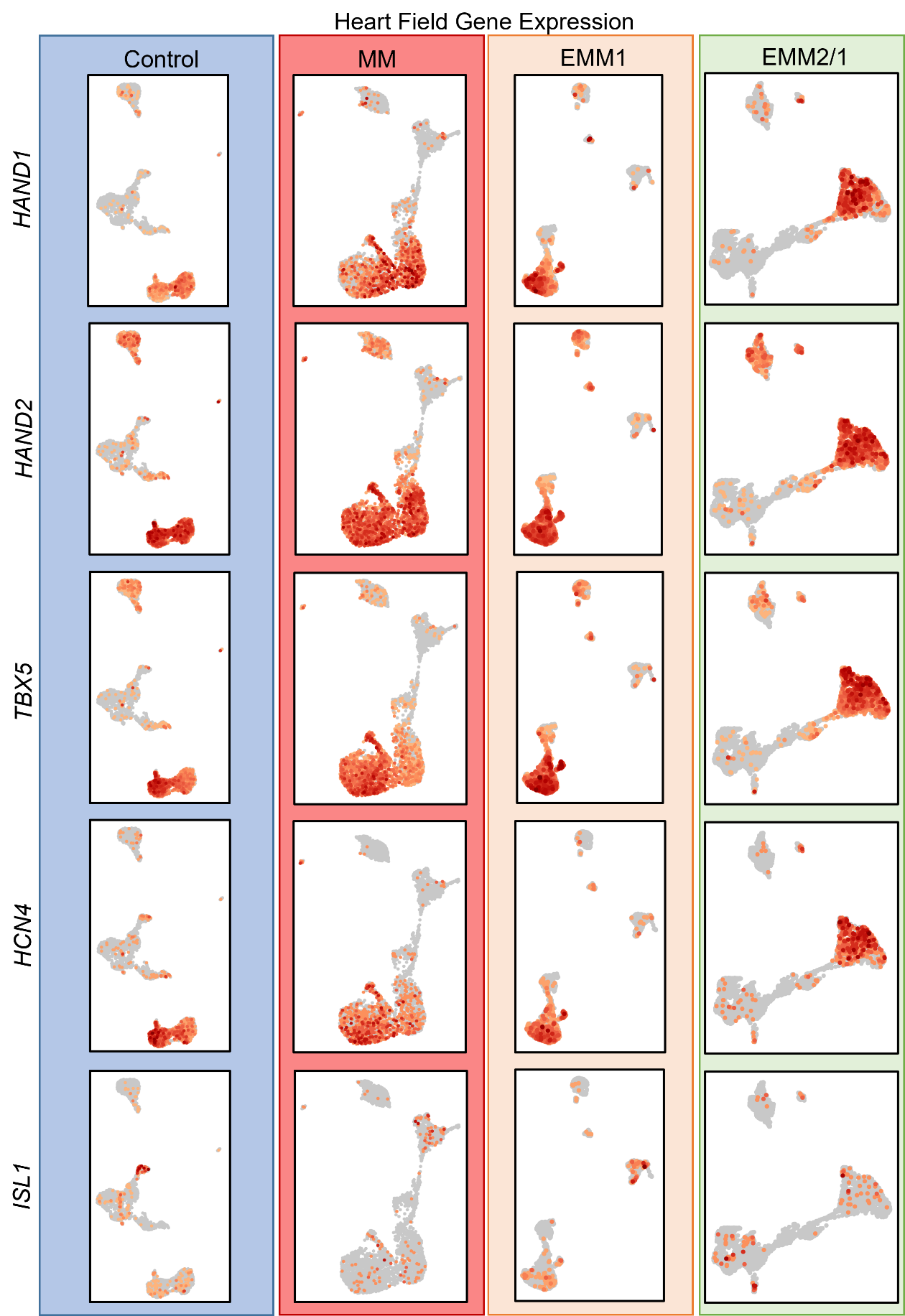

**Supplementary Fig. 11.** Spatial expression of heart field-related genes within single cell RNA sequencing datasets for day 34 organoids in each condition. Data presented as UMAP projections of k-means 8 clustering.

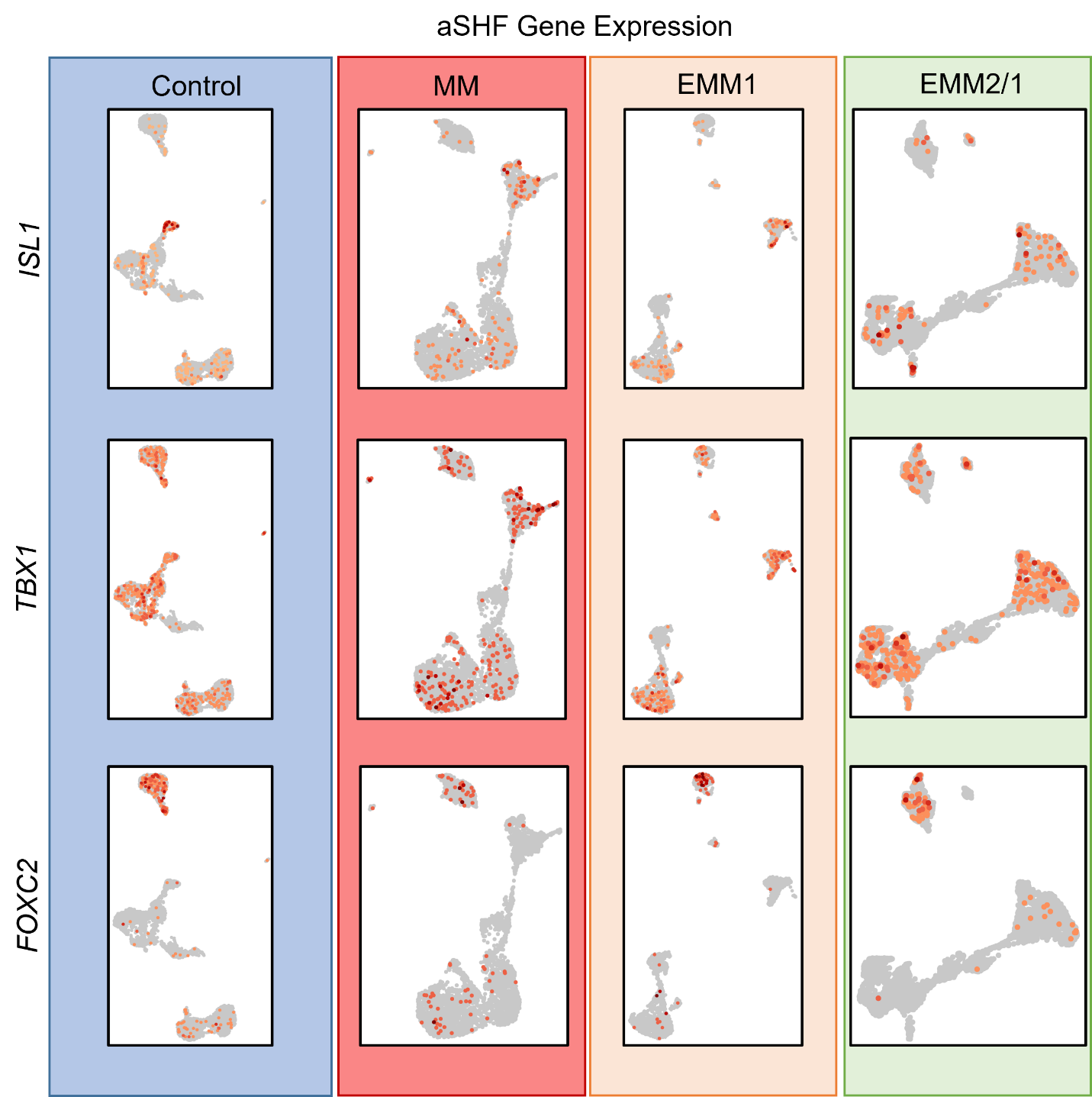

**Supplementary Fig. 12.** Spatial expression of anterior second heart field (aSHF) genes within single cell RNA sequencing datasets for day 34 organoids in each condition. Data presented as UMAP projections of k-means 8 clustering.

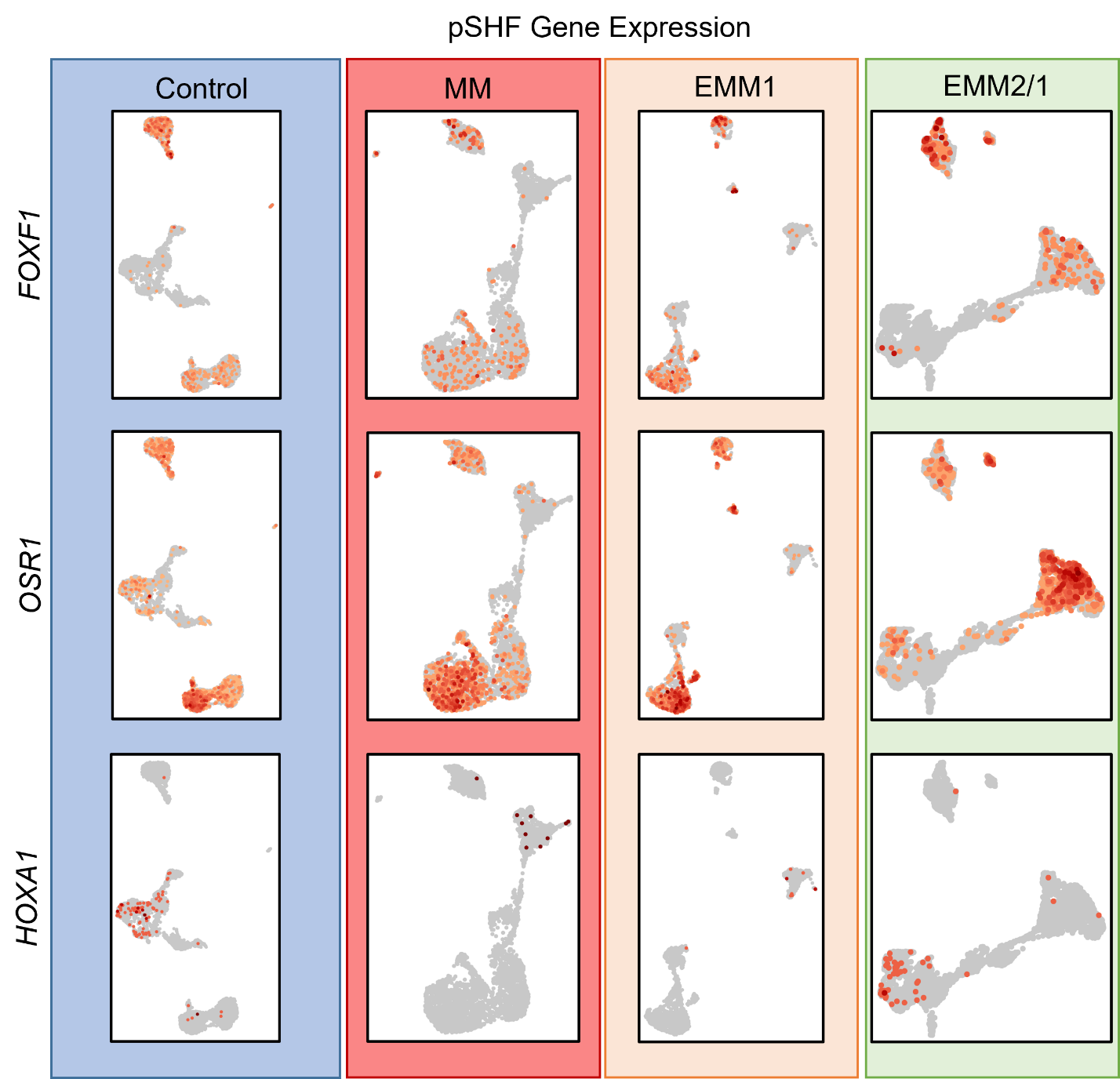

**Supplementary Fig. 13.** Spatial expression of posterior second heart field (pSHF) genes within single cell RNA sequencing datasets for day 34 organoids in each condition. Data presented as UMAP projections of k-means 8 clustering.

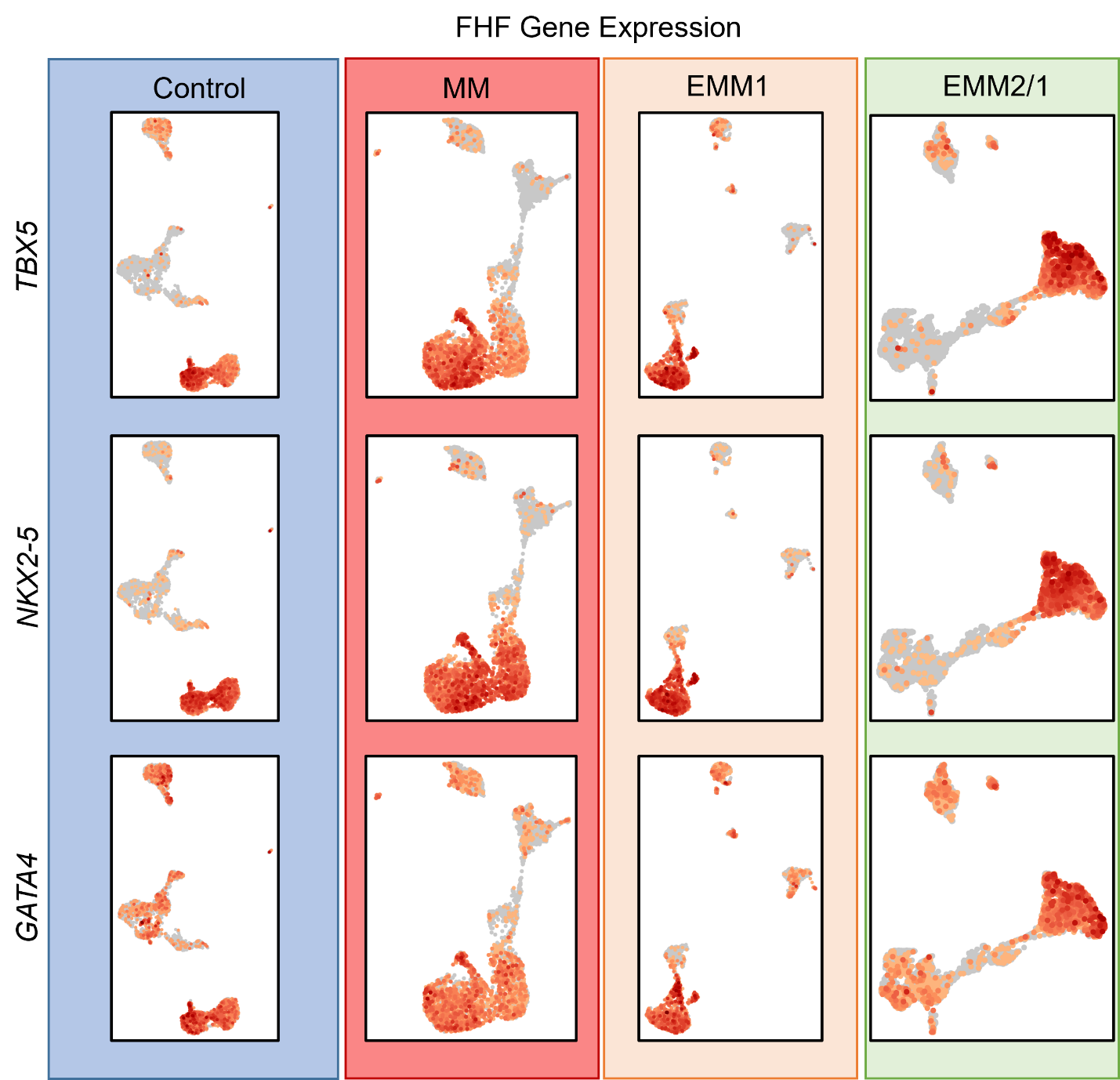

**Supplementary Fig. 14.** Spatial expression of first heart field (FHF) genes within single cell RNA sequencing datasets for day 34 organoids in each condition. Data presented as UMAP projections of k-means 8 clustering.

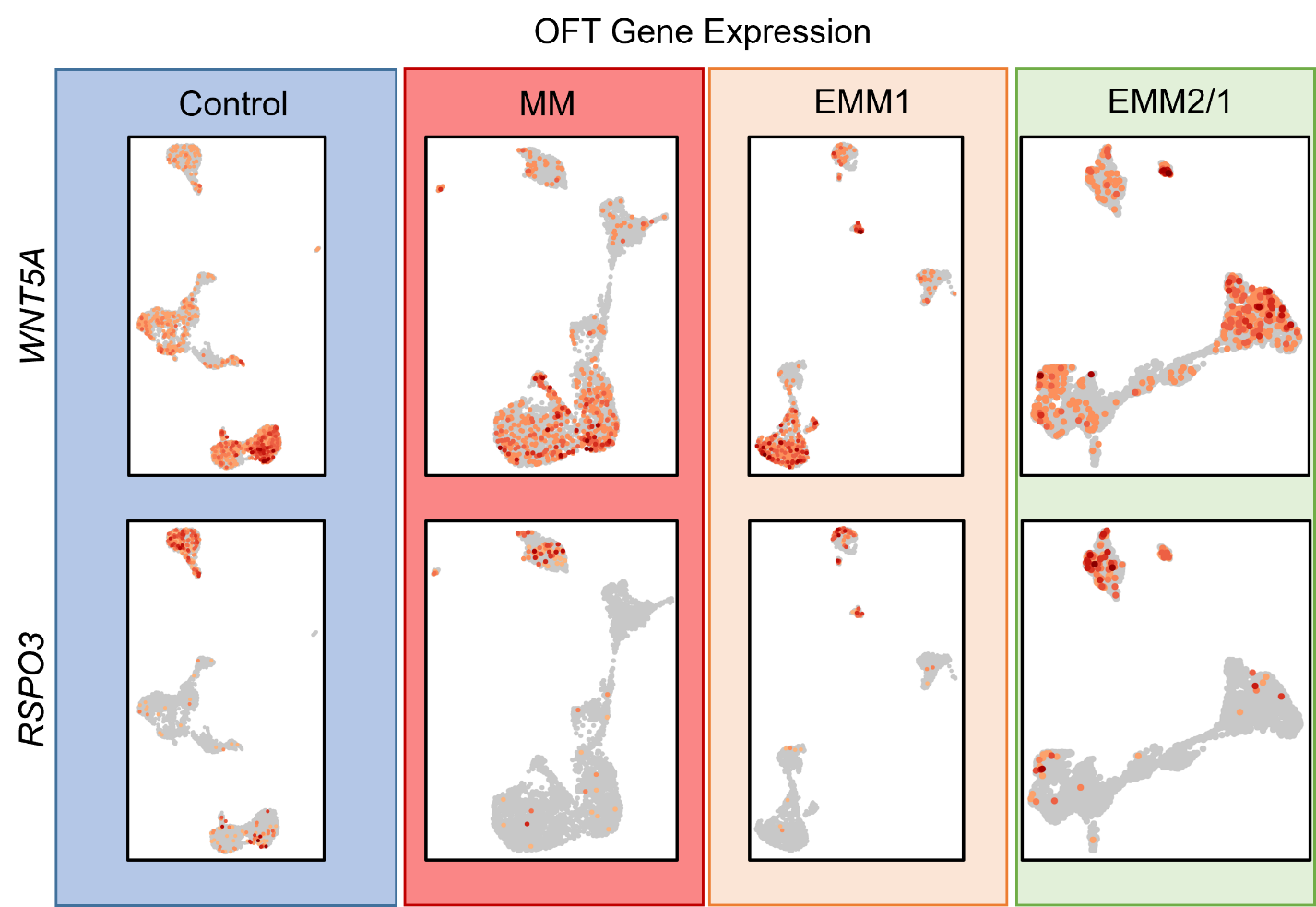

**Supplementary Fig. 15.** Spatial expression of outflow tract (OFT) genes within single cell RNA sequencing datasets for day 34 organoids in each condition. Data presented as UMAP projections of k-means 8 clustering.

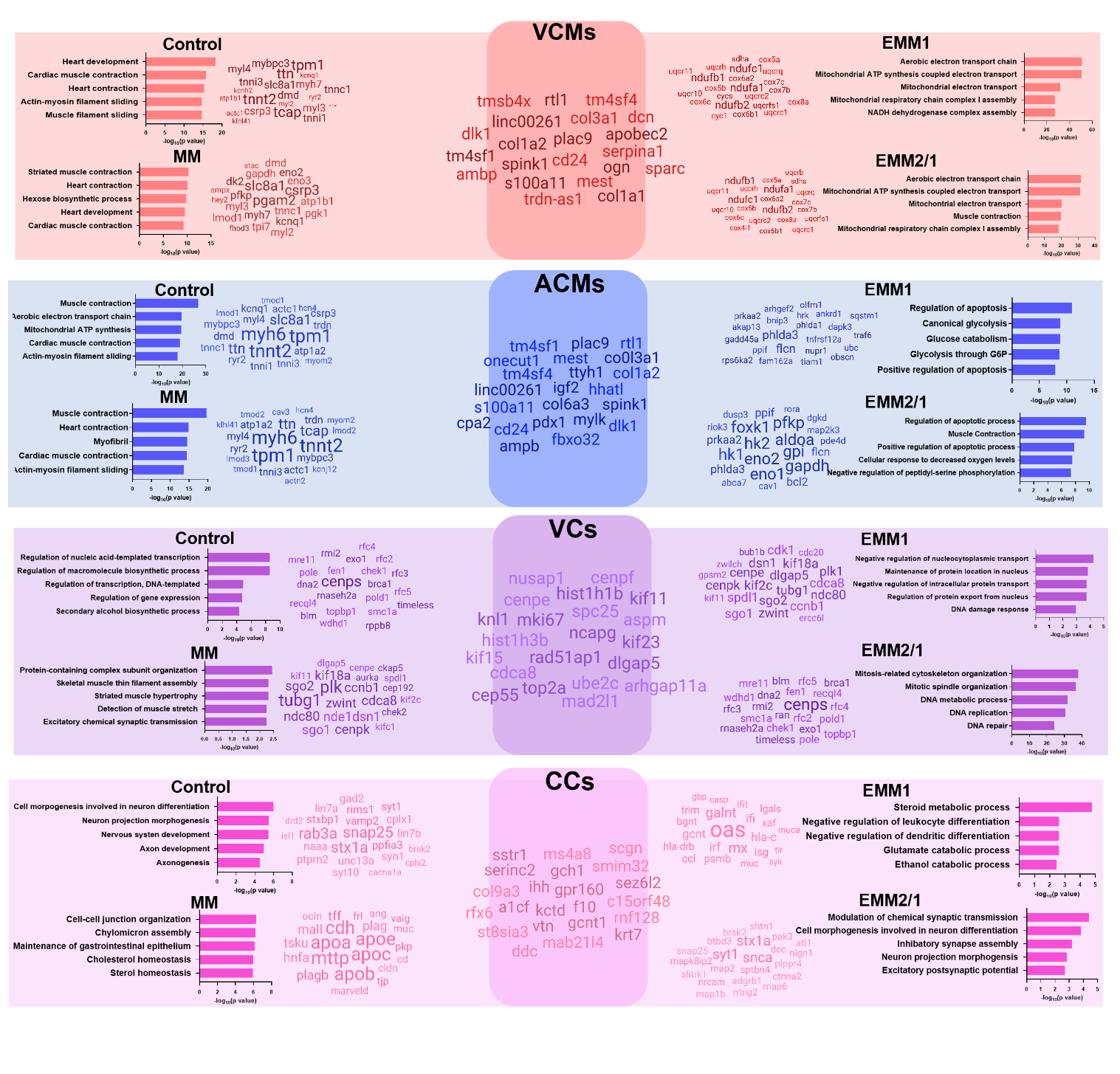
**Supplementary Fig. 16.** Gene ontologies (GO) for differentially expressed genes within the VCM, ACM, VC, and CC clusters for each maturation condition. Common differentially expressed genes shared amongst all four maturation conditions are displayed in the center of each cluster, and the top differentially expressed genes which contribute most to the GO for each condition are displayed in each corner, respectively.

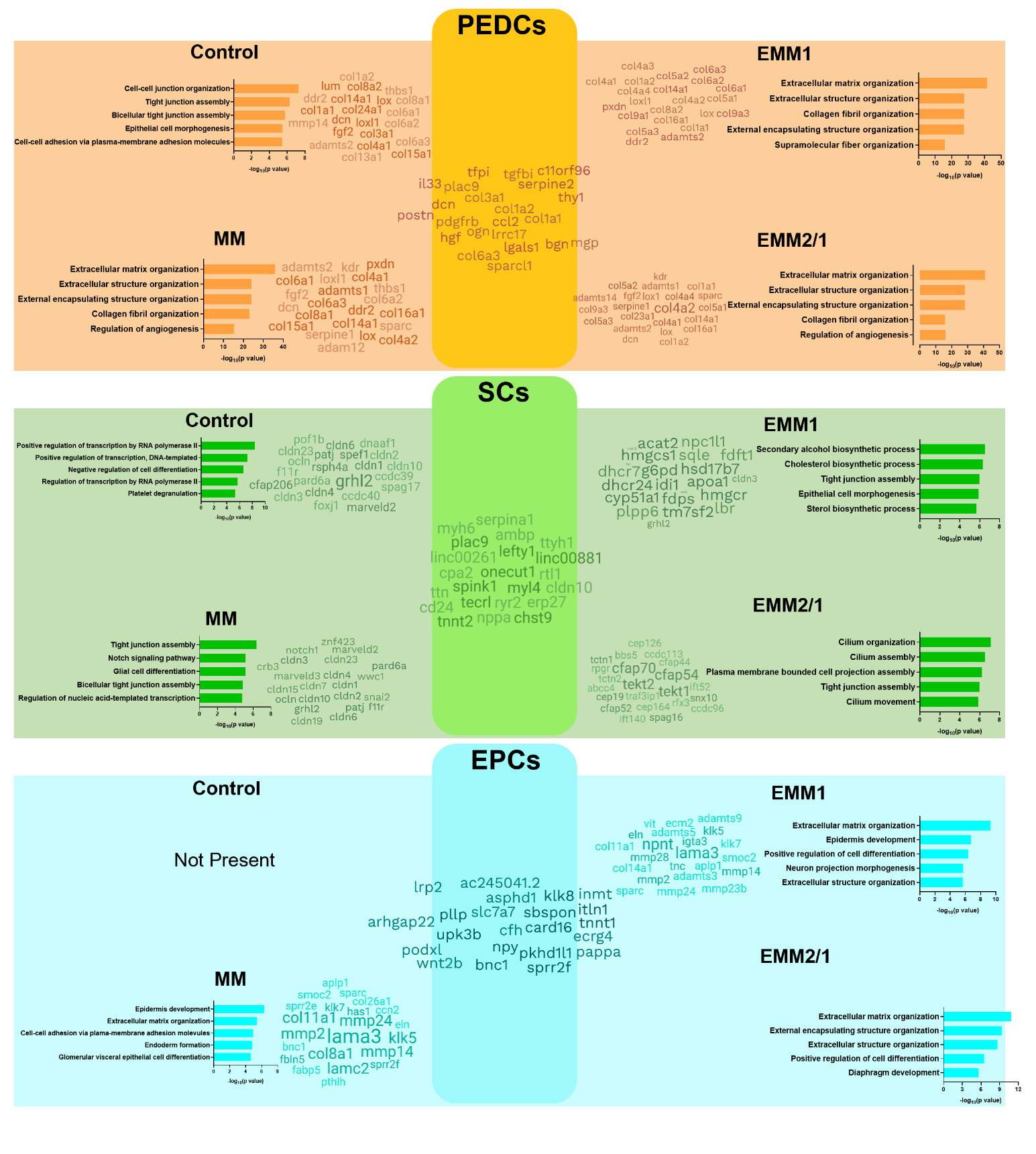
**Supplementary Fig. 17.** Gene ontologies (GO) for differentially expressed genes within the PEDC, SC, and EPC clusters for each maturation condition. Common differentially expressed genes shared amongst all four maturation conditions are displayed in the center of each cluster, and the top differentially expressed genes which contribute most to the GO for each condition are displayed in each corner, respectively.

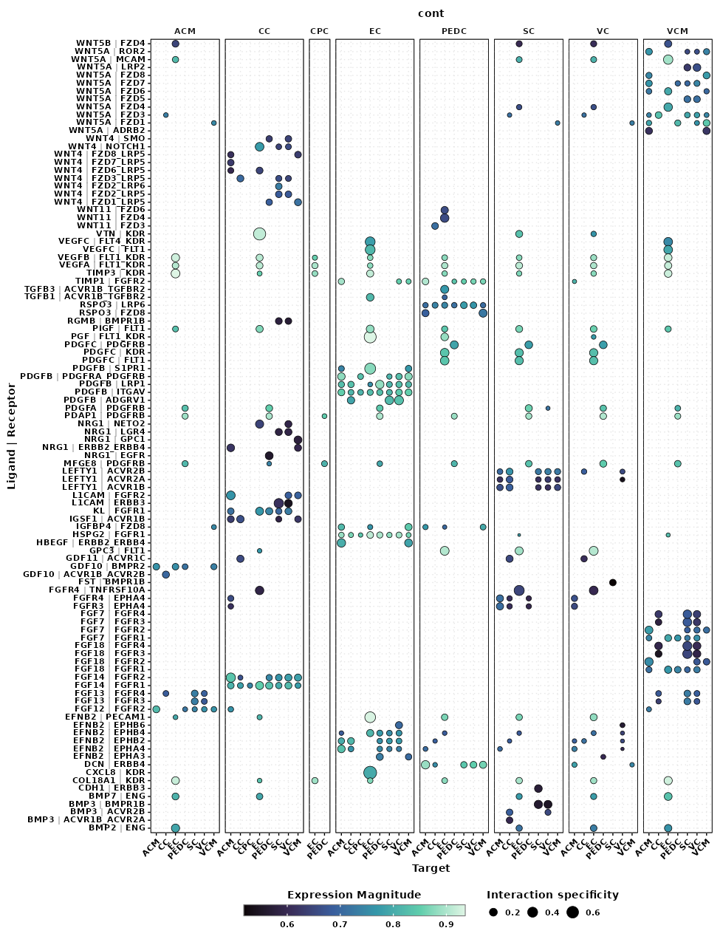

**Supplementary Fig. 18.** Cell-cell communication via ligand-receptor pairing for each cluster from scRNAseq data from the control condition.

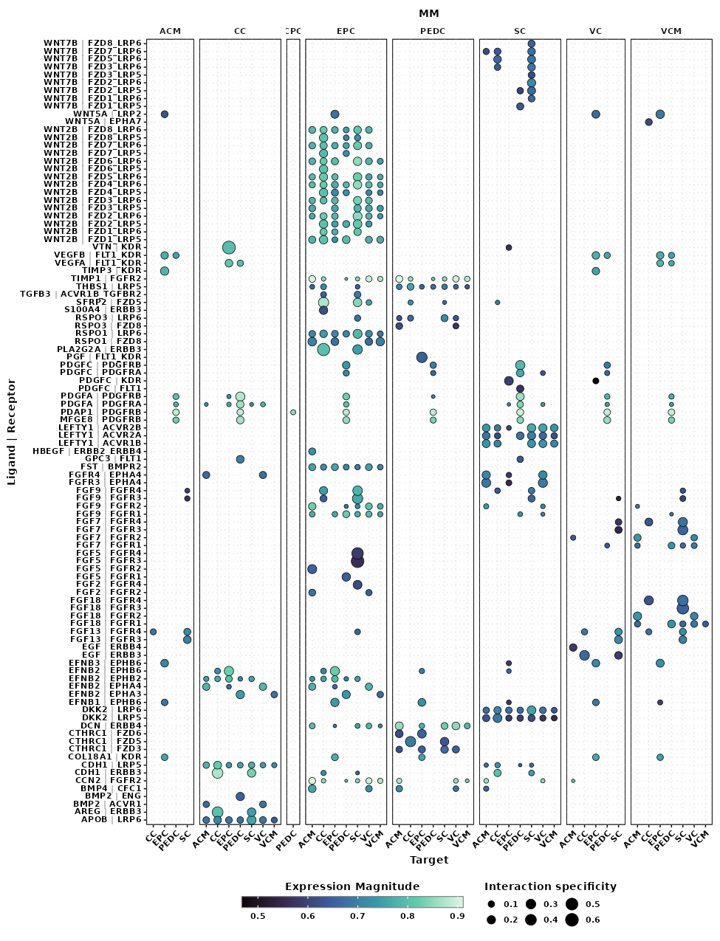

**Supplementary Fig. 19.** Cell-cell communication via ligand-receptor pairing for each cluster from scRNAseq data from the MM condition.

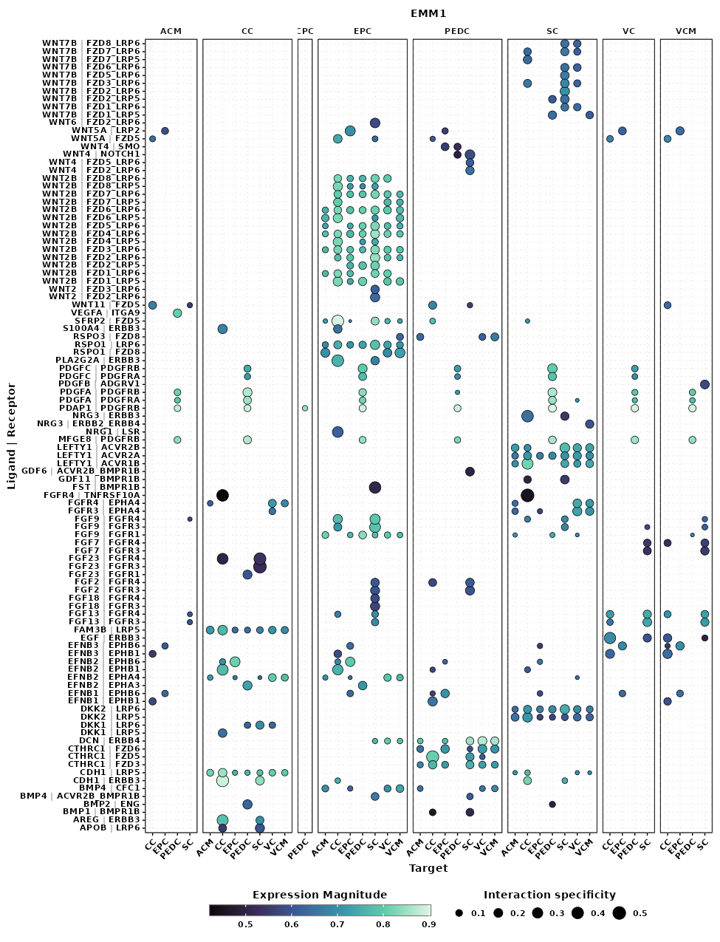

**Supplementary Fig. 20.** Cell-cell communication via ligand-receptor pairing for each cluster from scRNAseq data from the EMM1 condition.

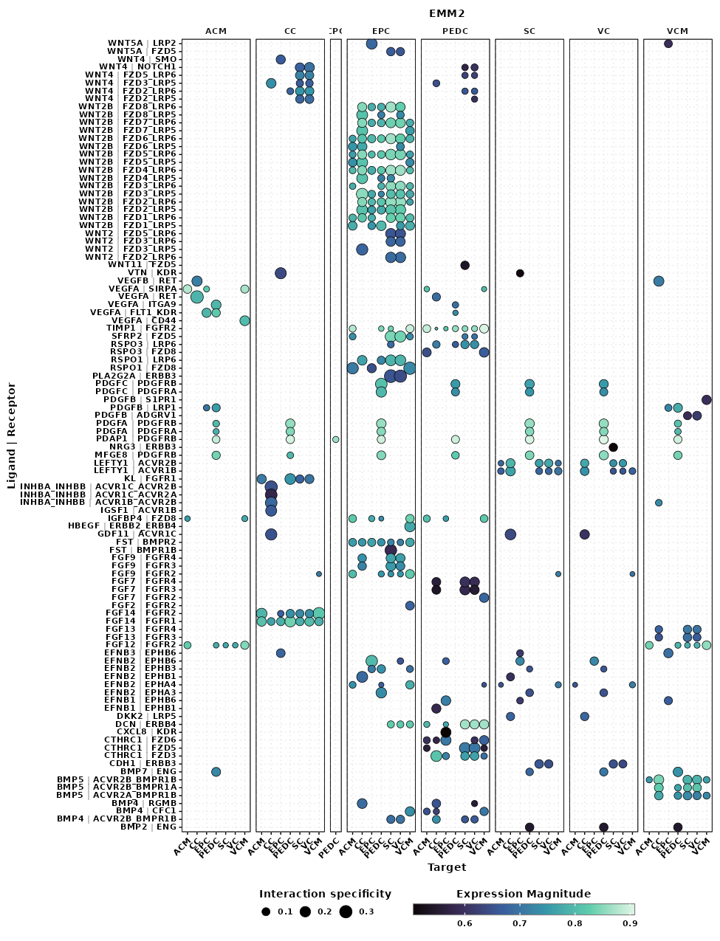

**Supplementary Fig. 21.** Cell-cell communication via ligand-receptor pairing for each cluster from scRNAseq data from the EMM2/1 condition.

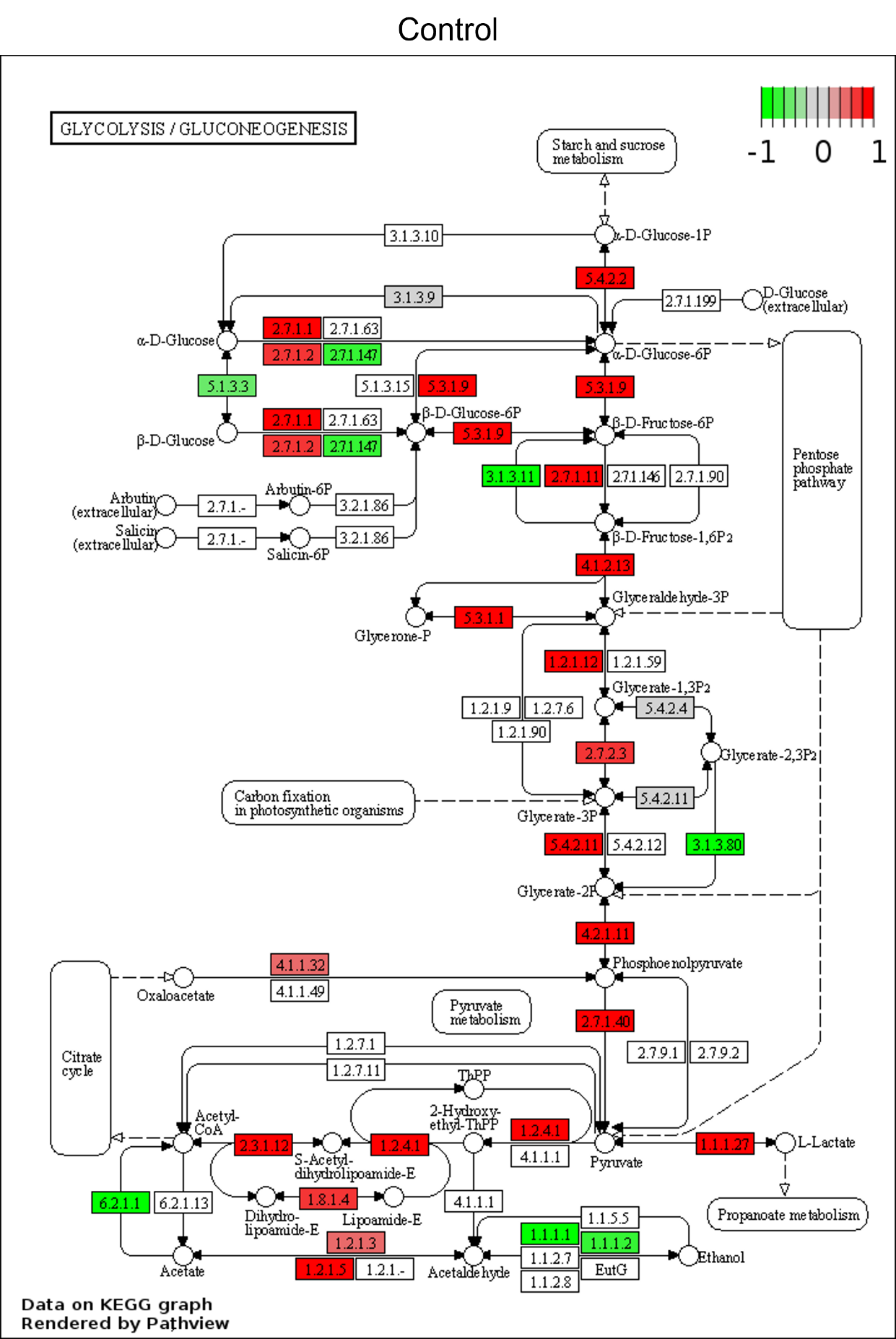

**Supplementary Fig. 22.** KEGG map of glycolysis and gluconeogenesis enzyme pathway rendered by Pathview. Log2-ratio calculated using scRNAseq data from the Control condition.

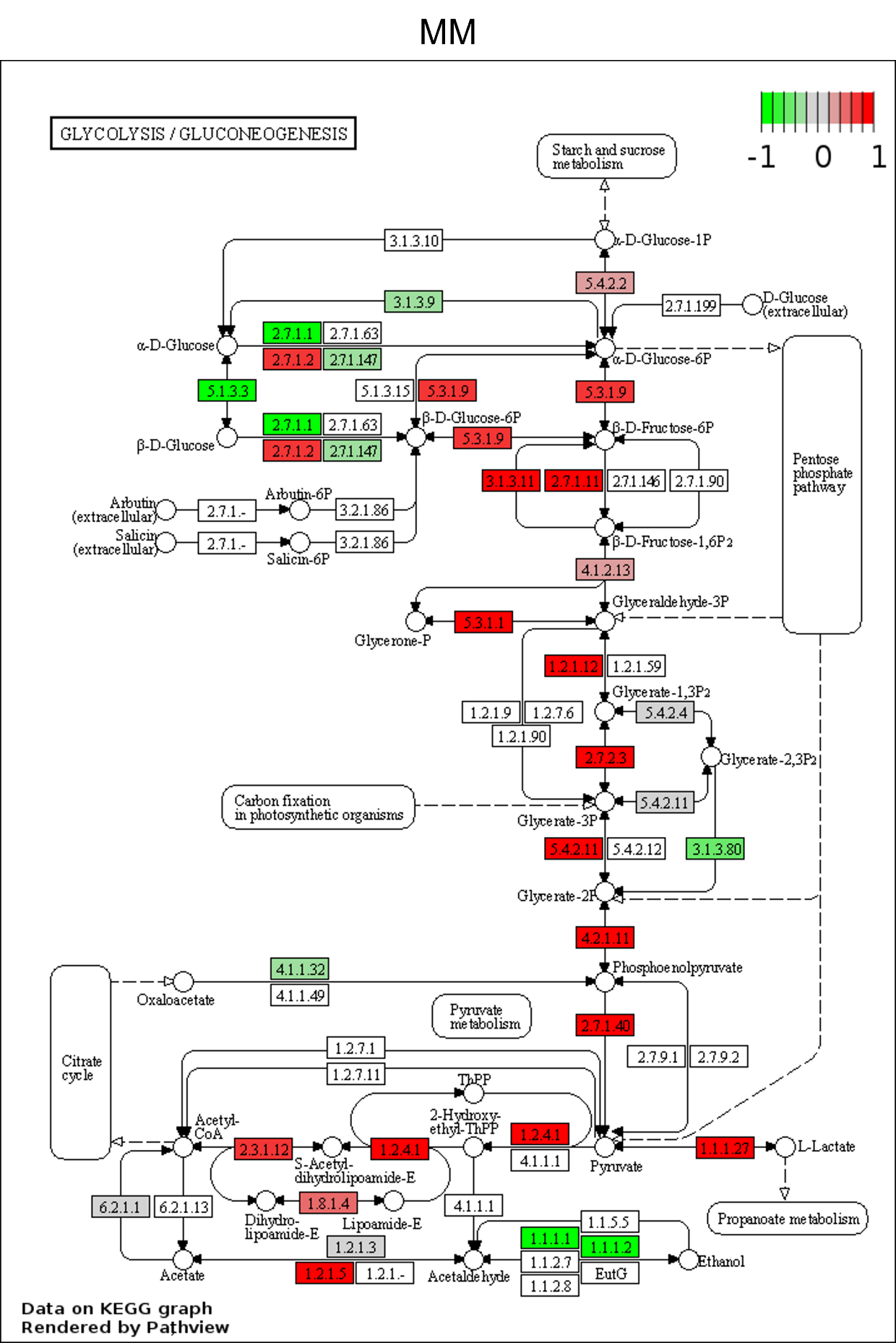

**Supplementary Fig. 23.** KEGG map of glycolysis and gluconeogenesis enzyme pathway rendered by Pathview. Log2-ratio calculated using scRNAseq data from the MM condition.

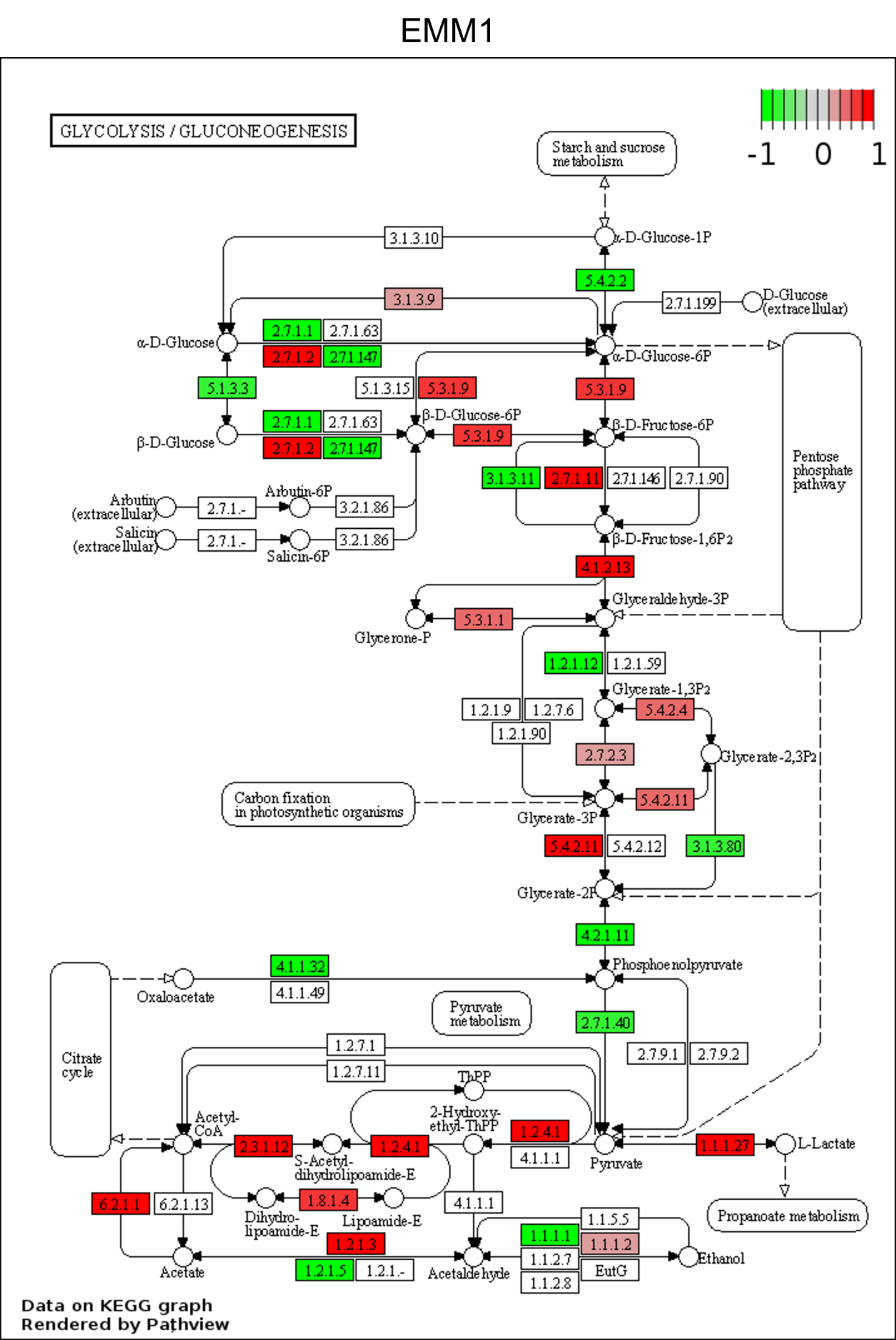

**Supplementary Fig. 24.** KEGG map of glycolysis and gluconeogenesis enzyme pathway rendered by Pathview. Log2-ratio calculated using scRNAseq data from the EMM1 condition.

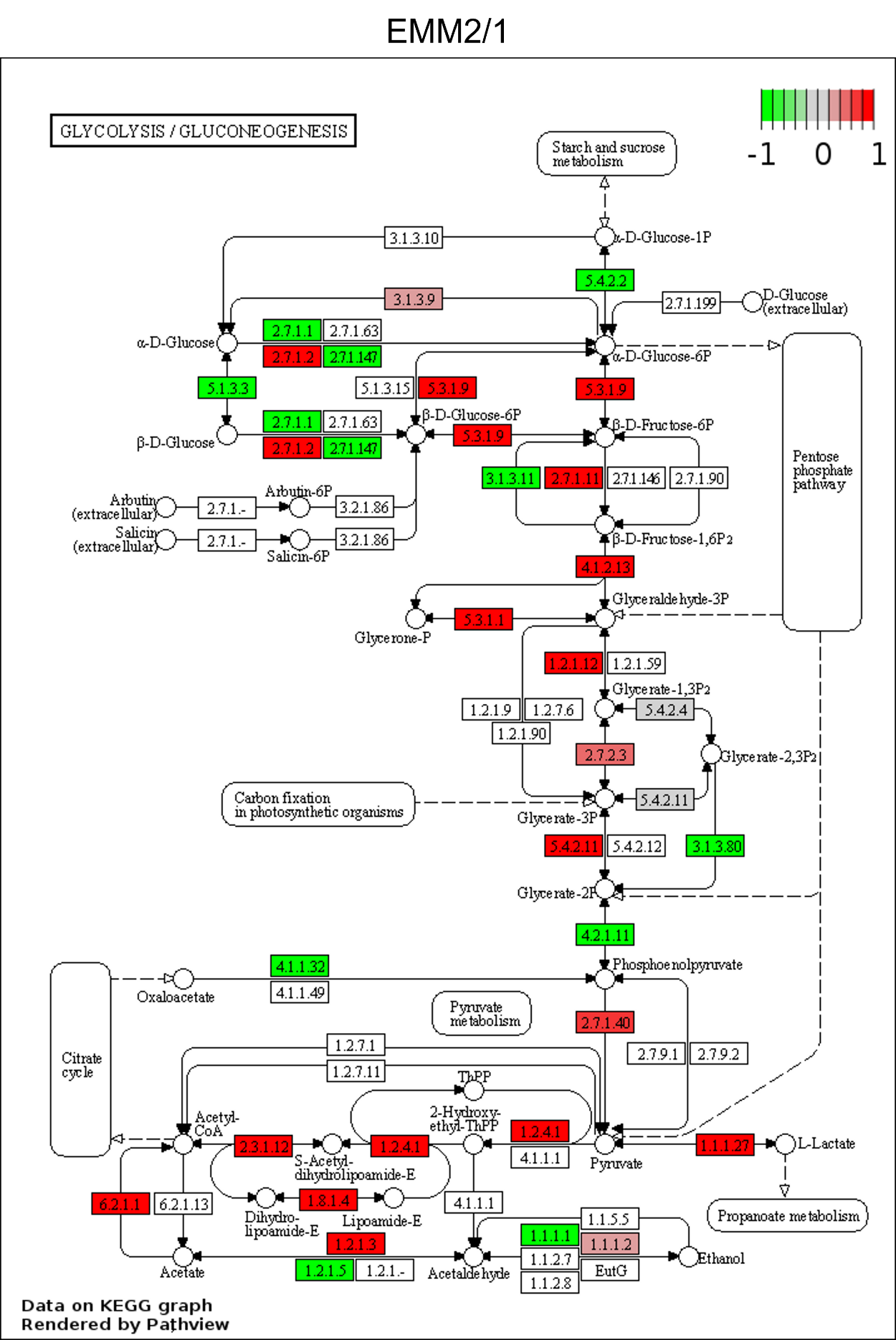

**Supplementary Fig. 25.** KEGG map of glycolysis and gluconeogenesis enzyme pathway rendered by Pathview. Log2-ratio calculated using scRNAseq data from the EMM2/1 condition.

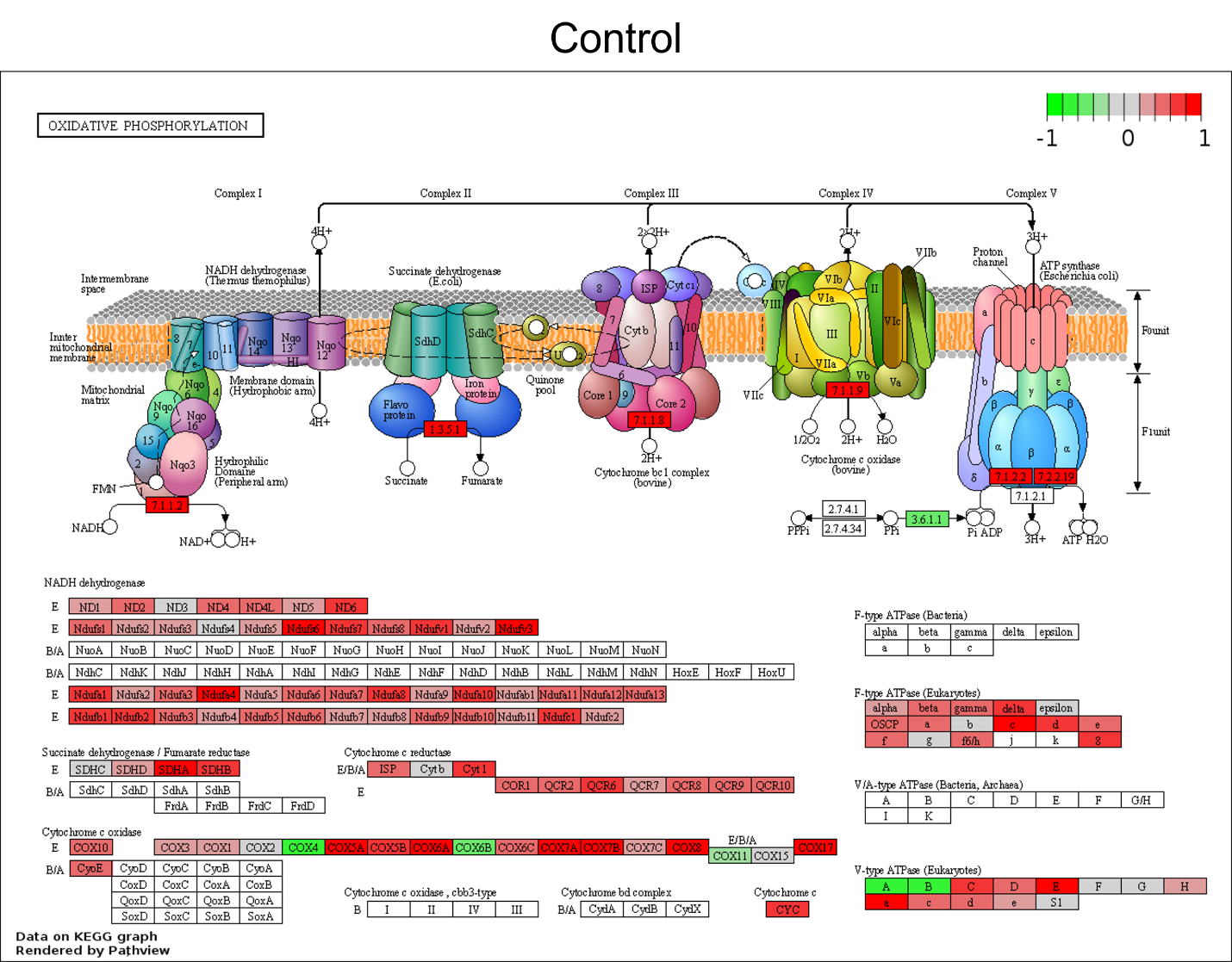

**Supplementary Fig. 26.** KEGG map of oxidative phosphorylation enzyme pathway rendered by Pathview. Log2-ratio calculated using scRNAseq data from the Control condition.

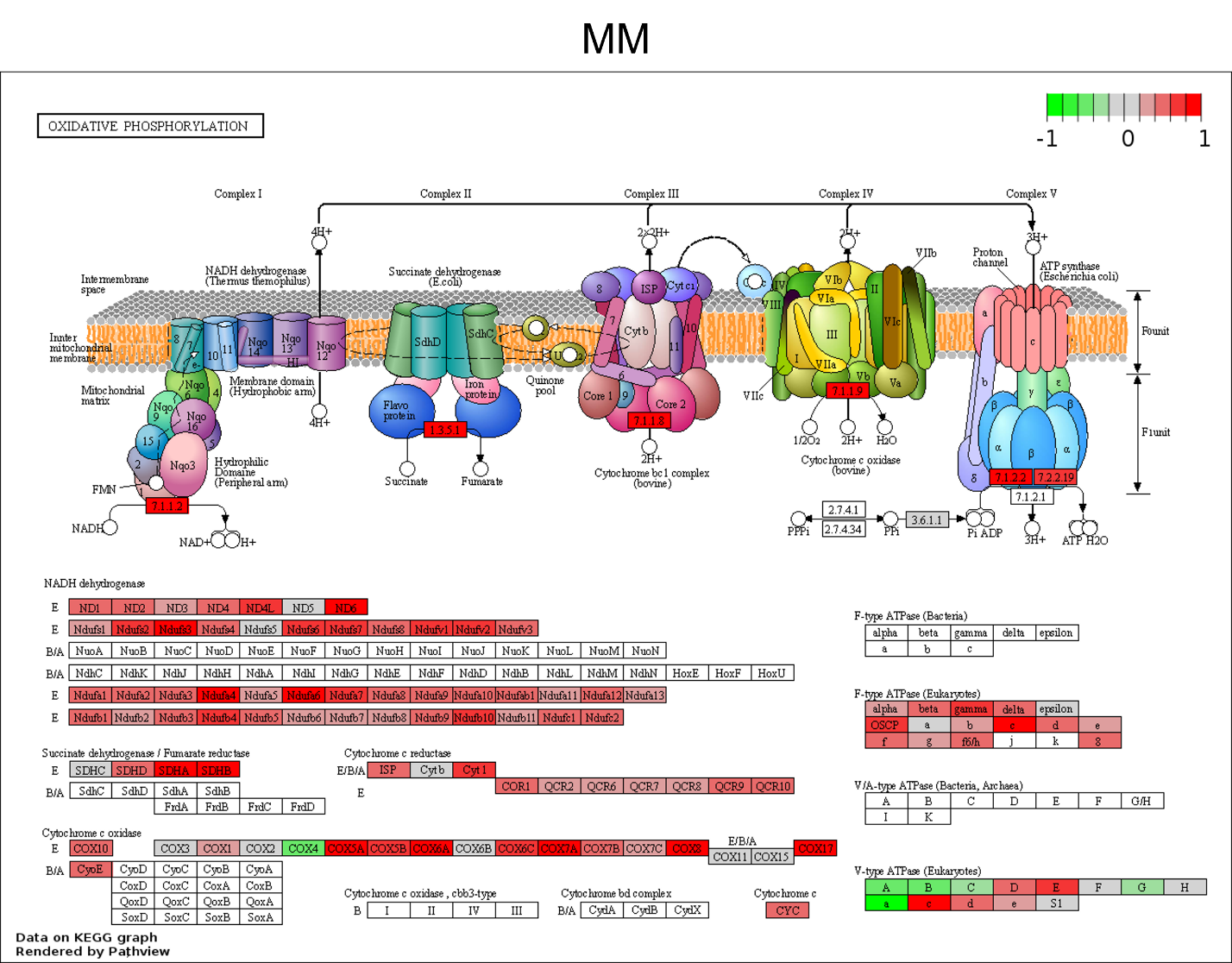

**Supplementary Fig. 27.** KEGG map of oxidative phosphorylation enzyme pathway rendered by Pathview. Log2-ratio calculated using scRNAseq data from the MM condition.

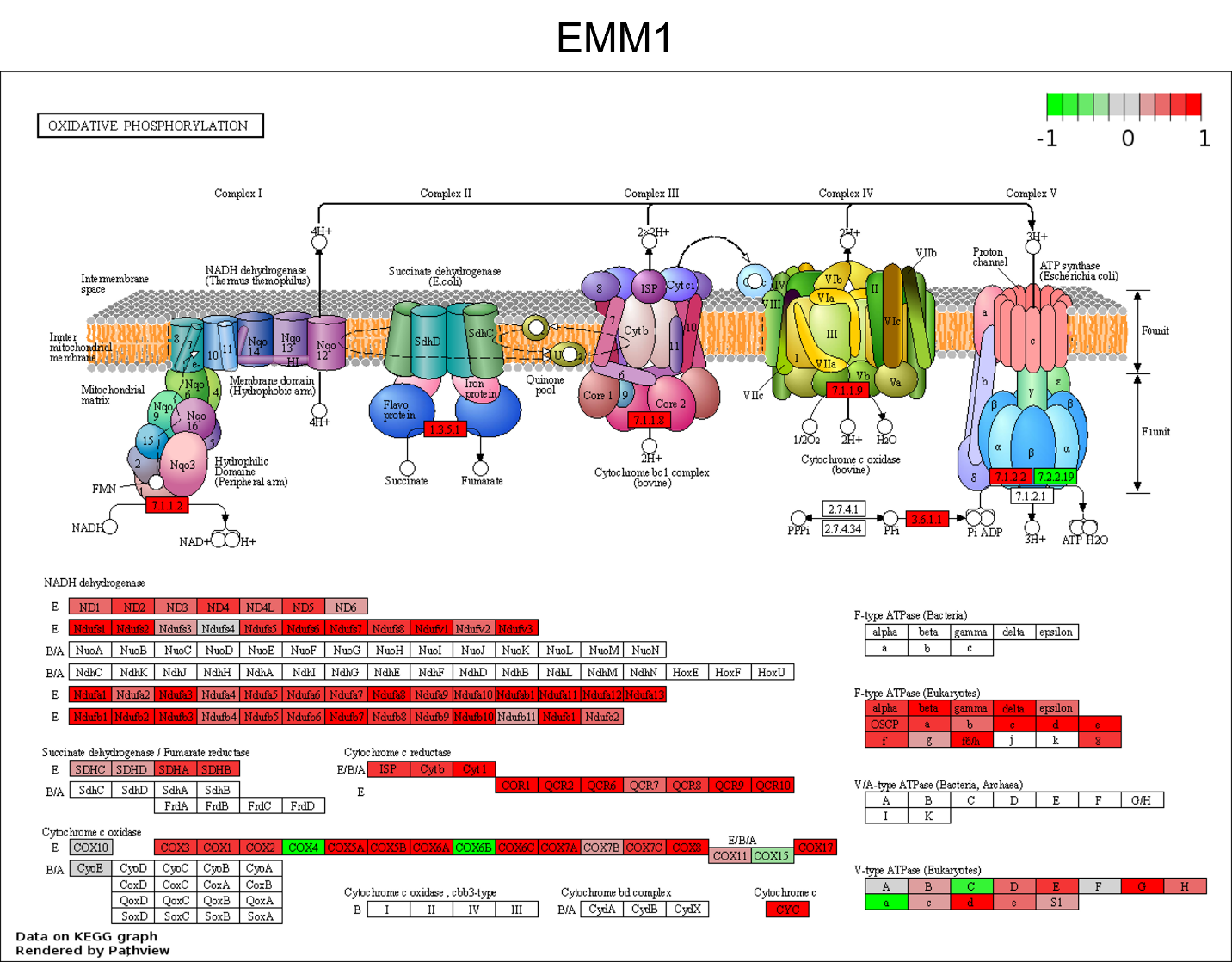

**Supplementary Fig. 28.** KEGG map of oxidative phosphorylation enzyme pathway rendered by Pathview. Log2-ratio calculated using scRNAseq data from the EMM1 condition.

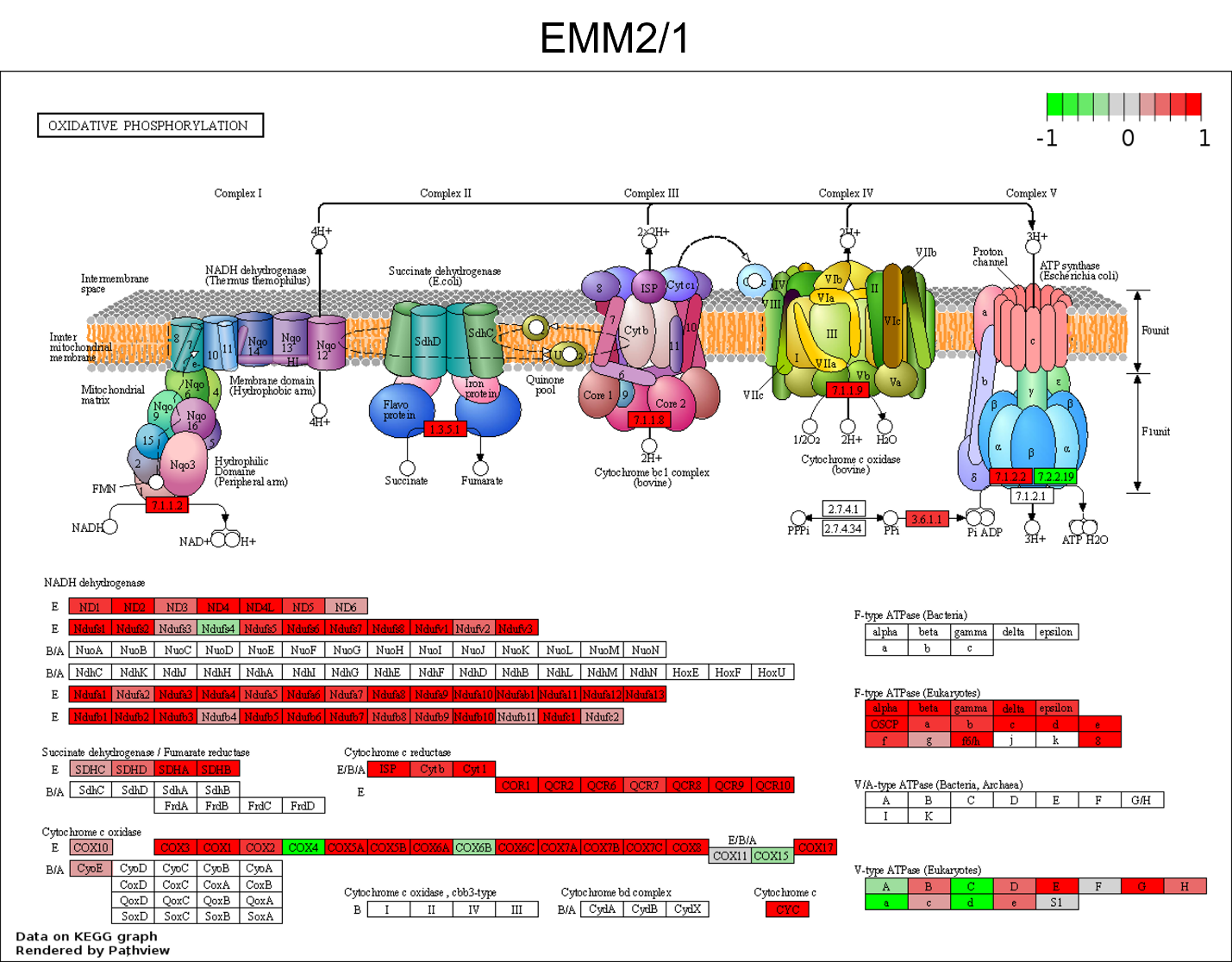

**Supplementary Fig. 29.** KEGG map of oxidative phosphorylation enzyme pathway rendered by Pathview. Log2-ratio calculated using scRNAseq data from the EMM2/1 condition.

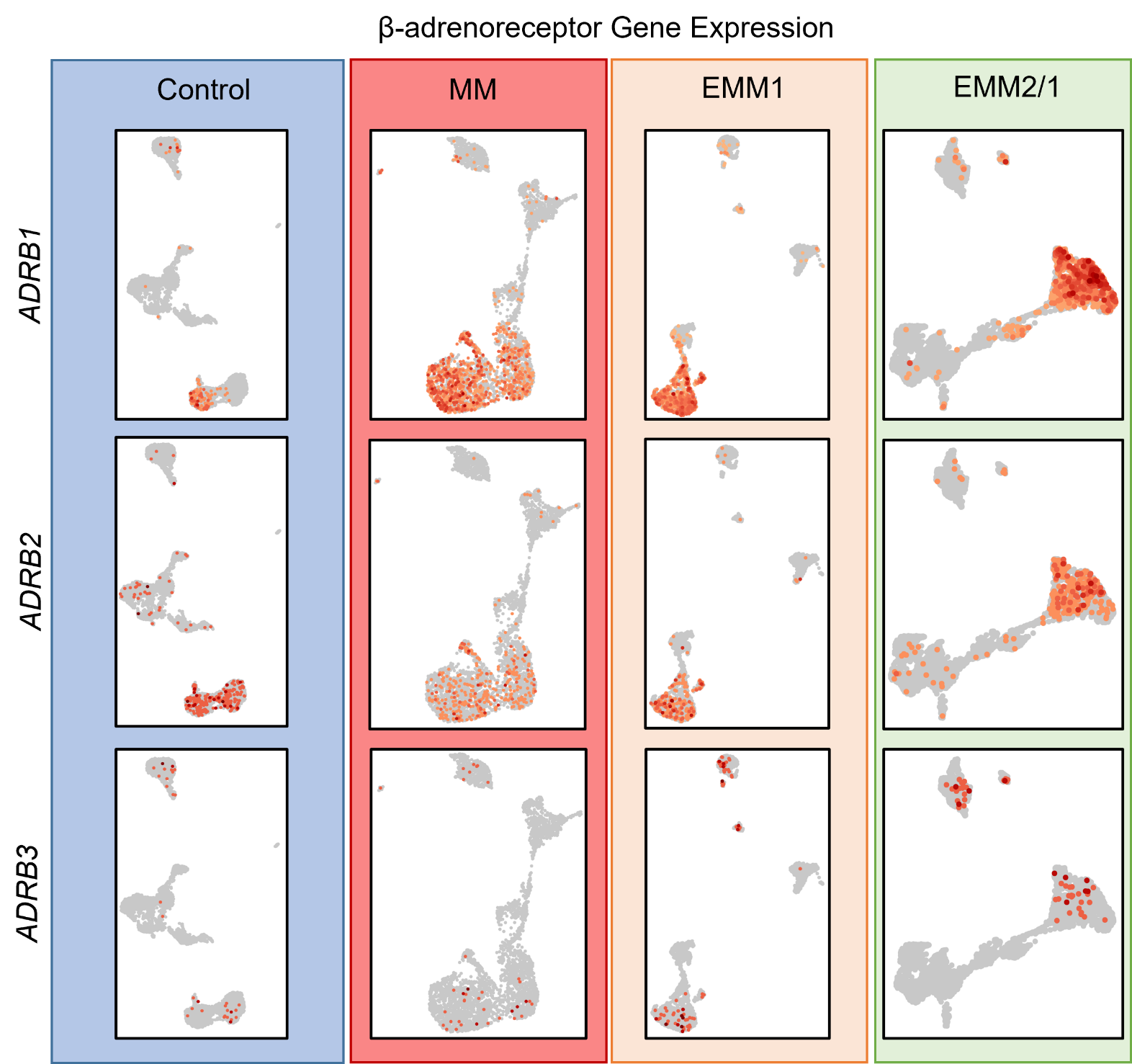

**Supplementary Fig. 30.** Spatial expression of β-adrenoreceptor genes within single cell RNA sequencing datasets for day 34 organoids in each condition. Data presented as UMAP projections of k-means 8 clustering.

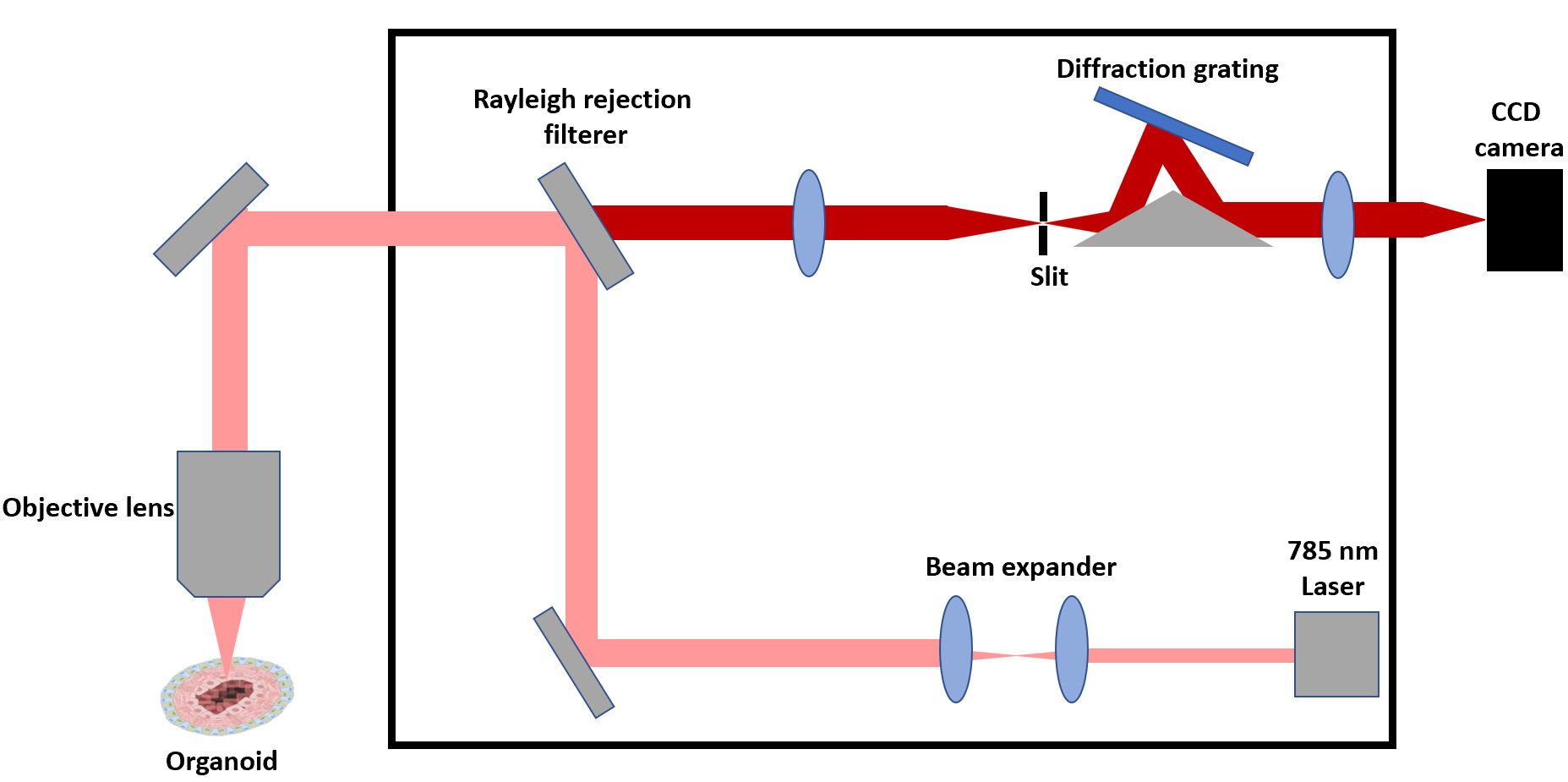

**Supplementary Fig. 31.** Schematic diagram for the Renishaw confocal Raman spectrometer for use with human heart organoids.
